# A Fast, Provably Accurate Approximation Algorithm for Sparse Principal Component Analysis Reveals Human Genetic Variation Across the World

**DOI:** 10.1101/2022.04.21.489052

**Authors:** Agniva Chowdhury, Aritra Bose, Samson Zhou, David P. Woodruff, Petros Drineas

## Abstract

Principal component analysis (PCA) is a widely used dimensionality reduction technique in machine learning and multivariate statistics. To improve the interpretability of PCA, various approaches to obtain sparse principal direction loadings have been proposed, which are termed Sparse Principal Component Analysis (SPCA). In this paper, we present `ThreSPCA`^1^, a provably accurate algorithm based on thresholding the Singular Value Decomposition for the SPCA problem, without imposing any restrictive assumptions on the input covariance matrix. Our thresholding algorithm is conceptually simple; much faster than current state-of-the-art; and performs well in practice. When applied to genotype data from the 1000 Genomes Project, `ThreSPCA` is faster than previous benchmarks, at least as accurate, and leads to a set of interpretable biomarkers, revealing genetic diversity across the world.

## 1 Introduction

Principal Component Analysis (PCA) and the related Singular Value Decomposition (SVD) are fundamental data analysis and dimensionality reduction tools that are used across a wide range of areas including machine learning, multivariate statistics, and many others. These tools return a set of orthogonal vectors of decreasing importance that are often interpreted as fundamental latent factors that underlie the observed data. Even though the vectors returned by PCA and SVD have strong optimality properties, they are notoriously difficult to interpret in terms of the underlying processes generating the data [1], since they are linear combinations of *all* available data points or *all* available features. The concept of Sparse Principal Components Analysis (SPCA) was introduced in the seminal work of [2], where sparsity constraints were enforced on the singular vectors in order to improve interpretability; see for example, document analysis applications in [2,1,3].

Formally, given a positive semidefinite (PSD) matrix 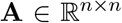, SPCA can be defined as the constrained maximization problem:^2^

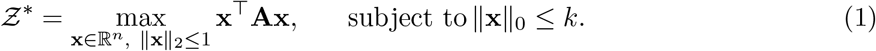

In the above formulation, **A** is a covariance matrix representing, for example, all pairwise feature or object similarities for an underlying data matrix. Therefore, SPCA can be applied to either the object or feature space of the data matrix, while the parameter *k* controls the sparsity of the resulting vector and is part of the input. Let **x**\* denote a vector that achieves the optimal value 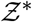 in the above formulation. Intuitively, the optimization problem of eqn. (1) seeks a *sparse*, unit norm vector **x**\* that maximizes the data variance. It is well-known that solving the above optimization problem is `NP`-hard [4] and that its hardness is due to the sparsity constraint. Indeed, if the sparsity constraint were removed, then the resulting optimization problem can be easily solved by computing the top left or right singular vector of **A** and its maximal value 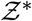 is equal to the top singular value of **A**.

In this work, we explore the potential of SPCA in the analysis of genetics data leveraging a *provably accurate* thresholding algorithm for SPCA. In genetics, PCA is a tool of paramount importance and is ubiquitously used to estimate population structure and extract ancestry information [5]. It is well-known that genome-wide association studies (GWAS) that attempt to identify genetic markers that are associated with complex traits in a typical case/control setting can be grossly confounded by the underlying population structure, due to the presence of subgroups in the population that belong to different ancestries in both the case and control groups [6]. To account for such population stratification and to minimize the underlying spurious associations, researchers typically use the top few principal components as covariates in the underlying model. However, the principal components are linear combinations of all available genetic markers and, therefore, are not interpretable. SPCA is an obvious remedy towards that end, since one can use it to identify Single Nucleotide Polymorphisms (SNPs) or genetic markers carrying information about the underlying genetic ancestry. See also [7,8,9] for prior work motivating and using SPCA in the context of human genetics data analysis.

### 1.1 Our Contributions

Thresholding is a simple algorithmic concept, where each coordinate of, say, a vector is retained if its value is sufficiently high; otherwise, it is set to zero. Thresholding naturally preserves entries that have large magnitude while creating sparsity by eliminating small entries. Therefore, it seems like a logical strategy for SPCA: after computing a dense vector that approximately solves a PCA problem, perhaps with additional constraints, thresholding can be used to sparsify it.

We present a simple, provably accurate, thresholding algorithm (`ThreSPCA`, Section 2.1) for SPCA that leverages the fact that the top singular vector is an optimal solution for the SPCA problem without the sparsity constraint. Our algorithm actually uses a thresholding scheme that leverages the top few singular vectors of the underlying covariance matrix; it is simple and intuitive, yet offers tradeoffs in running time vs. accuracy, the first of its kind. Our algorithm returns a vector that is provably sparse and, when applied to the input covariance matrix **A**, provably captures the optimal solution 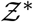 up to a small additive error. Indeed, our output vector has a sparsity that depends on *k* (the target sparsity of the original SPCA problem of eqn. (1)) and *ε* (an accuracy parameter between zero and one). Our analysis provides unconditional guarantees for the accuracy of the solution of the proposed thresholding scheme. To the best of our knowledge, no such analyses have appeared in prior work (see Section 1.2 for details). We emphasize that our approach only requires an approximate SVD and, as a result, `ThreSPCA` runs very quickly. In practice, `ThreSPCA` is much faster than current state-of-the-art and at least as accurate in the analysis of human genetics datasets. An additional contribution of our work is that, unlike prior work, our algorithm has a clear trade-off between quality of approximation and output sparsity. Indeed, by increasing the density of the final SPCA vector, one can improve the amount of variance that is captured by our SPCA output. See Theorem 2.1 for details on this sparsity vs. accuracy trade-off for `ThreSPCA`.

Importantly, we evaluate `ThreSPCA` on the genotype dataset from 1000 Genomes (1KG) Project [10] and on simulated genotype data in order to practically assess its performance. `ThreSPCA` identifies functionally relevant, interpretable SNPs from the 1KG data and, from an accuracy perspective, it performs comparably to current state-of-the-art SPCA algorithms while being much faster than its competitors.

### 1.2 Prior work

SPCA was formally introduced by [2]; however, previously studied PCA approaches based on rotating [11] or thresholding [12] the top singular vector of the input matrix seemed to work well, at least in practice, given sparsity constraints. Following [2], there has been an abundance of interest in SPCA, with extensions based on LASSO (ScoTLASS) on an *ℓ*_1_ relaxation of the problem [13] or a non-convex regression-type approximation, penalized similar to LASSO [14]. Additional heuristics based on LASSO [15] and non-convex *ℓ*_1_ regularization [14,16,17,18] have also been explored. Other approaches such as random sampling based on non-convex *ℓ*_1_ relaxations [19], branch-and-bound heuristic motivated by greedy spectral ideas [20] have also been studied. Spectral approaches based on iterative methods such as the power method have been extensively explored [21,3,22,23] including a SPCA algorithm with early stopping for the power method, based on the target sparsity [23]. Another line of work focused on using semidefinite programming (SDP) relaxations of SPCA [2,24,25,26]. Despite the variety of heuristic-based SPCA approaches, very few theoretical guarantees have been provided; this is partially explained by a line of hardness-of-approximation results. The sparse PCA problem is well-known to be `NP-Hard` [4]. [27] shows that if the input matrix is not PSD, then even the *sign* of the optimal value cannot be determined in polynomial time unless `P = NP`, ruling out any multiplicative approximation algorithm. In the case where the input matrix is PSD, [28] shows that it is NP-hard to approximate the optimal value up to multiplicative (1 + *ε*) error, ruling out any polynomial-time approximation scheme (PTAS).

Prior work that offers provable guarantees, typically given *some assumptions about the input matrix*, includes [3], which analyzed a specific set of vectors in a low-dimensional eigenspace of the input matrix and presented relative error guarantees for the optimal objective, given the assumption that the input covariance matrix has a decaying spectrum. The time complexity of the algorithm of [3] is given by 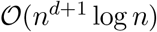 (due to solving an exact SVD), where *d* is the low rank parameter that affects the accuracy of the output. Even for *d* = 1, the theoretical time complexity boils down to 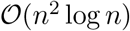 and it is not clear how to make use of an approximate SVD algorithm to improve this running time without affecting its theoretical bound. Furthermore, for a high precision output, one generally needs d to be larger than one, in which case the practical running time also increases drastically. [29] gave a polynomial-time algorithm that solves sparse PCA *exactly* for input matrices of constant rank. [28] showed that sparse PCA can be approximated in polynomial time within a factor of *n*^−1/3^ and also highlighted an additive PTAS of [30] based on the idea of finding multiple disjoint components and solving bipartite maximum weight matching problems. This PTAS needs time *n*^poly(1/*ε*)^, whereas `ThreSPCA` has running time that depends on the sparsity of the input data.

SPCA has been applied in the context of human genetics before, in the form of sparse factor analysis (SFA) [8] and with a penalty term in LASSO (`L-PCA`) or Adaptive LASSO (`AL-PCA`) [7]. However, there are a number of aspects that our work improves compared to prior studies. First, unlike `ThreSPCA`, the SFA method used some prior assumptions on the genotype matrix and none of these previous studies come with a theoretical guarantee showing a clear sparsity vs. accuracy trade-off. Second, prior work has to tune the penalty parameter in [7] several times in order to achieve a specific sparsity value in practice, which increases the running time of the method. Third, the convergence of the SPCA algorithm proposed by [7] depends on an initial PC score, which typically relies on the top right singular vector of the data and necessitates the computation of an exact SVD, which is expensive. It is not clear whether replacing the exact SVD with a fast approximate SVD would affect the results of [7].

## 2 Materials and Methods

### 2.1 The ThreSPCA algorithm

#### Notation

We use bold letters to denote matrices and vectors. For a matrix 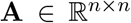, we denote its (*i,j*)-th entry by *A_i,j_*; its *i*-th row by **A**_*i*\*_, and its *j*-th column by **A**_\**j*_; its 2-norm by 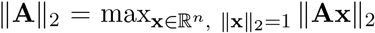; and its (squared) Frobenius norm by 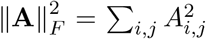. We use the notation **A** ≻ 0 to denote that the matrix **A** is symmetric positive semidefinite (PSD) and 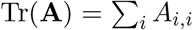 to denote its trace, which is also equal to the sum of its singular values. Given a PSD matrix 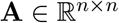, its Singular Value Decomposition is given by **A** = **UΣU**^*T*^, where **U** is the matrix of left/right singular vectors and **Σ** is the diagonal matrix of singular values.

#### Our approach: SPCA via SVD Thresholding

To achieve nearly input sparsity runtime, our thresholding algorithm is based upon using the top *ℓ* right (or left) singular vectors of the PSD matrix **A**. Given **A** and an accuracy parameter *ε*, our approach first computes 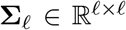 (the diagonal matrix of the top *ℓ* singular values of **A**) and 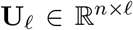 (the matrix of the top *ℓ* left singular vectors of **A**), for *ℓ* = 1/*ε*. Then, it *deterministically* selects a subset of 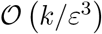 rows of **U**_*ℓ*_ using a simple thresholding scheme based on their squared row norms (recall that *k* is the sparsity parameter of the SPCA problem). In the last step, it returns the *top right singular vector* of the matrix consisting of the columns of 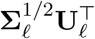 that correspond to the row indices of **U**_*ℓ*_ chosen in the thresholding step. Notice that this right singular vector is an 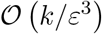-dimensional vector, which is finally expanded to a vector in 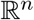 by appropriate padding with zeros. This sparse vector is our approximate solution to the SPCA problem of eqn. (1).

This simple algorithm is somewhat reminiscent of prior thresholding approaches for SPCA. However, to the best of our knowledge, no provable a priori bounds were known for such algorithms without strong assumptions on the input matrix. This might be due to the fact that prior approaches focused on thresholding only the top right singular vector of **A**, whereas our approach thresholds the top *ℓ* = 1/*ε* right singular vectors of **A**. This slight relaxation allows us to present provable bounds for the proposed algorithm.

In more detail, let the SVD of **A** be **A** = **UΣU**^*T*^. Let 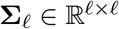 be the diagonal matrix of the top *ℓ* singular values and let 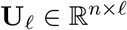 be the matrix of the top *ℓ* right (or left) singular vectors. Let *R* = {*i*_1_,…, *i*_|*R*|_} be the set of indices of rows of **U**_*ℓ*_ that have squared norm at least *ε*^2^/*k* and let 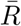 be its complement. Here |*R*| denotes the cardinality of the set *R* and 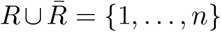. Let 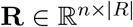 be a sampling matrix that selects^3^ the rows of **U**_*ℓ*_ whose indices are in the set *R*. Given this notation, we are now ready to state Algorithm 1. Notice that **Ry** satisfies ||**Ry**||_2_ = ||**y**||_2_ = 1 (since **R** has orthogonal columns) and **||Ry||**_0_ =|*R*|. Since *R* is the set of rows of **U**_*ℓ*_ with squared norm at least *ε*^2^/*k* and 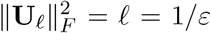, it follows that |*R*| ≤*k*/*ε*^3^. Thus, the vector returned by Algorithm 1 has *k*/*ε*^3^ sparsity and unit norm. (See the Appendix for more details.)

#### Algorithm 1: ThreSPCA: fast thresholding SPCA via SVD

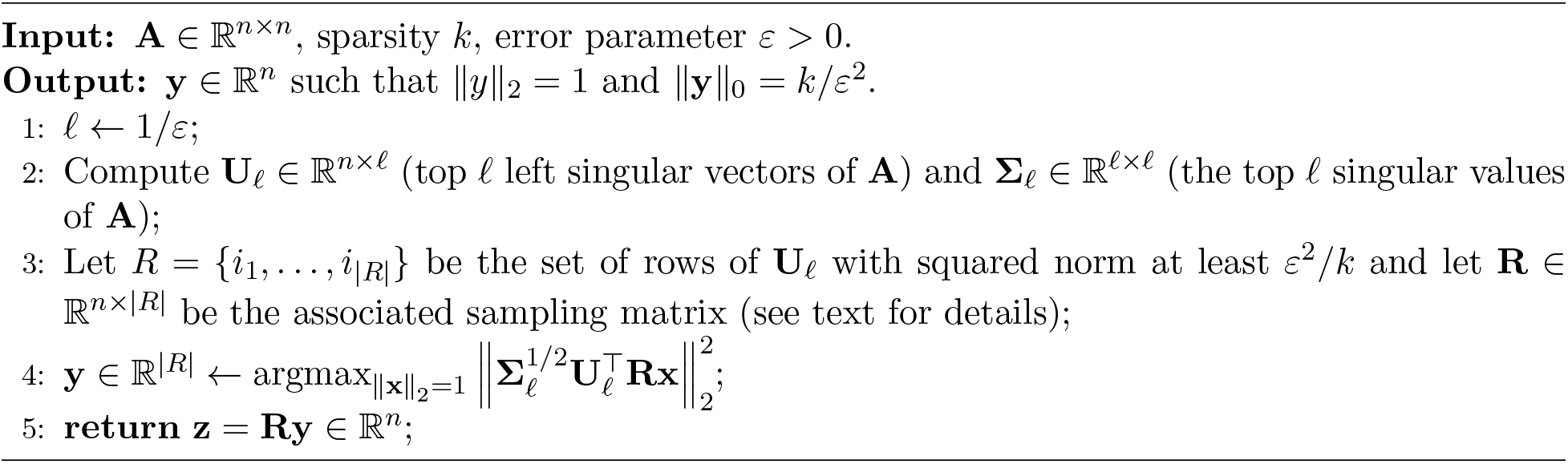

#### Theorem 2.1

*Let k be the sparsity parameter and ε* ∈ (0,1] *be the accuracy parameter. Then, the vector* 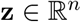 (*the output of Algorithm 1) has sparsity k/ε*^3^, *unit norm, and satisfies*

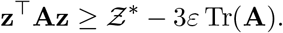

The optimality gap of Theorem 2.1 depends on Tr(**A**), which is the sum of the eigenvalues of **A** and can also be viewed as the total variance of the data. Therefore, if we divide both sides of the bound in Theorem 2.1 by Tr(**A**), the resulting bound is given by 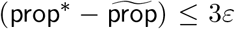, where for a given *k*, 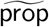 is the proportion of the total variance explained by the output of `ThreSPCA` and `prop`* is the proportion of the total variance explained by the optimal Sparse PC. Now, trivially, we have 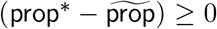, since `prop`* is the maximum variance explained by Sparse PC for a given sparsity value. Thus, combining these two yields 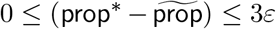, which can be interpreted as the quality-of-approximation in terms of the proportion of total variance explained by `ThreSPCA`.

The proof of Theorem 2.1 is deferred to the appendix. See Section A for the proof of Theorem 2.1 as well as an intermediate result (Lemma A.1) that leads to the final bound in Theorem 2.1. The running time of Algorithm 1 is dominated by the computation of the top *ℓ* singular vectors and singular values of the matrix **A**. One could always use the SVD of the full matrix **A** (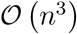 time) to compute the top *ℓ* singular vectors and singular values of **A**. In practice, any iterative method, such as subspace iteration using a random initial subspace or the Krylov subspace of the matrix, can be used towards this end. We now address the inevitable approximation error incurred by such approximate SVD methods below.

#### Using approximate SVD algorithms

Although the guarantees of Theorem 2.1 in Algorithm 1 use an exact SVD computation, which could take time 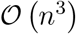, we can further improve the running time by using an approximate SVD algorithm such as the randomized block Krylov method of [31], which runs in nearly input sparsity running time. Our analysis uses the relationships 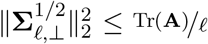 and *σ*_1_(Σ_*ℓ*_) ≤Tr(**A**). The randomized block Krylov method of [31] recovers these guarantees up to a multiplicative (1 + *ε*) factor, in 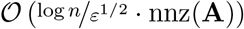 time. Here nnz(**A**) denotes the number of non-zero entries of the matrix **A**, which is 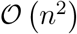 for dense matrices.

#### Extracting additional sparse PCs

To get multiple sparse PCs using Algorithm 1, we remove the top principal component from the data and run `ThreSPCA` on the residual dataset. In other words, let 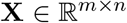 be the *mean-centered* data matrix corresponding to **A**, *i.e*., **A** =**X**^T^**X**. Let 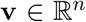 be the top right singular vector of **X**; then, in order to get the second sparse PC, we run `ThreSPCA` on the covariance matrix 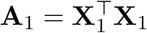, where **X**_1_ = **X**– **Xvv**^T^.

### 2.2 Data

#### 1000 Genome Data

In order to evaluate the speed and accuracy of `ThreSPCA` as well as to interpret its output, we first analyzed data from the 1000 Genome Project (1KG) [10], which contained genotype data from 2, 503 individuals with 39,517,397 SNPs sampled from 26 different populations across all continents. After performing Quality Control (QC) with minor allele frequency below 5% and, subsequently, pruning related genotypes for Linkage Disequilibrium (LD) using a window size of 50 kb and *r*^2^ >0.2, we finally retained 360,498 variants.

#### Simulated Data

We generated simulated data emulating real-world populations to evaluate whether `ThreSPCA` can correctly identify markers which contribute to the genetic differences between and within the populations. Based on previous work [32], we simulated two datasets varying *m* = {5000,10000} SNPs genotyped across *n* = {500,1000} individuals based on the Pritchard- Stephens-Donelly (PSD) model [33] with the mixing parameter between populations, *α* = 0.01. The allele frequencies were simulated based on real-world data from three divergent populations, namely CEU (Utah residents with Northern and Western European ancestry), ASW (African ancestry in Southwestern US), and MXL (Mexican ancestry in California) from the HapMap Phase 3 data [34]. We selected a threshold *t* and varied it across the range *t* ={100,250, 500}, representing the number of SNPs which contribute to population structure between the populations (true positives); the remaining *m* – *t* genotypes were simulated such as they had minimal genetic differences between populations (false positives). We simulated 200 data sets (100 each for values of *m* and *n*) and applied `ThreSPCA`, `L-PCA` and `AL-PCA` for comparative analyses to evaluate their efficacy.

### 2.3 Experiments

We performed QC on the 1KG data, including LD pruning using PLINK2.0 [35]. PCA was performed using TeraPCA [36]. Annotation of ThreSPCA derived variants were performed in Ensembl Variant Effect Predictor (VEP) [37]. We performed Gene Ontology (GO) pathway analyses using `clusterProfiler` [38] in R. We ran `ThreSPCA`, with the threshold parameter *ℓ*, fixed to one.

## 3 Results

### 3.1 ThreSPCA reveals genetic diversity across the world

We applied ThreSPCA with a sparsity threshold of *k*=500 on the 1KG data after quality control and pruning for correlated SNPs. We obtained sets of informative markers of cardinality *k* from each of the PCs. We restricted our analysis to the top three PCs, resulting in a total of 1,500 SNPs, which explained approximately 83% of the variance. Thus, we performed PCA on a reduced 1KG data with 2,503 individuals and 1,500 SNPs. We observed that both the PCA plot and the allele frequency bar plot, grouped by populations across the world, are almost identical. The squared Pearson correlation coefficient (*r*^2^) between the top two PCs from the original 1KG data and ThreSPCA informed variants are very high, equal to 0.98, 0.97 and 0.94 for PCs 1, 2 and 3 respectively. Thus, the PCA plot of the informative markers clearly preserves the clusters of each subgroup (Figure 1) and reveals fine-scale population structure among the groups.

**Fig. 1:**
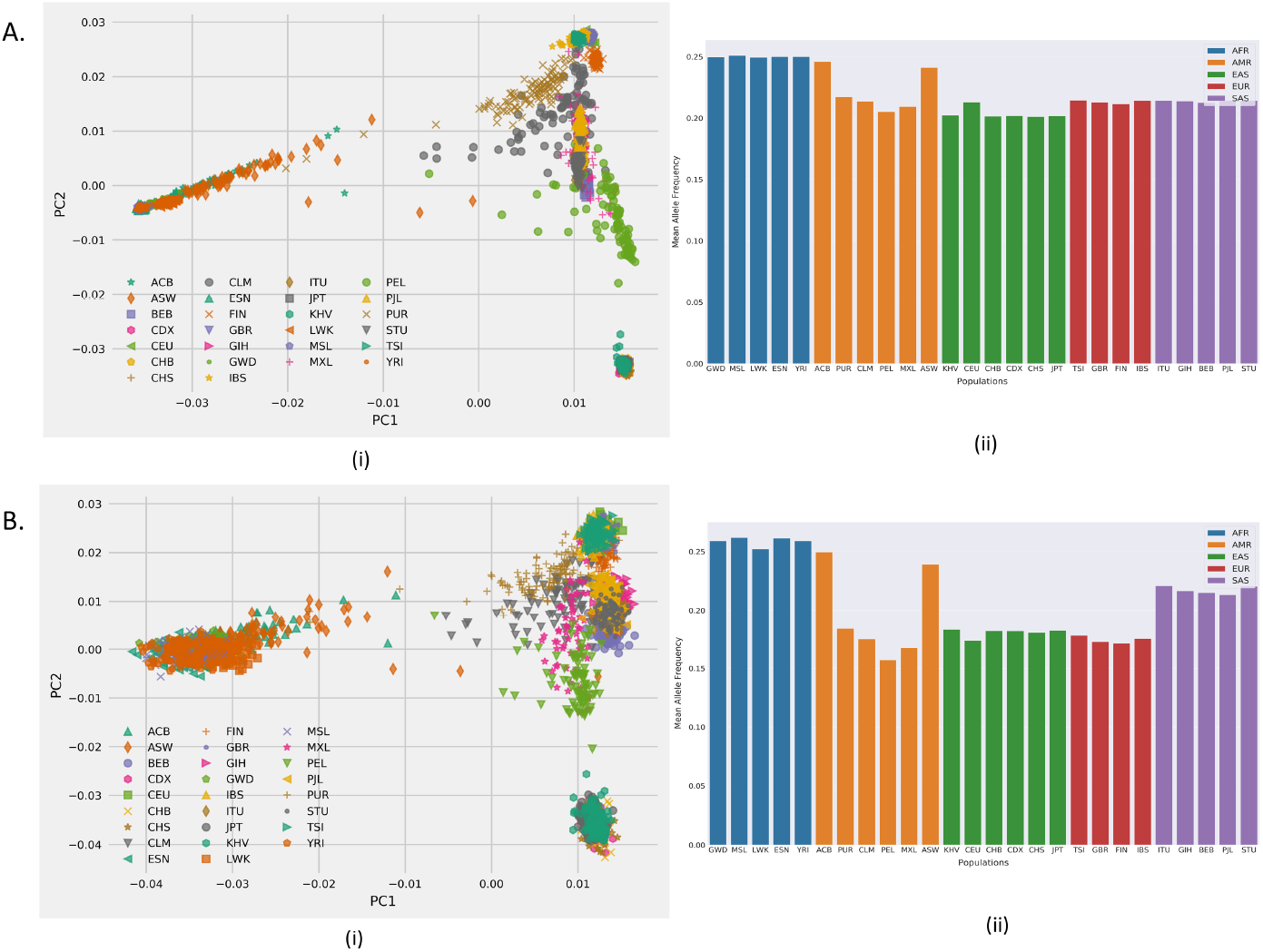
Population structure of world populations from: A. pruned 1KG data with 360,498 SNPs, and B. 1KG data with 1,500 `ThreSPCA` derived variants corresponding to the top three PCs, captured by (i) PCA plot and (ii) mean allele frequency bar plots colored by continental populations arranged in order from Africa (AFR), Americas (AMR), East Asia (EAS), Europe (EUR) and South Asia (SAS).

Examining each of the three PCs closely shows that the mean allele frequency distribution from PC1 is skewed towards the African populations (Appendix Figure 2) and also from the mixed ancestry populations of ASW (Africans in Southwestern US) and ACB (African Caribbeans from Barbados). SNPs obtained from PC2 were almost equally distributed across the continental populations with a slightly higher frequencies in East Asians. PC3 shows a skewness towards South Asian populations (Appendix Figures 3 and 4). To make an informed choice of the sparsity threshold *k*, we computed the *PC scores* from the top two PCs by projecting the sparse vectors obtained from ThreSPCA on the original pruned 1KG data for a range of values of *k* (Appendix Figure 11). We computed *r*^2^ between the *PC scores* obtained from each PC for each value of k and the original PC obtained from the pruned 1KG data. We observe high correlation values for the top two PCs, cumulatively reaching their peak when the sparsity parameter *k* is set to approximately 500 (Appendix Figure 6).

### 3.2 Interpretability of ThreSPCA informed variants

#### Annotating the selected variants

To understand whether the variants derived from `ThreSPCA` for each PC are functionally relevant and biologically interpretable, we annotated them using VEP [37]. We also explored whether these variants were mapping to a trait or disease in the GWAS catalog [39]. Most of the variants were introns with some intergenic and small number of Transcription Factor binding sites, upstream and downstream gene variants, etc. Interestingly, among the coding consequences, 58 variants were missense and likely disease causing. SIFT [40] and PolyPhen [41] statistics revealed that there are seven variants which are deleterious and nine *probably* or *possibly* damaging variants (Figure 2). We also performed GO pathway analyses on `ThreSPCA` informed variants and found significantly enriched pathways common to humans across the world, such as pathways related to synapses and potassium, cation and ion channels, transporter complex, among others (Appendix Figure 7a). We found the calcium signaling pathway from KEGG (Kyoto Encyclopedia of Genes and Genomes) to be significantly enriched (Appendix Figure 7b).

**Fig. 2:**
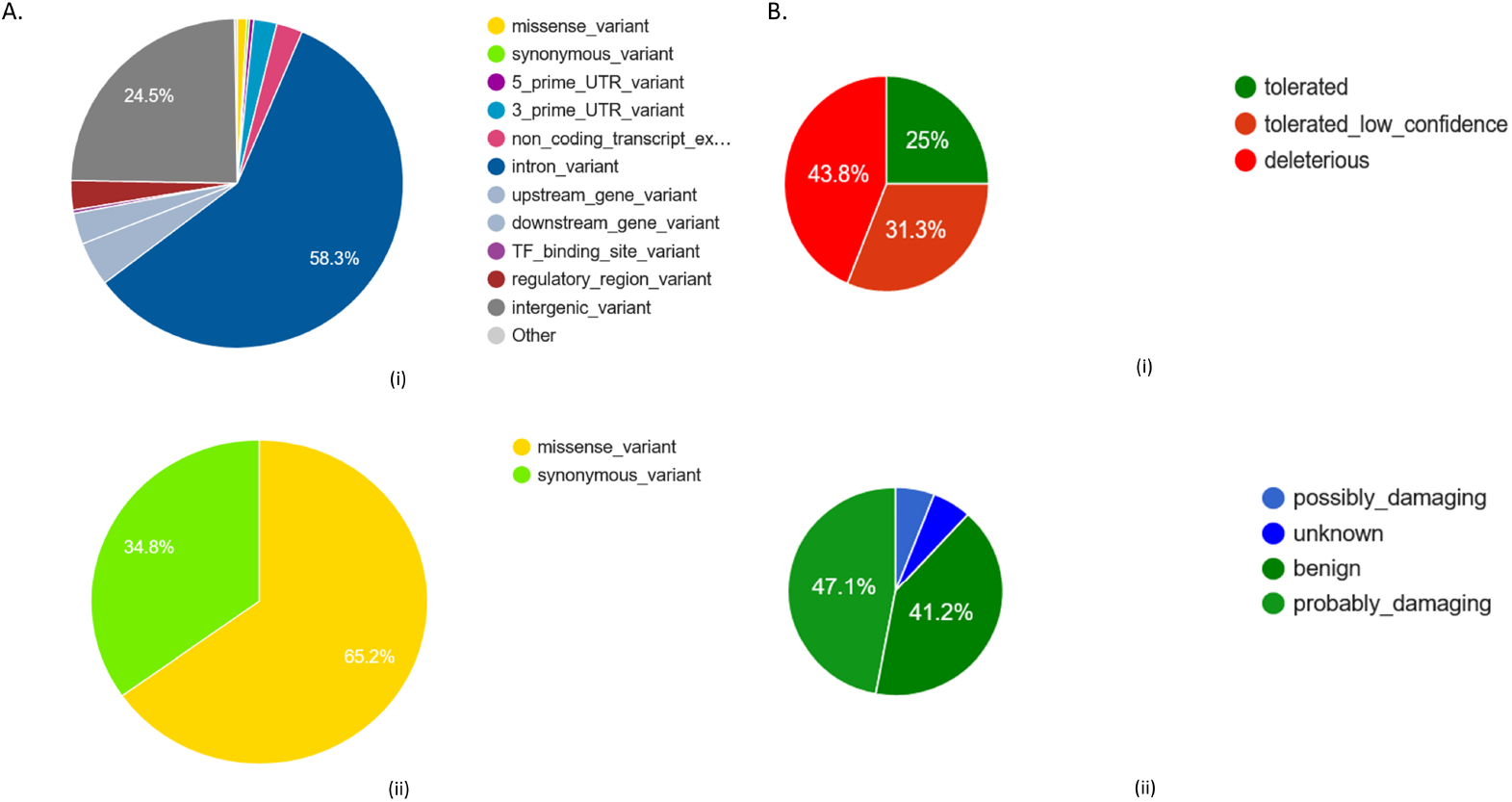
Pie charts showing the percentage of variants from A. (i) most severe consequences and (ii) coding consequences obtained from VEP. B. Deleterious and probably damaging from (i) SIFT and (ii) PolyPhen.

#### Mapping the selected variants to traits

Mapping these variants in GWAS catalog, we found that variants from PC1 mapped to skin pigment measurement (Appendix Table 1), justifying our observation from the PCA plot and mean allele frequency distribution (Figure 2).This is concordant with our observation that ThreSPCA observed variants from PC1 were skewed towards populations of African ancestry (Figure 2), who are darker skinned than the rest of the world. PC2 and PC3 on the other hand mapped to various traits which are commonly found to be varying in populations across the world such as body height, BMI, hip and waist circumference, circadian rhythm, gut microbiome, smoking status, cardiovascular diseases (coronary artery disease, myocardial infarction, etc.), calcium channel blocker use (concordant with calcium signaling pathway found in GO analyses), blood measurements (platelet count, hemoglobin, leukocyte count, etc.), among others.

### 3.3 Comparing ThreSPCA to state-of-the-art

#### Simulation studies

We designed a simulation study to evaluate the correctness of `ThreSPCA` and compared with the state-of-the-art SPCA methods in genetics, namely, `L-PCA` and `AL-PCA` from [7]. The population structure of the simulation shows three distinct clusters for each population with signs of admixture between them (Appendix Figure 1). Applying `ThreSPCA` on the simulated dataset with 10,000 markers and 1,000 individuals, we observed that `ThreSPCA` identified similar numbers (mean) of true positives, i.e., markers contributing to the genetic diversity between and within the populations when compared to its counterparts `L-PCA` and `AL-PCA`, while identifying a significantly smaller number of false positives, i.e., noisy markers which have no difference in allele frequencies between populations (Figure 3b).

**Fig. 3:**
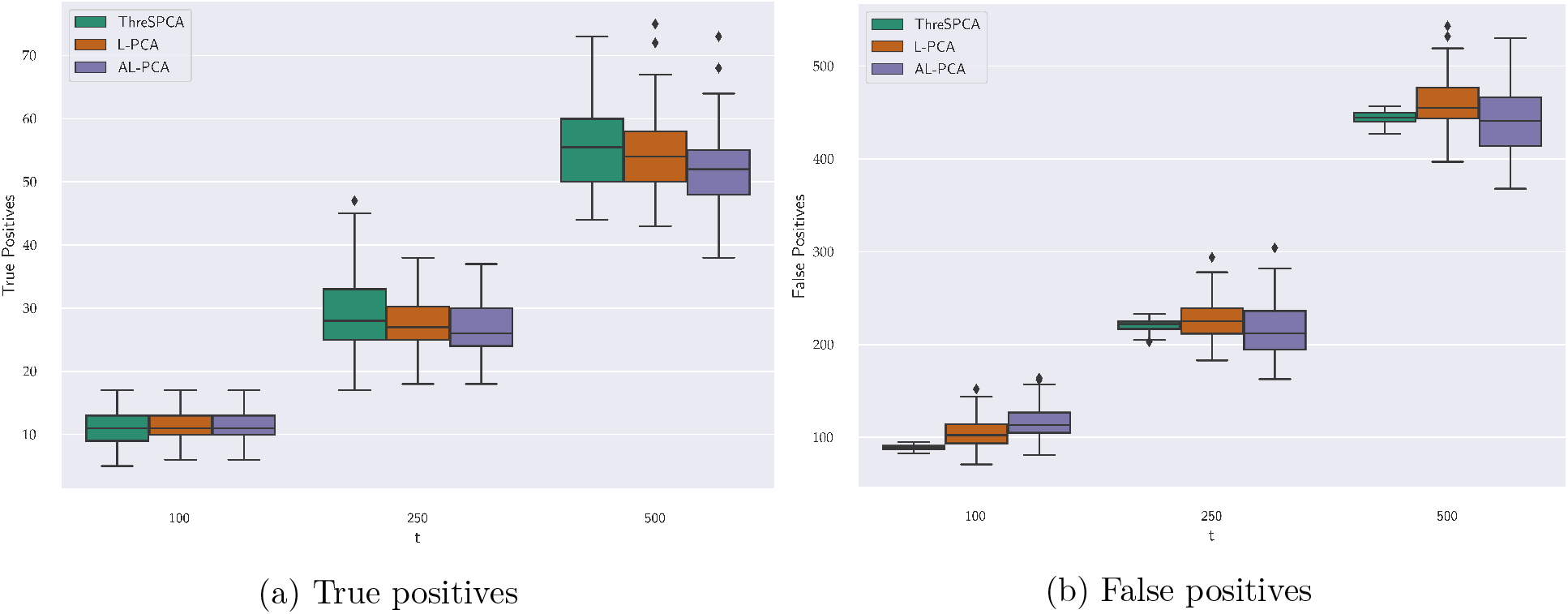
Box and whisker plots comparing between `ThreSPCA`, `L-PCA` and `AL-PCA` for true and false positives obtained from the simulated dataset of *m* = 10, 000 and *n* = 1, 000 and varying values of *t*, i.e., the number of SNPs which contribute to population structure.

#### Real Data

We apply both `ThreSPCA` and `AL-PCA`^4^ on the 1KG data with *k* = 500 and compare the *PC1 scores vs. PC2 scores* generated from the outputs of the aforementioned methods. `ThreSPCA` (Appendix Figure 10a) and `AL-PCA` are almost identical to the corresponding standard PC plot (Appendix Figures 10b and 11a), clearly preserving the clusters of each subgroup. We also plot the *PC scores* computed from `ThreSPCA` and `AL-PCA` against the traditional *PC scores* (Appendix Figure 9) and observed a near-linear relationship between the two SPCA algorithms for both PCs with *r*^2^ = 0.9808 and 0.9426 for PC1 & PC2, respectively. This validates that `ThreSPCA` and AL-PCA are qualitatively very similar to each other in inferring genetic structure.

#### Running Time

`ThreSPCA` clearly outperforms `AL-PCA`. In particular, for any given *k*, while `ThreSPCA` takes less than two minutes in 1KG data, `AL-PCA` takes about 15 minutes to do the same for a given penalty parameter λ> 0, since it needs a full SVD. Moreover, as already mentioned in Section 1.2, λ is a hyper-parameter which needs to be tuned with many cross-nested runs of the data in order to achieve a desired sparsity value. In our case, for the sparsity parameter set to 500, it took at least six runs for each PC. Therefore, the resulting speed-up achieved by `ThreSPCA` is more than 45x for real data set and around 80x for simulated data.

## 4 Discussion

We present `ThreSPCA`, a simple and intuitive approximation algorithm for SPCA, based on a deterministic thresholding scheme, without imposing any restrictive assumption on the input covariance matrix. `ThreSPCA` comes with a provable accuracy guarantee and provides a clear sparsity vs. accuracy trade-off. In practice, it is much faster than the other *state-of-the-art* SPCA methods and indeed, can be implemented in nearly input sparsity time.

Applying `ThreSPCA` on the 1KG data, we observed that the set of derived SNPs accurately approximates the genetic diversity across world populations. For each PC, the derived set of *k* SNPs (we used *k* = 500 throughout the analyses) captured genetic structure within different continental populations. Together, the top three PCs which explain most of the variance in the 1KG data, we observed that `ThreSPCA` selected 1500 meaningful, ancestry information preserving SNPs which leads to similar inference of population structure across the world as the original 1KG data with 360,498 SNPs. Annotating `ThreSPCA` derived variants further showed that they are interpretable and mostly missense in nature, thus likely disease causing. To interpret this, we mapped these variants to various traits in GWAS catalog and found that indeed these variants were mapped to different common traits such as body height, BMI, etc. which vary within and between populations across the world, sometimes leading to spurious associations due to population structure among populations [42]. These variants also mapped to various diseases, which vary across populations such as cardiovascular diseases [43]. Although the scale of the data used in this analysis is small when compared to large-scale genomic data, we observe that `ThreSPCA` is designed to handle biobank-scale datasets since it only need to run a randomized SVD/PCA analysis, which can be implemented efficiently in out-of-core settings [36]. `ThreSPCA` can also be used in GWAS as a population stratification correction step by identifying informative markers which highlight the ancestry stratification of cases/controls.

In summary, `ThreSPCA` provides a fast and provably accurate approximate method for computing SPCA. It provides a method to find interpretable markers in population genetics, which can immensely help understand population stratification, a major cause of spurious associations in GWAS. Also, it highlights the genetic sub-structure among different populations and the `ThreSPCA` derived variants are likely disease causing, often mapped to potential diseases and traits.

## Supporting information

zip

## Acknowledgements

PD and AC were partially supported by NSF III-10001674, NSF III10001225, and an IBM Faculty Award to PD. AB was supported by IBM. DPW and SZ would like to thank partial support from NSF grant No. CCF- 181584, Office of Naval Research (ONR) grant N00014-18-1-2562, National Institute of Health grant 5R01HG010798, and a Simons Investigator Award.

## Code Availability

See Appendix B for the ThreSPCA Python code availability.

## Appendices

## Appendix A SPCA via thresholding: Discussions and Proofs

The intuition behind Theorem 2.1 is that we can decompose the value of the optimal solution into the value contributed by the coordinates in *R*, the value contributed by the coordinates outside of *R,* and a cross term. The first term we can upper bound by the output of the algorithm, which maximizes with respect to the coordinates in *R*. For the latter two terms, we can upper bound the contribution due to the upper bound on the squared row norms of indices outside of *R* and due to the largest singular value of **U** being at most the trace of **A**.

We highlight that, as an intermediate step in the proof of Theorem 2.1, we need to prove the following Lemma A.1, which is very much at the heart of our proof of Theorem 2.1 and, unlike prior work, allows us to provide provably accurate bounds for the thresholding Algorithm 1. At a high level, the proof of Lemma A.1 first decomposes a basis for the columns spanned by **U** into those spanned by the top *ℓ* singular vectors and the remaining *n* – *ℓ* singular vectors. We then lower bound the contribution of the top *ℓ* singular vectors by upper bounding the contribution of the remaining *n* – *ℓ* singular vectors after noting that the largest remaining singular value is at most a 1/*ℓ*-fraction of the trace. We look at the detailed proof of Lemma A.1 below where we use the notation of Section 2.1. For notational convenience, let *σ*_1_,…, *σ_n_* be the diagonal entries of the matrix 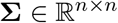, i.e., the singular values of **A**.

#### Lemma A.1

*Let* 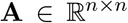 *be a PSD matrix and* 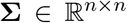 (*respectively*, 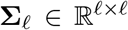*) be the diagonal matrix of all (respectively, top ℓ) singular values and let* 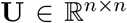 (*respectively*, 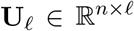*) be the matrix of all (respectively, top ℓ) singular vectors. Then, for all unit vectors* 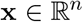,

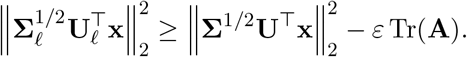

**Proof :** Let 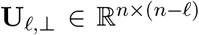
be a matrix whose columns form a basis for the subspace perpendicular to the subspace spanned by the columns of **U**_*ℓ*_. Similarly, let 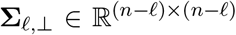 be the diagonal matrix of the bottom *n* – *ℓ* singular values of **A**. Notice that **U** = [**U**_*ℓ*_ **U**_*ℓ*,⊥_] and **Σ** = [**Σ**_*ℓ*_ **0**; **0 Σ**_*ℓ*,⊥_]; thus,

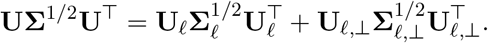

By the Pythagorean theorem,

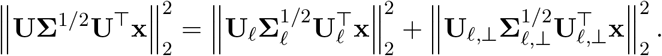

Using invariance properties of the vector two-norm and sub-multiplicativity, we get

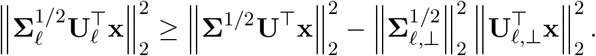

We conclude the proof by noting that 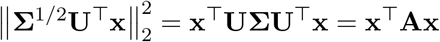 and

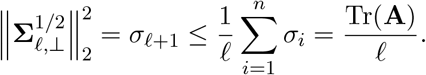

The inequality above follows since *σ*_1_ ≥ *σ*_2_ ≥ … *σ*_*ℓ*_ ≥ *σ*_*ℓ* +1_ ≥ … ≥ *σ_n_*. We conclude the proof by setting *ℓ* = 1/*ε*.

### Theorem 2.1

*Let k be the sparsity parameter and ε* ∈(0, 1] *be the accuracy parameter. Then, the vector* 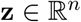 (*the output of Algorithm 1) has sparsity k/ε*^3^, *unit norm, and satisfies*

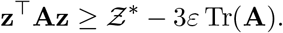

**Proof :** Let *R* ={*i*_1_,…, *i*_|*R*|_} be the set of indices of rows of **U**_*ℓ*_ columns of 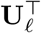) that have squared norm at least *ε*^2^/*k* and let 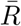 be its complement. Here |*R*| denotes the cardinality of the set *R* and 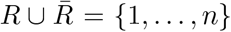. Let 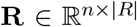 be the sampling matrix that selects the columns of **U**_*ℓ*_ whose indices are in the set *R* and let 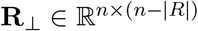 be the sampling matrix that selects the columns of **U**_*ℓ*_ whose indices are in the set 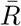. Thus, each column of **R** (respectively **R**_⊥)_ has a single non-zero entry, equal to one, corresponding to one of the |*R*| (respectively 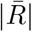) selected columns. Formally, **R**_*i*_*t*_;*t*_ = 1 for all *t* = 1,…, |*R*|, while all other entries of **R** (respectively **R**_⊥_) are set to zero; **R**_⊥_ can be defined analogously. The following properties are easy to prove: 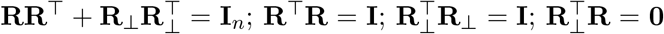. Recall that **x**\* is the optimal solution to the SPCA problem from eqn. (1). We proceed as follows:

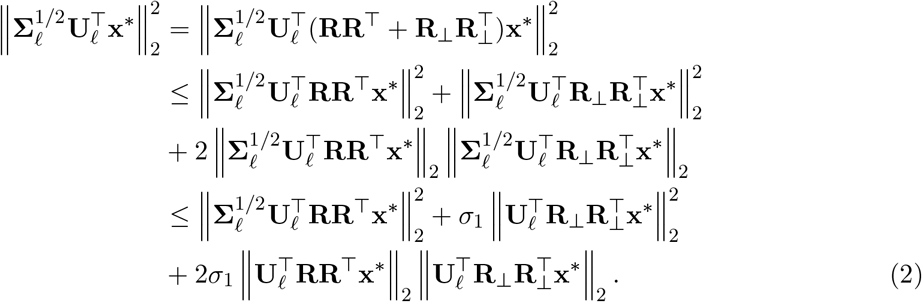

The above inequalities follow from the Pythagorean theorem and sub-multiplicativity. We now bound the second term in the right-hand side of the above inequality.

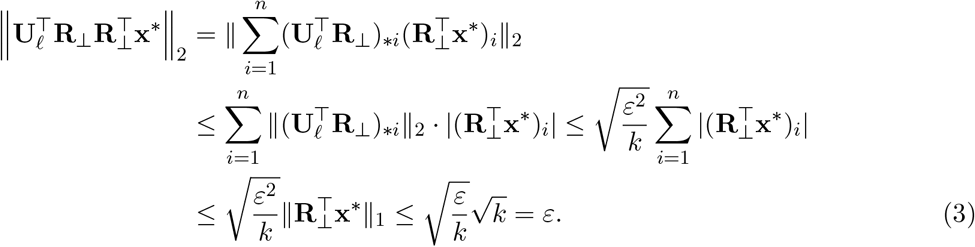

In the above derivations we use standard properties of norms and the fact that the columns of 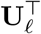 that have indices in the set 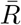 have squared norm at most *ε*^2^/*k*. The last inequality follows from 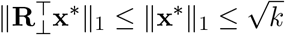, since **x**\* has at most *k* non-zero entries and Euclidean norm at most one.

Recall that the vector **y** of Algorithm 1 maximizes 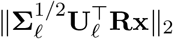 over all vectors **x** of appropriate dimensions (including **Rx**^*^) and thus

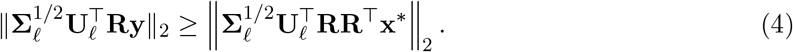

Combining eqns. (2), (3), and (4), we get that for sufficiently small *ε*,

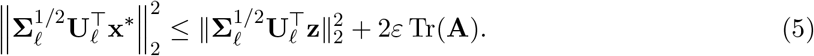

In the above we used **z** = **Ry** (as in Algorithm 1) and *σ*_1_ ≤Tr(**A**). Notice that

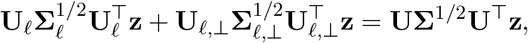

and using the Pythagorean theorem we get

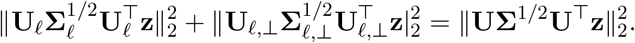

Using the unitary invariance of the two norm and dropping a non-negative term, we get the bound

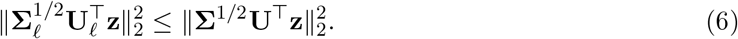

Combining eqns. (5) and (6), we conclude

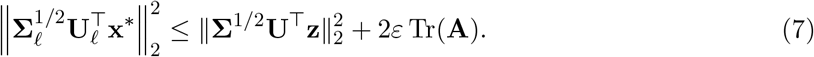

We now apply Lemma A.1 to the optimal vector **x**\* to get

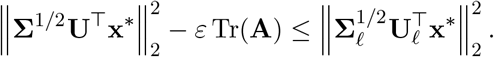

Combining with eqn. (7) we get

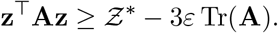

In the above we used 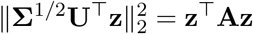 and 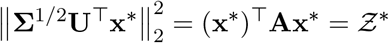. The result then follows from rescaling *ε*.

## Appendix B Code

A Python implementation of `ThreSPCA` can be found at: https://www.dropbox.com/s/czbvz5m5g42ey0n/ThreSPCA.py?dl=0.

## Appendix C Additional Experiments

### C.1 Simulated Studies

The genotype matrix 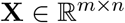 consisting of the simulated allele frequencies was generated using the algorithms of [33]. More specifically, we set **F** = **TS**, where 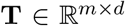 and 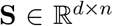, where *d* ≤*n* is the number of population groups. **S** is the indicator matrix that encapsulates structure with *n* individuals and contained in *d* populations. On the other hand, **T** characterizes how the structure is manifested in the allele frequencies of each SNP. Finally, projecting **S** onto the column space of **T**, we obtain the allele frequency matrix **F**. We sample **X** as a special case of **F** for the Pritchard-Stephens-Donelly (PSD) model. The allele frequency matrix was drawn from a beta distribution with the allele frequency, Fst distances and number of populations as parameters. We simulate **S** using i.i.d draws from the Dirichlet distribution with varying values of α, which denotes the parameter influencing the relatedness between the individuals. We show results for *α* = 0.01. *α* is directly proportional to the admixture of populations. Appendix Figure 1 shows the population structure observed in this simulated data.

**Fig. 1:**
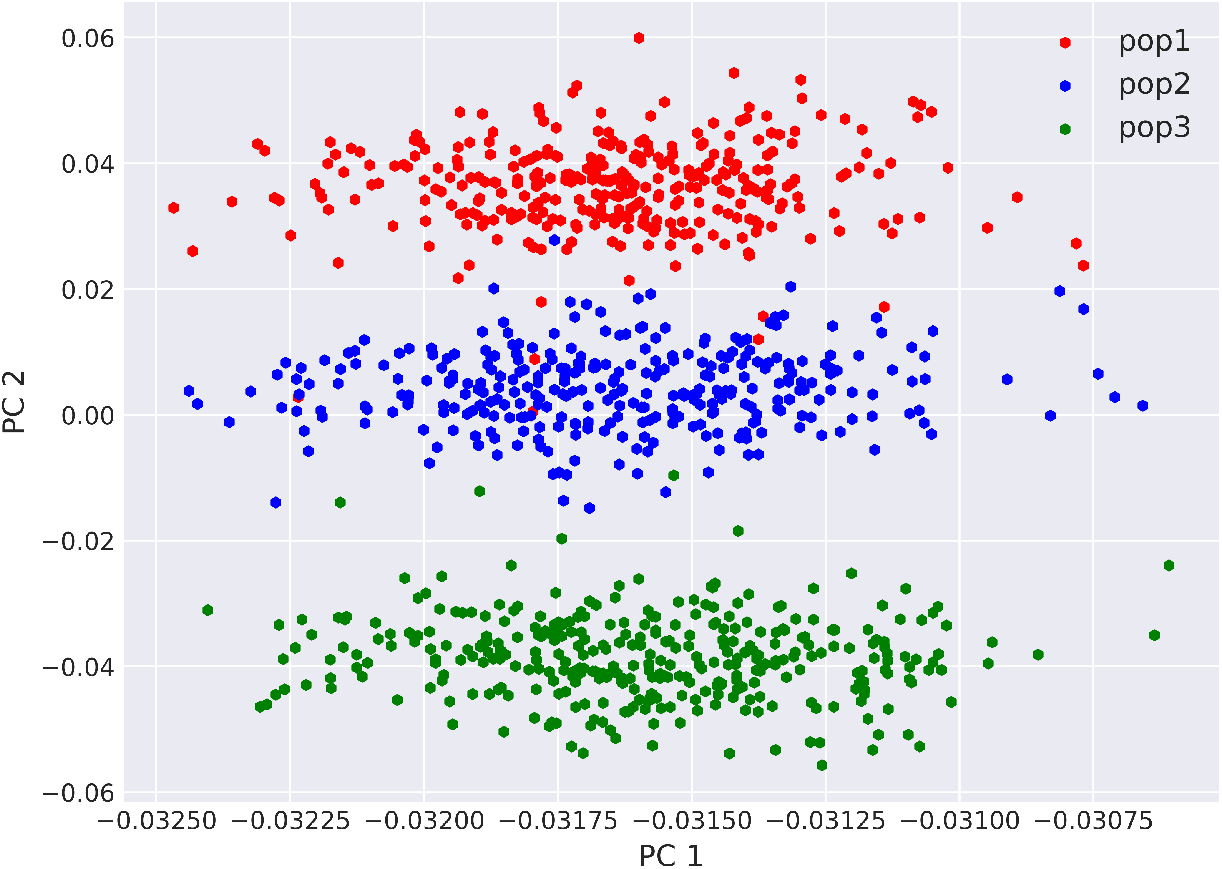
PCA plot of the simulated data with three distinct populations simulated from the PSD model with an *α* of 0.01, *n* = 1, 000, *m* = 10, 000 and *t* = 100

As it is difficult to establish notions of statistical significance in `ThreSPCA` capturing the ancestry informative markers from the original data, we simulated two different data set with varying numbers of individuals (*n*) and SNPs (*m*) and allowed *t* true SNPs which contribute to genetic ancestry. We varied *m*, *n* and *t* from {5000, 10000}, {500, 1000} and {100, 250, 500}, respectively.

For the random markers which do not contribute to the genetic differentiation we sampled the Fst distances between the individuals from a uniform distribution in the range {0, 0.005}which indicates minimum difference in populations. Thus, with this step we achieve the “true” markers contributing to genetic difference are the *t* SNPs and the remaining *m* — *t* SNPs, we conclude, are noise.

### C.2 Experiments on 1KG data

#### Population structure captured by PCA plots

We filtered the original 1KG data for the `ThreSPCA` derived *k* = 500 SNPs for each of the first three PCs and in the PCA plots we observe the population structure and the allele frequency distribution captured by each of the PCs. We clearly see that the SNPs from PC1 loadings are most frequent in the African populations or mixed populations of African ancestry (Appendix Figure 2). The PC2 SNPs are most frequent in East Asians but more or less commonly found in other populations as well (Appendix Figure 3) and the third PC SNPs are most frequent in South Asian populations (Appendix Figure 3). These shows that the SNP loadings from the top three PCs accurately captures the population structure across the world and merging them together to form a data set of 2503 individuals and 1500 SNPs, we not only capture the entire population structure in the PCA plot but also find some fine-grain substructure of populations (Figure 1).

**Fig. 2:**
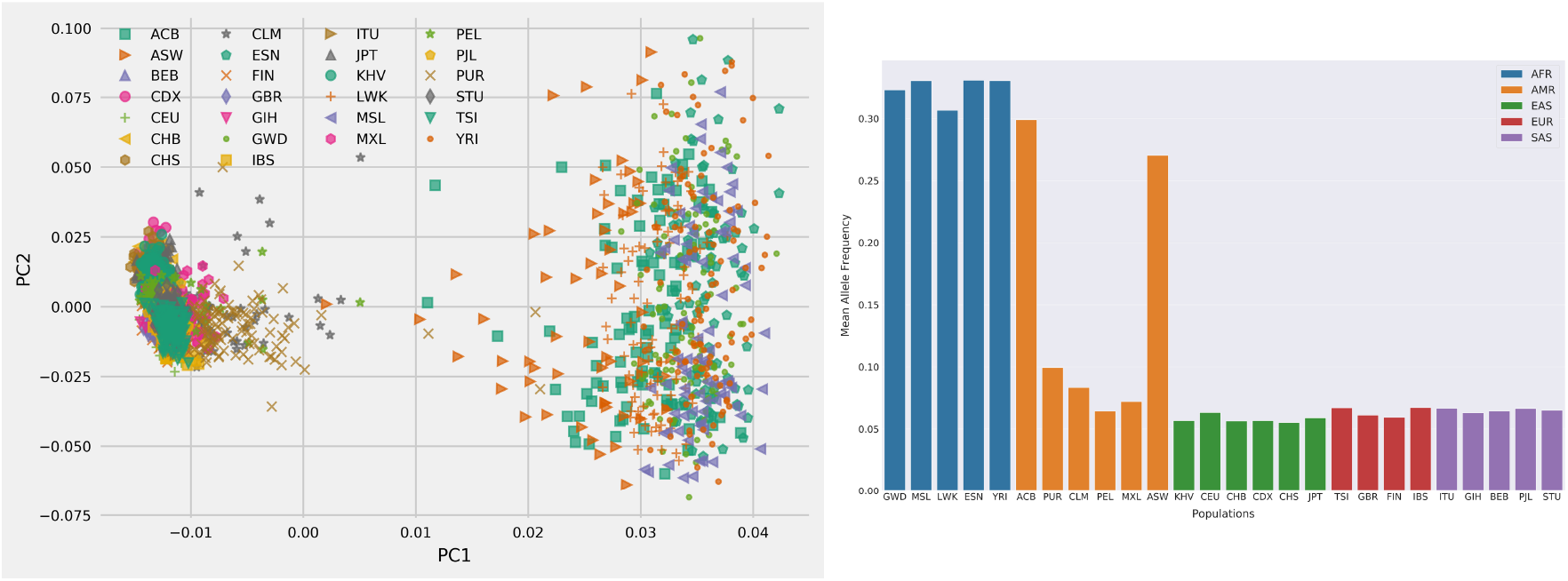
`ThreSPCA` with *k* = 500 obtained from PC1

#### Tuning input sparsity *k*

We tried a range of *k*’s varying it from 50 to 1500 and observed the *r*^2^ between the PCs derived from the original 1KG data and the 1500 SNPs derived from ThreSPCA. We observed that for the top two PCs the *r*^2^ is high from 0.96 to 0.99 wit the peak for both the PCs reaching around *k* = 500. PC1 continues to increase by two decimal points before saturing at *k* = 1000. Thus, we selected *k* = 500 for all the experiments as both the PCs reached their respective peaks.

**Fig. 3:**
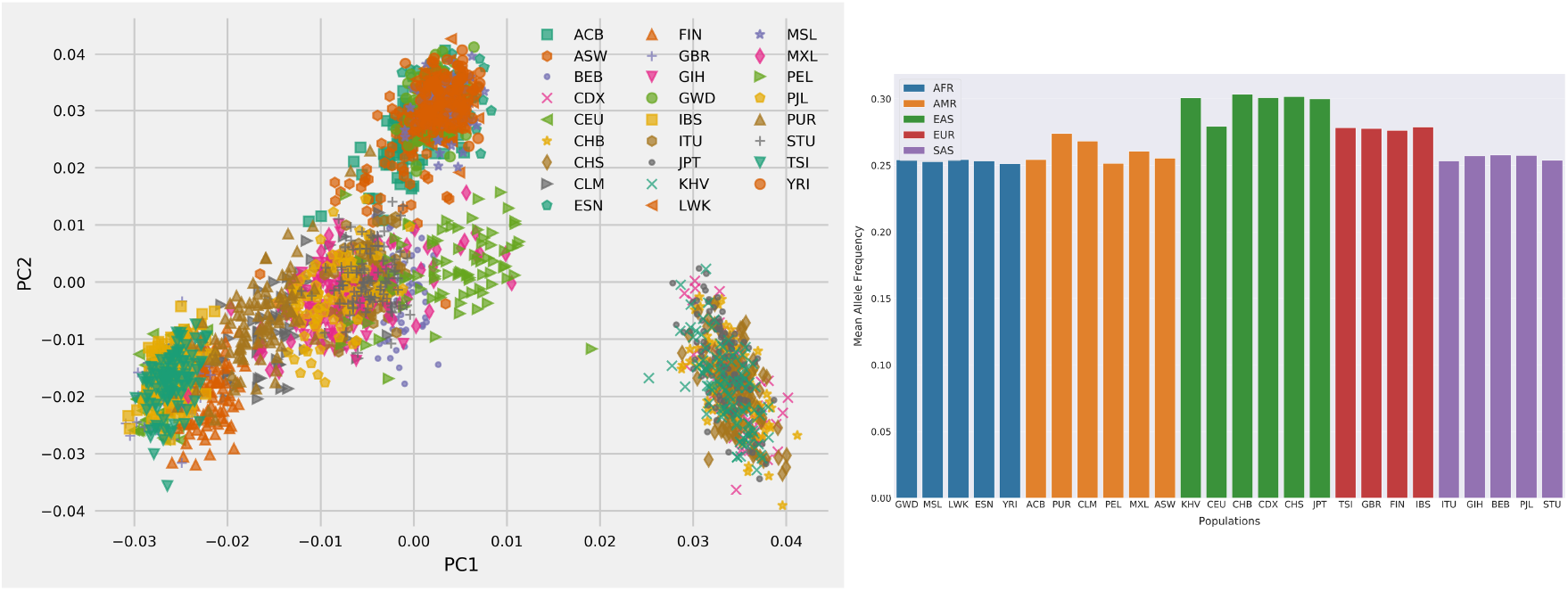
`ThreSPCA` with *k* = 500 obtained from PC2

**Fig. 4:**
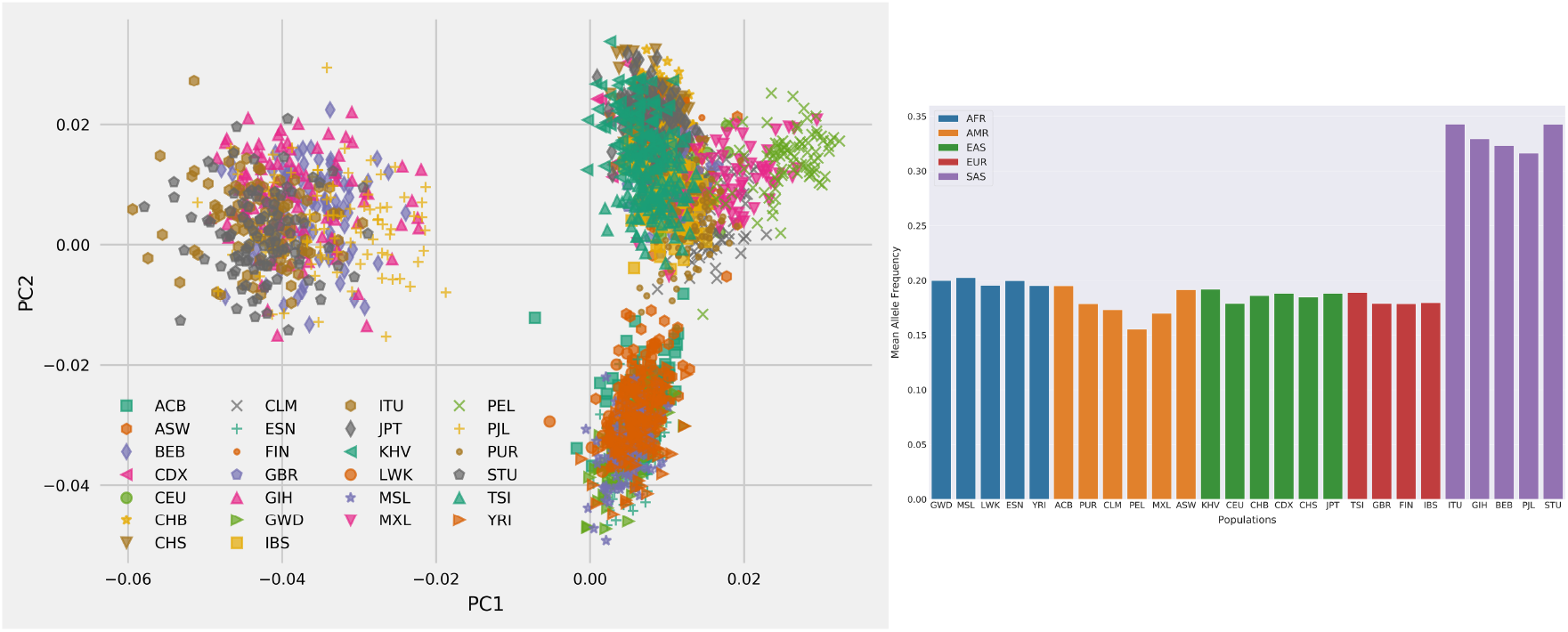
`ThreSPCA` with *k* = 500 obtained from PC3

Fig. 5: PCA plots of `ThreSPCA` filtered 1KG data for each PC with 500 SNPs.

**Fig. 6:**
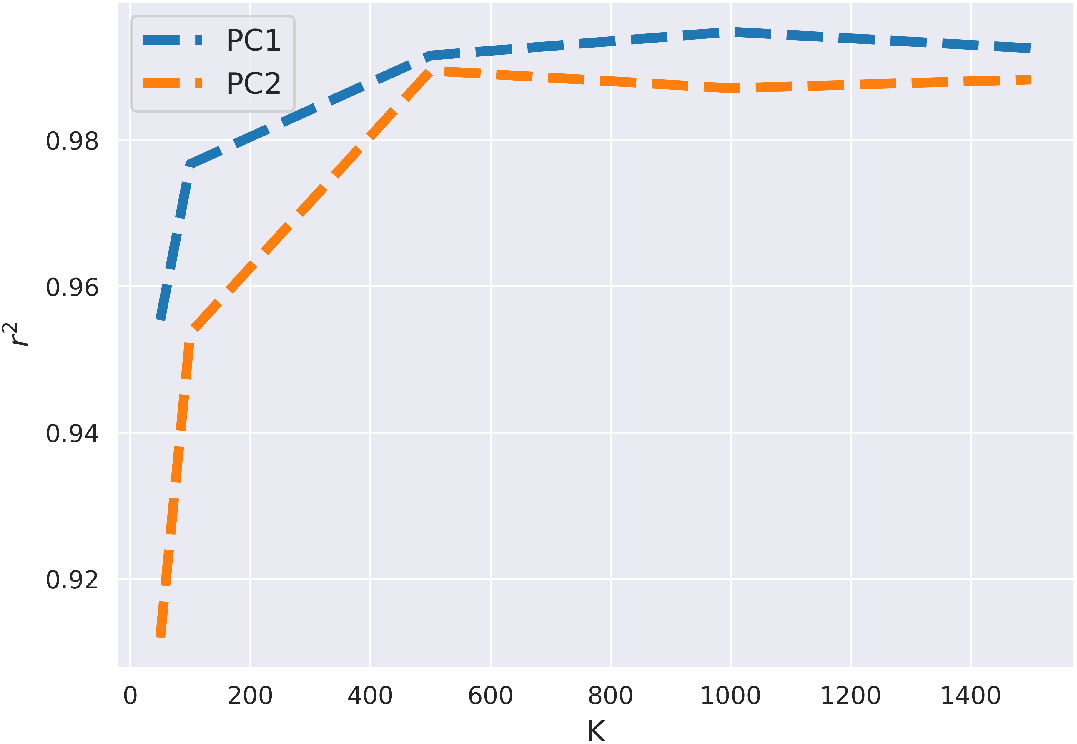
Line plot between the *r*^2^ between the PC scores of each PC obtained from ThreSPCA and the original PC from 1KG data with varying values of sparsity, *k*.

**Fig. 7:**
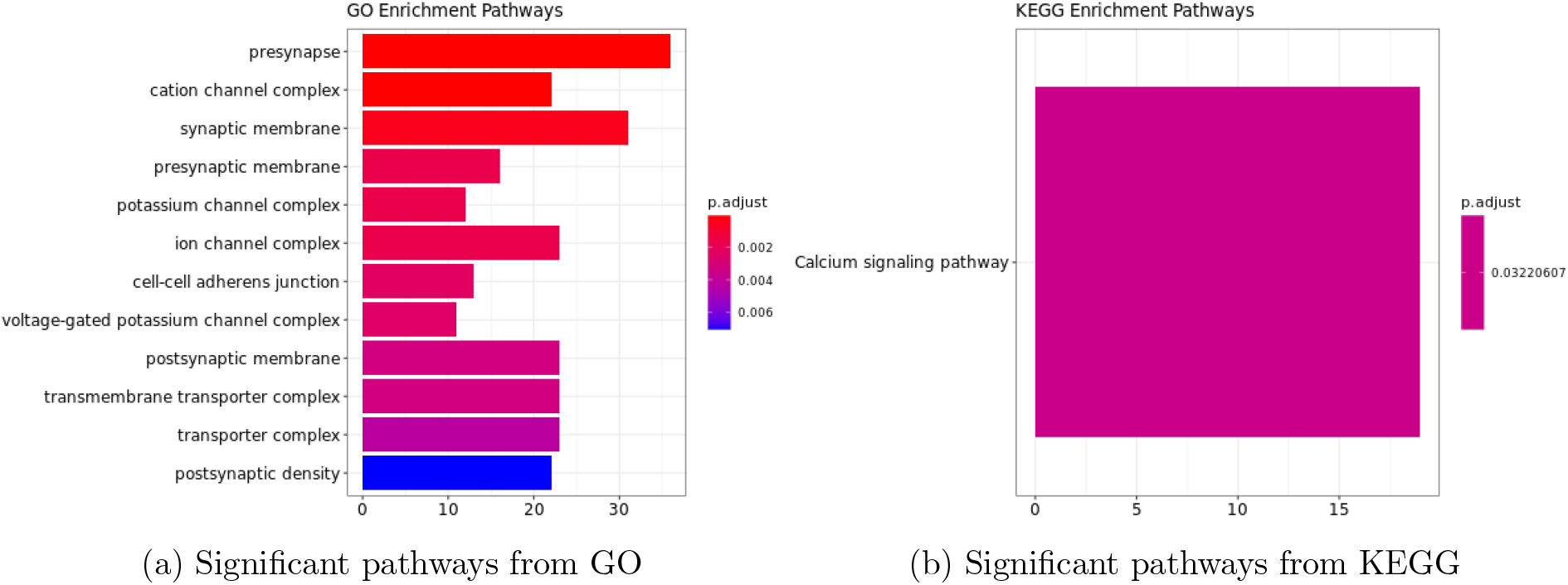
GO pathway analyses of the ThreSPCA informed variants, colored by p-values.

**Table 1:**
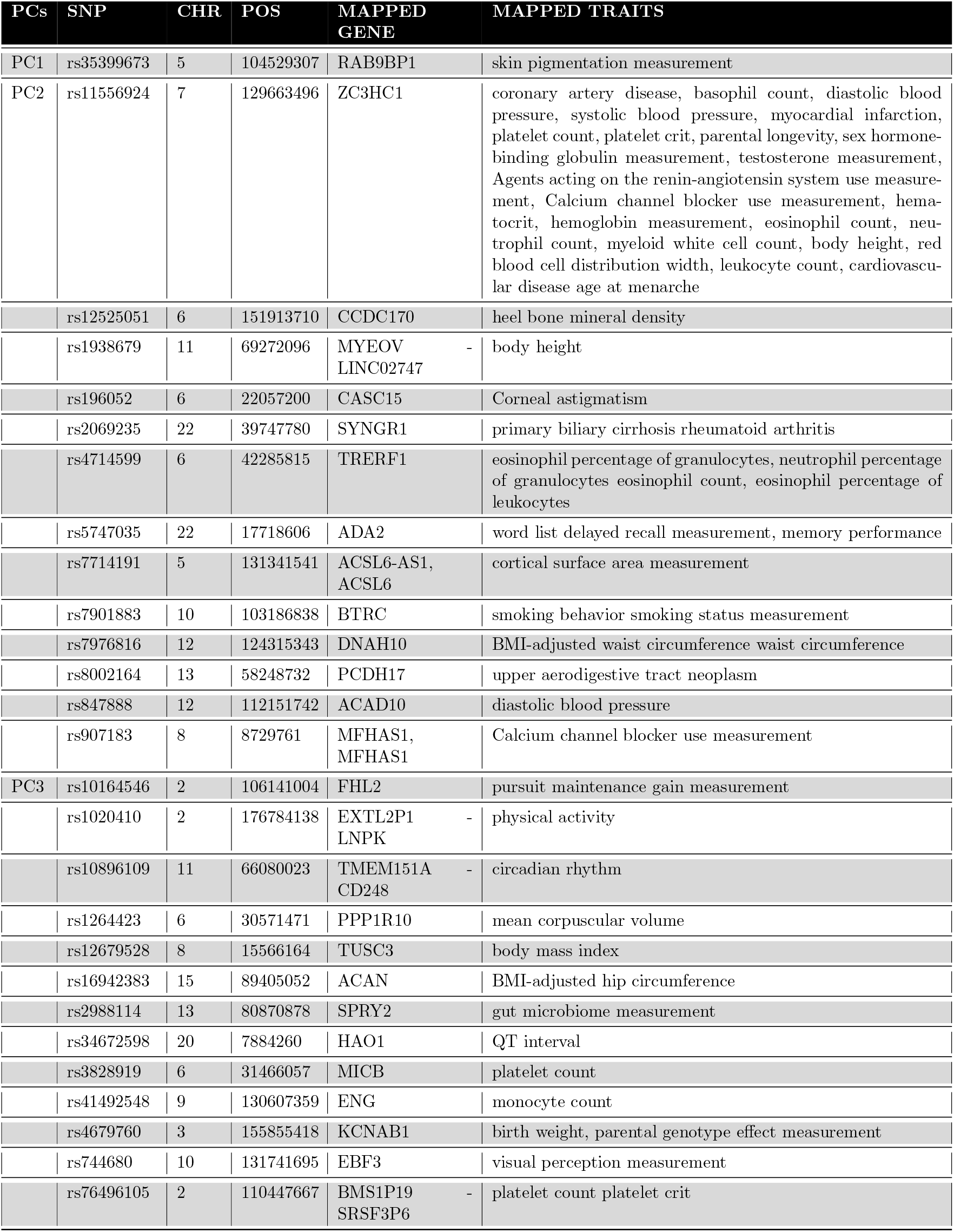
Traits and genes mapped in GWAS catalog from ThreSPCA informed variants.

### C.3 Comparing ThreSPCA with the state-of-the-art

#### C.3.1 Simulated data

We observed that increasing the threshold of true positives (markers which contribute to genetic structure) *t* led to an increase of the number of true positives observed in `ThreSPCA`. It found l

**Fig. 8:**
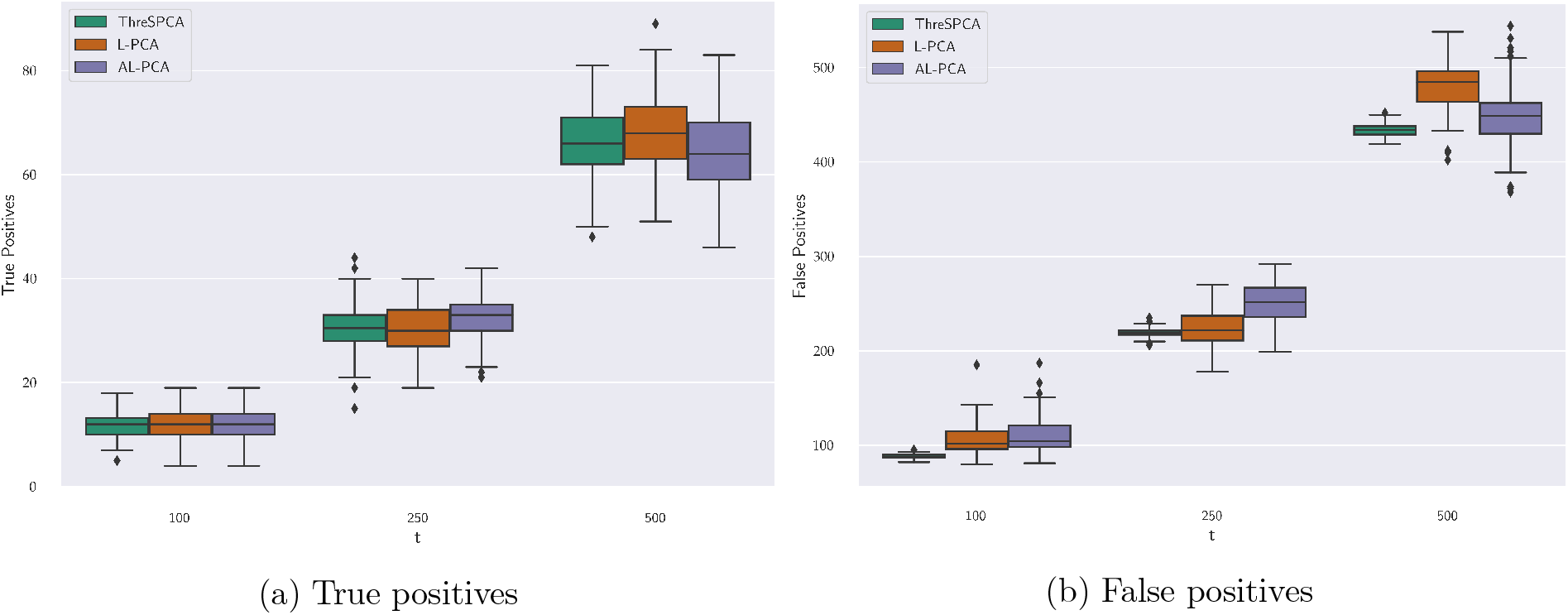
Box and whisker plots comparing between `ThreSPCA, L-PCA and AL-PCA` for true and false positives obtained from the simulated dataset of *m* = 5, 000 SNPs and *n* = 500 individuals and varying *t*.

#### C.3.2 Real data

On 1KG data we found perfect correlation with `ThreSPCA` and `AL-PCA` for PC1 and PC2 with *r*^2^ = 0.97 and 0.94 respectively.

**Fig. 9:**
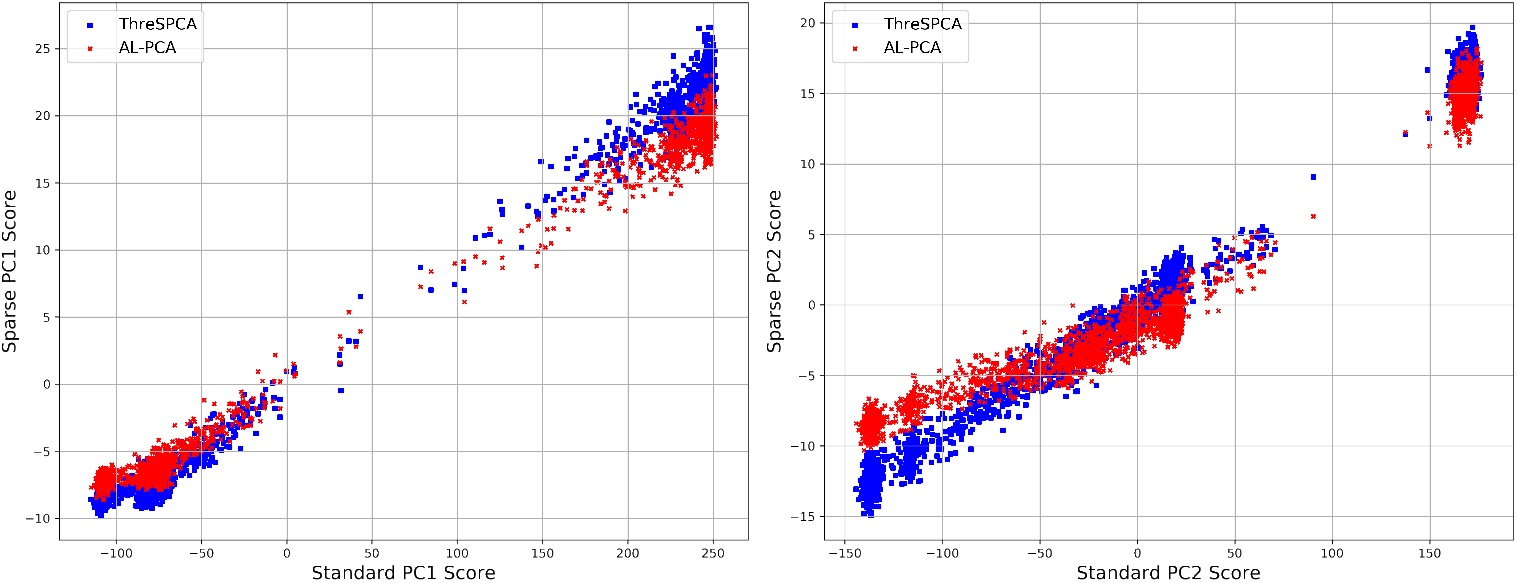
Relationship between the *PC scores* for PC1 and PC2 between `ThreSPCA` and `AL-PCA`

**Fig. 10:**
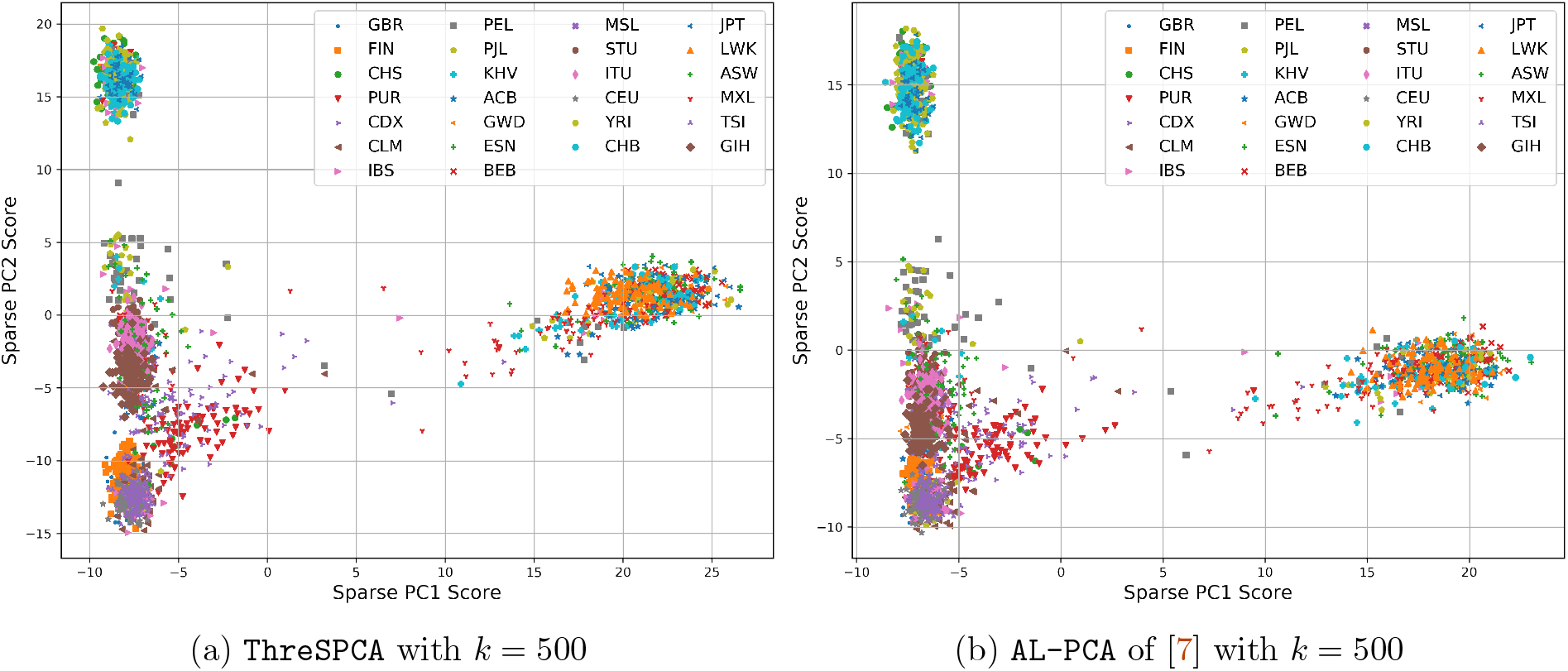
PCA plots of 1KG data with *PC scores*

**Fig. 11:**
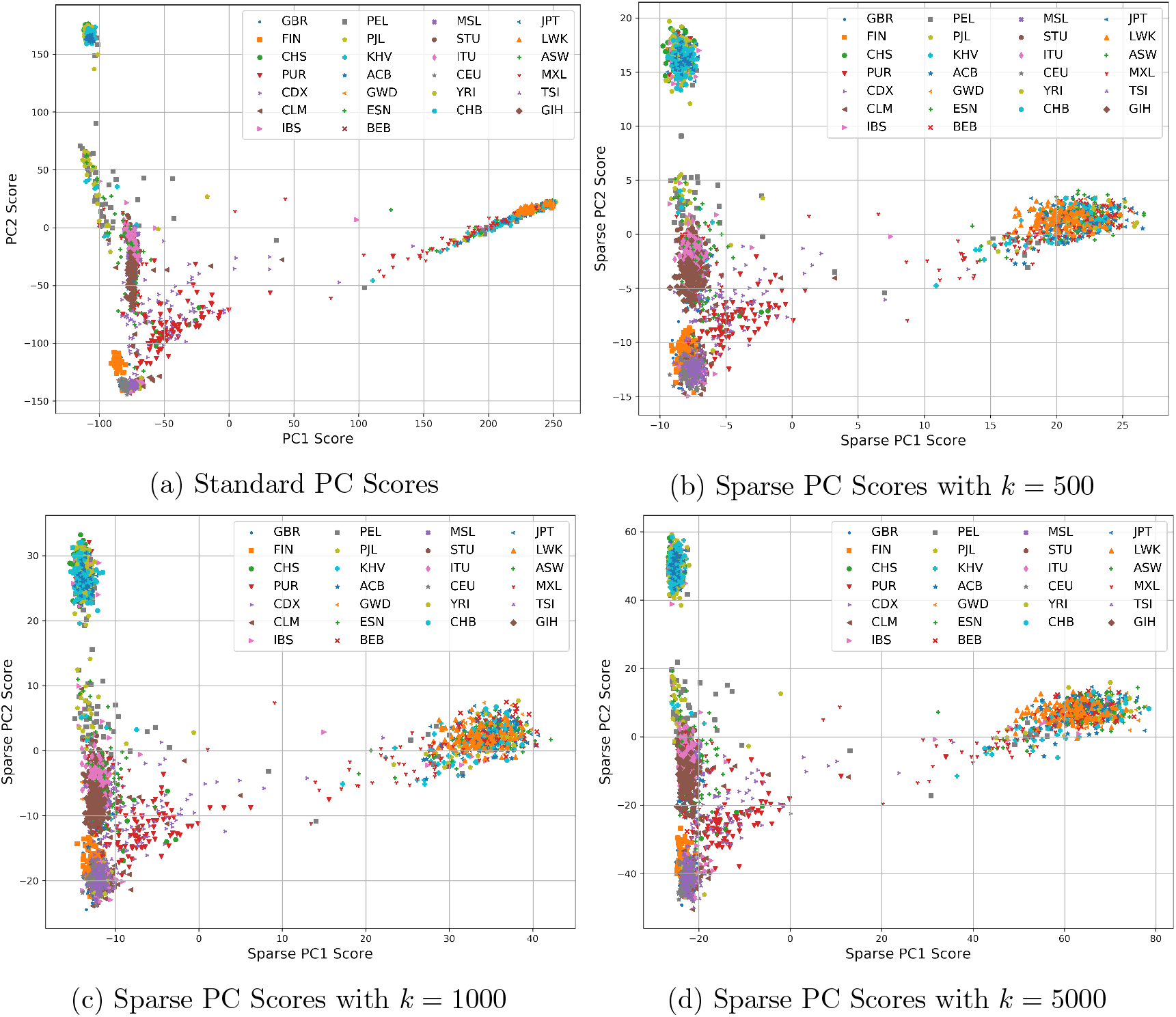
Scatterplots of the first two PC scores. Scatterplots of the first two PC scores from standard PCA (top left) and our `ThreSPCA` algorithm with *k* = 500 (top right), *k* = 1000 (bottom left) and *k* = 5000 (bottom right). Combination of different marker-shapes and marker-colors denote various population subgroups.

## Footnotes

1 Code available at https://github.com/aritra90/ThreSPCA

2 Recall that the *p*-th power of the *ℓ_p_* norm of a vector 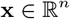 is defined as 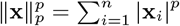 for 0 < *p* < ∞. For *p* = 0, ||*x*||_0_ is a semi-norm denoting the number of non-zero entries of *x*.

3 Each column of **R** has a single non-zero entry (set to one), corresponding to one of the |*R*| selected columns. Formally, **R**_*i*_*t*_,*t*_ = 1 for *t* = 1,…, |*R*|; all other entries of **R** are set to zero.

4 Results from `L-PCA` are qualitatively very similar to `AL-PCA` and we only report results for the latter.

