## Supplementary material for "A Fast, Provably Accurate Approximation Algorithm for Sparse Principal Component Analysis Reveals Human Genetic Variation Across the World": zip: abstract.tex

Principal component analysis (PCA) is a widely used dimension reduction technique in machine learning and multivariate statistics. To improve the interpretability of PCA, various approaches to obtain sparse principal direction loadings have been proposed, which are termed Sparse Principal Component Analysis (SPCA). In this paper, we present thresholding as a provably accurate, polynomial time, approximation algorithm for the SPCA problem, without imposing any restrictive assumptions on the input covariance matrix. Our thresholding algorithm using the Singular Value Decomposition is conceptually simple; much faster than current state-of-the-art; and performs well in practice. 
% On the negative side, our (novel) theoretical bounds do not accurately predict the strong practical performance of this approach.  The second algorithm solves a well-known semidefinite programming relaxation and then uses a novel, two step, deterministic thresholding scheme to compute a sparse principal vector. It works very well in practice and, remarkably, this solid practical performance is accurately predicted by our theoretical bounds, which bridge the theory-practice gap better than current state-of-the-art.
