## Supplementary material for "A Fast, Provably Accurate Approximation Algorithm for Sparse Principal Component Analysis Reveals Human Genetic Variation Across the World": zip: appendixA.tex

\begin{appendices}
%\appendix
\section{SPCA via thresholding: Discussions and Proofs}\seclab{sxn:threshold:proofs}

The intuition behind \thmref{thm:pcpmain} is that we can decompose the value of the optimal solution into the value contributed by the coordinates in $R$, the value contributed by the coordinates outside of $R$, and a cross term. 
The first term we can upper bound by the output of the algorithm, which maximizes with respect to the coordinates in $R$. 
For the latter two terms, we can upper bound the contribution due to the upper bound on the squared row norms of indices outside of $R$ and due to the largest singular value of $\Ub$ being at most the trace of $\Ab$. 
%We defer the full proof of \thmref{thm:pcpmain} to the supplementary material. 

We highlight that, as an intermediate step in the proof of \thmref{thm:pcpmain}, we need to prove the following \lemref{lem:sz:pcp}, which is very much at the heart of our proof of \thmref{thm:pcpmain} and, unlike prior work, allows us to provide provably accurate bounds for the thresholding~\algref{alg:sparse:pca}.
%
%\lemref{lem:sz:pcp} is very much at the heart of our proof of \thmref{thm:pcpmain} and, unlike prior work, allows us to provide provably accurate bounds for the thresholding~\algref{alg:sparse:pca}. 
At a high level, the proof of \lemref{lem:sz:pcp} first decomposes a basis for the columns spanned by $\Ub$ into those spanned by the top $\ell$ singular vectors and the remaining $n-\ell$ singular vectors. 
We then lower bound the contribution of the top $\ell$ singular vectors by upper bounding the contribution of the remaining $n-\ell$ singular vectors after noting that the largest remaining singular value is at most a $1/\ell$-fraction of the trace. 
%For additional details, we defer the full proof to the supplementary material.
We look at the detailed proof of \lemref{lem:sz:pcp} below where we use the notation of~\secref{sec:pcp}. 
For notational convenience, let $\sigma_1,\ldots,\sigma_n$ be the diagonal entries of the matrix $\Sigmab \in \mathbb{R}^{n \times n}$, i.e., the singular values of $\Ab$.

%\lemszpcp*
\begin{restatable}{lemma}{lemszpcp}
\lemlab{lem:sz:pcp}
Let $\A\in\mathbb{R}^{n\times n}$ be a PSD matrix and $\Sigmab \in \mathbb{R}^{n \times n}$ (respectively,
$\Sigmab_{\ell} \in \mathbb{R}^{\ell \times \ell}$) be the diagonal matrix of all (respectively, top $\ell$) singular values and let
$\Ub \in \mathbb{R}^{n \times n}$ (respectively, $\Ub_{\ell} \in \mathbb{R}^{n \times \ell}$) be the matrix of all (respectively, top $\ell$) singular vectors. Then, for all unit vectors $\xb \in \mathbb{R}^n$,
%
$$\norm{\Sigmab_{\ell}^{1/2} \Ub_{\ell}^\top\x}_2^2 \ge \norm{\Sigmab^{1/2} \Ub^\top \x}_2^2- \eps\trace(\A).$$
%
\end{restatable}

\begin{proof}
%
Let $\Ub_{\ell,\perp} \in \mathbb{R}^{n \times (n-\ell)}$ be a matrix whose columns form a basis for the subspace perpendicular to the subspace spanned by the columns of $\Ub_{\ell}$. Similarly, let $\Sigmab_{\ell,\perp} \in \mathbb{R}^{(n-\ell) \times (n-\ell)}$ be the diagonal matrix of the bottom $n-\ell$ singular values of $\Ab$. Notice that $\Ub = [\Ub_{\ell}\ \ \Ub_{\ell,\perp}]$ and $\Sigmab = [\Sigmab_{\ell}\ \ \zero;\ \zero\ \ \Sigmab_{\ell,\perp}]$; thus,
%
\begin{flalign*}
%
\Ub\Sigmab^{1/2}\Ub^{\top} = \Ub_{\ell}\Sigmab_{\ell}^{1/2}\Ub_{\ell}^{\top} +
\Ub_{\ell,\perp}\Sigmab_{\ell,\perp}^{1/2}\Ub_{\ell,\perp}^{\top}.
%
\end{flalign*}
%
By the Pythagorean theorem,
%
\begin{align*}
%
\norm{\Ub\Sigmab^{1/2}\Ub^{\top}\x}_2^2=
\norm{\Ub_{\ell}\Sigmab_{\ell}^{1/2}\Ub_{\ell}^{\top}\x}_2^2 +
\norm{\Ub_{\ell,\perp}\Sigmab_{\ell,\perp}^{1/2}\Ub_{\ell,\perp}^{\top}\x}_2^2.
%
\end{align*}
%
Using invariance properties of the vector two-norm and sub-multiplicativity, we get
%
\begin{align*}
%
\norm{\Sigmab_{\ell}^{1/2}\Ub_{\ell}^{\top}\x}_2^2 \geq
%
\norm{\Sigmab^{1/2}\Ub^{\top}\x}_2^2-
%
\norm{\Sigmab_{\ell,\perp}^{1/2}}_2^2\norm{\Ub_{\ell,\perp}^{\top}\x}_2^2.
%
\end{align*}
%
We conclude the proof by noting that $\norm{\Sigmab^{1/2}\Ub^{\top}\x}_2^2 = \x^\top \Ub\Sigmab\Ub^{\top}\x = \x^\top \Ab \x$ and
%
$$\norm{\Sigmab_{\ell,\perp}^{1/2}}_2^2 = \sigma_{\ell+1} \leq \frac{1}{\ell}\sum_{i=1}^n \sigma_i = \frac{\trace(\Ab)}{\ell}.$$
%
The inequality above follows since $\sigma_1\geq \sigma_2\geq \ldots \sigma_\ell \geq \sigma_{\ell+1}\geq \ldots \geq \sigma_n$. We conclude the proof by setting $\ell = 1/\varepsilon$.
\end{proof}


\thmpcpmain*
\begin{proof}
%
Let $R = \{i_1,\ldots,i_{|R|}\}$ be the set of indices of rows of $\Ub_{\ell}$ (columns of $\Ub_{\ell}^\top$) that have squared norm at least $\varepsilon^2/k$ and let $\bar{R}$ be its complement. Here $|R|$ denotes the cardinality of the set $R$ and $R \cup \bar{R} = \{1,\ldots,n\}$. Let $\Rb \in \mathbb{R}^{n \times |R|}$ be the sampling matrix that selects the columns of $\Ub_{\ell}$ whose indices are in the set $R$ and let
%
$\Rb_{\perp} \in \mathbb{R}^{n \times (n-|R|)}$ be the sampling matrix that selects the columns of $\Ub_{\ell}$ whose indices are in the set $\bar{R}$.
Thus, each column of $\Rb$ (respectively $\Rb_{\perp}$) has a single non-zero entry, equal to one, corresponding to one of the $|R|$ (respectively $|\bar{R}|$) selected columns. Formally, $\Rb_{i_t,t}=1$ for all $t=1,\ldots, |R|$, while all other entries of $\Rb$ (respectively $\Rb_{\perp}$) are set to zero; $\Rb_{\perp}$ can be defined analogously.
%
The following properties are easy to prove: $\Rb\Rb^{\top}+\Rb_{\perp}\Rb_{\perp}^{\top} = \Ib_n$; $\Rb^{\top}\Rb=\Ib$; $\Rb_{\perp}^{\top}\Rb_{\perp}=\Ib$;
$\Rb_{\perp}^{\top}\Rb=\zero$.
%
Recall that $\xb^*$ is the optimal solution to the SPCA problem from eqn.~(\ref{eqn:spca}). We proceed as follows:

%
\begin{flalign}
%
\norm{\Sigmab_{\ell}^{1/2}\U_{\ell}^\top\x^*}_2^2&=\norm{\Sigmab_{\ell}^{1/2}\U_{\ell}^\top(\Rb\Rb^{\top}+\Rb_{\perp}\Rb_{\perp}^{\top})\x^*}_2^2\nonumber\\
%
&\leq \norm{\Sigmab_{\ell}^{1/2}\U_{\ell}^\top\Rb\Rb^{\top}\x^*}_2^2+\norm{\Sigmab_{\ell}^{1/2}\U_{\ell}^\top\Rb_{\perp}\Rb_{\perp}^{\top}\x^*}_2^2\nonumber\\
&+2\norm{\Sigmab_{\ell}^{1/2}\U_{\ell}^\top\Rb\Rb^{\top}\x^*}_2\norm{\Sigmab_{\ell}^{1/2}\U_{\ell}^\top\Rb_{\perp}\Rb_{\perp}^{\top}\x^*}_2\nonumber\\
%
&\leq \norm{\Sigmab_{\ell}^{1/2}\U_{\ell}^\top\Rb\Rb^{\top}\x^*}_2^2+\sigma_1\norm{\U_{\ell}^\top\Rb_{\perp}\Rb_{\perp}^{\top}\x^*}_2^2\nonumber\\
&+2\sigma_1\norm{\U_{\ell}^\top\Rb\Rb^{\top}\x^*}_2\norm{\U_{\ell}^\top\Rb_{\perp}\Rb_{\perp}^{\top}\x^*}_2\label{eqn:sdp:svd:1}.
%
\end{flalign}
%
The above inequalities follow from the Pythagorean theorem and sub-multiplicativity. 
We now bound the second term in the right-hand side of the above inequality.
%
\begin{flalign}
  %
  \norm{\U_{\ell}^\top\Rb_{\perp}\Rb_{\perp}^{\top}\x^*}_2 &=  \|\sum_{i=1}^n (\U_{\ell}^\top\Rb_{\perp})_{*i}(\Rb_{\perp}^{\top}\x^*)_i\|_2\nonumber\\
  %
  &\leq \sum_{i=1}^n \|(\U_{\ell}^\top\Rb_{\perp})_{*i}\|_2 \cdot |(\Rb_{\perp}^{\top}\x^*)_i|\leq \sqrt{\frac{\varepsilon^2}{k}}\sum_{i=1}^n |(\Rb_{\perp}^{\top}\x^*)_i|\nonumber\\
  %
  &\leq \sqrt{\frac{\varepsilon^2}{k}}\|\Rb_{\perp}^{\top}\x^*\|_1\leq \sqrt{\frac{\varepsilon}{k}}\sqrt{k} = \varepsilon.\label{eqn:sdp:svd:2}
  %
\end{flalign}
%
In the above derivations we use standard properties of norms and the fact that the columns of $\U_{\ell}^\top$ that have indices in the set $\bar{R}$ have squared norm at most $\varepsilon^2/k$. The last inequality follows from $\|\Rb_{\perp}^{\top}\x^*\|_1 \leq \|\x^*\|_1 \leq \sqrt{k}$, since $\xb^*$ has at most $k$ non-zero entries and Euclidean norm at most one.

Recall that the vector $\yb$ of~\algref{alg:sparse:pca} maximizes $\|\Sigmab_{\ell}^{1/2}\U_{\ell}^\top\Rb\xb\|_2$ over all vectors $\xb$ of appropriate dimensions (including $\Rb \x^*$) and thus
%
\begin{flalign}\label{eqn:sdp:svd:3}
  \|\Sigmab_{\ell}^{1/2}\U_{\ell}^\top\Rb\yb\|_2 \geq \norm{\Sigmab_{\ell}^{1/2}\U_{\ell}^\top\Rb\Rb^{\top}\x^*}_2.
\end{flalign}
%
Combining eqns.~(\ref{eqn:sdp:svd:1}),~(\ref{eqn:sdp:svd:2}), and~(\ref{eqn:sdp:svd:3}), we get that for sufficiently small $\eps$, 
%
\begin{flalign}\label{eqn:sdp:svd:4}
%
\norm{\Sigmab_{\ell}^{1/2}\U_{\ell}^\top\x^*}_2^2 \leq \|\Sigmab_{\ell}^{1/2}\U_{\ell}^\top\zb\|_2^2 + 2\varepsilon\trace(\Ab).
%
\end{flalign}
%
In the above we used $\zb = \Rb\yb$ (as in~\algref{alg:sparse:pca}) and $\sigma_1 \leq \trace(\Ab)$. Notice that
%
$$\U_{\ell}\Sigmab_{\ell}^{1/2}\U_{\ell}^\top\zb + \U_{\ell,\perp}\Sigmab_{\ell,\perp}^{1/2}\U_{\ell,\perp}^\top\zb=
\U\Sigmab^{1/2}\U^\top\zb,$$
%
and using the Pythagorean theorem we get
%
$$\|\U_{\ell}\Sigmab_{\ell}^{1/2}\U_{\ell}^\top\zb\|_2^2 + \|\U_{\ell,\perp}\Sigmab_{\ell,\perp}^{1/2}\U_{\ell,\perp}^\top\zb|_2^2=
\|\U\Sigmab^{1/2}\U^\top\zb\|_2^2.$$
%
Using the unitary invariance of the two norm and dropping a non-negative term, we get the bound
%
\begin{flalign}\label{eqn:sdp:svd:5}
\|\Sigmab_{\ell}^{1/2}\U_{\ell}^\top\zb\|_2^2 \leq
\|\Sigmab^{1/2}\U^\top\zb\|_2^2.
\end{flalign}
%
Combining eqns.~(\ref{eqn:sdp:svd:4}) and~(\ref{eqn:sdp:svd:5}), we conclude
%
\begin{flalign}\label{eqn:sdp:svd:6}
%
\norm{\Sigmab_{\ell}^{1/2}\U_{\ell}^\top\x^*}_2^2 \leq \|\Sigmab^{1/2}\U^\top\zb\|_2^2 + 2\varepsilon\trace(\Ab).
%
\end{flalign}
%
We now apply~\lemref{lem:sz:pcp} to the optimal vector $\xb^*$ to get
%
$$\norm{\Sigmab^{1/2} \Ub^\top \xb^*}_2^2- \eps\trace(\A) \leq \norm{\Sigmab_{\ell}^{1/2} \Ub_{\ell}^\top\xb^*}_2^2.$$
%
Combining with eqn.~(\ref{eqn:sdp:svd:6}) we get
%
$$\zb^\top \Ab \zb \geq\mathcal{Z}^* - 3\eps\trace(\A).$$
%
In the above we used
$\|\Sigmab^{1/2}\U^\top\zb\|_2^2 = \zb^\top \Ab \zb$ and
$\norm{\Sigmab^{1/2} \Ub^\top\xb^*}_2^2=(\xb^*)^\top \Ab \xb^* = \mathcal{Z}^*$. 
The result then follows from rescaling $\eps$. 
\end{proof}

\pagebreak 
\section{Code}\label{app:code}

A Python implementation of \texttt{ThreSPCA} can be found at:\\ \url{https://github.com/aritra90/ThreSPCA}.

\section{Additional Experiments}\label{app:expt}
\setcounter{figure}{0}    

\subsection{Simulated Studies}
The genotype matrix $\Xb \in \mathbb{R}^{m \times n}$ consisting of the simulated allele frequencies was generated using the algorithms of~\cite{pritchard2000inference}. More specifically, we set $\Fb = \Tb\Sb$, where $\Tb \in \mathbb{R}^{m \times d}$ and $\Sb \in \mathbb{R}^{d \times n}$, where $d \leq n$ is the number of population groups. $\Sb$ is the indicator matrix that encapsulates structure with n individuals and contained in d populations. On the other hand, $\Tb$ characterizes how the structure is manifested in the allele frequencies of each SNP. Finally, projecting $\Sb$ onto the column space of $\Tb$, we obtain the allele frequency matrix $\Fb$. We sample $\Xb$ as a special case of $\Fb$ for the Pritchard-Stephens-Donelly (PSD) model. 
The allele frequency matrix was drawn from a beta distribution with the allele frequency, Fst distances and number of populations as parameters. We simulate $\Sb$ using i.i.d draws from the Dirichlet distribution with varying values of $\alpha$, which denotes the parameter influencing the relatedness between the individuals. We show results for $\alpha = 0.01$. $\alpha$ is directly proportional to the admixture of populations. Appendix Figure~\ref{fig:simPC} shows the population structure observed in this simulated data.

\begin{figure*}[!htbp]
    \centering
    \includegraphics[scale=0.6]{figures/PSD0.01_m10K_n1K_t100_PCs.png}
    \caption{PCA plot of the simulated data with three distinct populations simulated from the PSD model with an $\alpha$ of 0.01, $n=1,000$, $m=10,000$ and $t=100$}
    \label{fig:simPC}
\end{figure*}

As it is difficult to establish notions of statistical significance in \texttt{ThreSPCA} capturing the ancestry informative markers from the original data, we simulated two different data set with varying numbers of individuals ($n$) and SNPs ($m$) and allowed $t$ true SNPs which contribute to genetic ancestry. We varied $m$, $n$ and $t$ from \{5000, 10000\}, \{500, 1000\} and \{100, 250, 500\}, respectively. For the random markers which do not contribute to the genetic differentiation we sampled the Fst distances between the individuals from a uniform distribution in the range $\{0,0.005\}$ which indicates minimum difference in populations. Thus, with this step we achieve the ``true" markers contributing to genetic difference are the $t$ SNPs and the remaining $m-t$ SNPs, we conclude, are noise. 

\subsection{Experiments on 1KG data}
\paragraph{Population structure captured by PCA plots.}
We filtered the original 1KG data for the \texttt{ThreSPCA} derived $k=500$ SNPs for each of the first three PCs and in the PCA plots we observe the population structure and the allele frequency distribution captured by each of the PCs. We clearly see that the SNPs from PC1 loadings are most frequent in the African populations or mixed populations of African ancestry (Appendix Figure~\ref{fig:SPCA_PC1}). The PC2 SNPs are most frequent in East Asians but more or less commonly found in other populations as well (Appendix Figure~\ref{fig:SPCA_PC2}) and the third PC SNPs are most frequent in South Asian populations (Appendix Figure~\ref{fig:SPCA_PC2}). These shows that the SNP loadings from the top three PCs accurately captures the population structure across the world and merging them together to form a data set of 2503 individuals and 1500 SNPs, we not only capture the entire population structure in the PCA plot but also find some fine-grain substructure of populations (Figure~\ref{fig:1KGcompare}). 
%Each PC with 500 SNPs
\begin{figure*}[!htbp]

      \centering
      \includegraphics[scale=0.4]{figures/ThreSPCA_PC1_k500.png}
      \caption{\texttt{ThreSPCA} with $k=500$ obtained from PC1}
      \label{fig:SPCA_PC1}
\end{figure*}
\begin{figure*}[!htbp]

      \centering
      \includegraphics[scale=0.4]{figures/ThreSPCA_PC2_k500.png}
      \caption{\texttt{ThreSPCA} with $k=500$ obtained from PC2}
      \label{fig:SPCA_PC2}
\end{figure*}
\begin{figure*}[!htbp]
      \centering
      \includegraphics[scale=0.4]{figures/ThreSPCA_PC3_k500.png}
      \caption{\texttt{ThreSPCA} with $k=500$ obtained from PC3}
      \label{fig:SPCA_PC3}
    %
    \caption{PCA plots of \texttt{ThreSPCA} filtered 1KG data for each PC with 500 SNPs.}
\end{figure*}
\paragraph{Tuning input sparsity $k$.}
We tried a range of $k$'s varying it from 50 to 1500 and observed the $r^2$ between the PCs derived from the original 1KG data and the 1500 SNPs derived from \texttt{ThreSPCA}. We observed that for the top two PCs the $r^2$ is high from 0.96 to 0.99 wit the peak for both the PCs reaching around $k=500$. PC1 continues to increase by two decimal points before saturing at $k=1000$. Thus, we selected $k=500$ for all the experiments as both the PCs reached their respective peaks.

\begin{figure*}[!htbp]
    \centering
    \includegraphics[scale=0.6]{figures/k_by_r2.png}
    \caption{Line plot between the $r^2$ between the PC scores of each PC obtained from \texttt{ThreSPCA} and the original PC from 1KG data with varying values of sparsity, $k$.}
    \label{fig:kbyr2}
\end{figure*}

\begin{table}[htbp]
  \centering
    %\begin{tabular}{|c|c|c|c|c|c|}
    \resizebox{0.925\textwidth}{!}{
    \begin{tabular}{|p{0.8cm}|p{2cm}|p{1cm}|p{1.7cm}|p{3cm}|p{10cm}|}
    \toprule
    \rowcolor[rgb]{ 0,  0,  0} \textcolor[rgb]{ 1,  1,  1}{\textbf{PCs}} & \textcolor[rgb]{ 1,  1,  1}{ \textbf{SNP}} & \textcolor[rgb]{ 1,  1,  1}{\textbf{CHR}} & \textcolor[rgb]{ 1,  1,  1}{\textbf{POS}} & \textcolor[rgb]{ 1,  1,  1}{\textbf{MAPPED GENE}} & \textcolor[rgb]{ 1,  1,  1}{\textbf{MAPPED TRAITS}} \\
    \midrule
    \rowcolor[rgb]{ .851,  .851,  .851} PC1   & rs35399673 & 5     & 104529307 & RAB9BP1  & skin pigmentation measurement \\
    \midrule
    PC2   & rs11556924 & 7     & 129663496 & ZC3HC1 & coronary artery disease, basophil count, diastolic blood pressure, systolic blood pressure, myocardial infarction, platelet count,  platelet crit, parental longevity,  sex hormone-binding globulin measurement, testosterone measurement,  Agents acting on the renin-angiotensin system use measurement, Calcium channel blocker use measurement,  hematocrit, hemoglobin measurement, eosinophil count,  neutrophil count, myeloid white cell count, body height,  red blood cell distribution width, leukocyte count,  cardiovascular disease age at menarche \\
    \midrule
    \rowcolor[rgb]{ .851,  .851,  .851}       & rs12525051 & 6     & 151913710 & CCDC170 & heel bone mineral density \\
    \midrule
          & rs1938679 & 11    & 69272096 & MYEOV - LINC02747 & body height \\
    \midrule
    \rowcolor[rgb]{ .851,  .851,  .851}       & rs196052 & 6     & 22057200 & CASC15 & Corneal astigmatism \\
    \midrule
          & rs2069235 & 22    & 39747780 & SYNGR1 & primary biliary cirrhosis rheumatoid arthritis \\
    \midrule
    \rowcolor[rgb]{ .851,  .851,  .851}       & rs4714599 & 6     & 42285815 & TRERF1 & eosinophil percentage of granulocytes,  neutrophil percentage of granulocytes eosinophil count,  eosinophil percentage of leukocytes \\
    \midrule
          & rs5747035 & 22    & 17718606 & ADA2  & word list delayed recall measurement, memory performance \\
    \midrule
    \rowcolor[rgb]{ .851,  .851,  .851}       & rs7714191 & 5     & 131341541 & ACSL6-AS1, ACSL6 & cortical surface area measurement \\
    \midrule
          & rs7901883 & 10    & 103186838 & BTRC  & smoking behavior smoking status measurement \\
    \midrule
    \rowcolor[rgb]{ .851,  .851,  .851}       & rs7976816 & 12    & 124315343 & DNAH10 & BMI-adjusted waist circumference waist circumference \\
    \midrule
          & rs8002164 & 13    & 58248732 & PCDH17 & upper aerodigestive tract neoplasm \\
    \midrule
    \rowcolor[rgb]{ .851,  .851,  .851}       & rs847888 & 12    & 112151742 & ACAD10 & diastolic blood pressure \\
    \midrule
          & rs907183 & 8     & 8729761 & MFHAS1, MFHAS1 & Calcium channel blocker use measurement \\
    \midrule
    \rowcolor[rgb]{ .851,  .851,  .851} PC3   & rs10164546 & 2     & 106141004 & FHL2  & pursuit maintenance gain measurement \\
    \midrule
          & rs1020410 & 2     & 176784138 & EXTL2P1 - LNPK & physical activity \\
    \midrule
    \rowcolor[rgb]{ .851,  .851,  .851}       & rs10896109 & 11    & 66080023 & TMEM151A - CD248 & circadian rhythm \\
    \midrule
          & rs1264423 & 6     & 30571471 & PPP1R10 & mean corpuscular volume \\
    \midrule
    \rowcolor[rgb]{ .851,  .851,  .851}       & rs12679528 & 8     & 15566164 & TUSC3 & body mass index \\
    \midrule
          & rs16942383 & 15    & 89405052 & ACAN  & BMI-adjusted hip circumference \\
    \midrule
    \rowcolor[rgb]{ .851,  .851,  .851}       & rs2988114 & 13    & 80870878 & SPRY2 & gut microbiome measurement \\
    \midrule
          & rs34672598 & 20    & 7884260 & HAO1  & QT interval \\
    \midrule
    \rowcolor[rgb]{ .851,  .851,  .851}       & rs3828919 & 6     & 31466057 & MICB  & platelet count \\
    \midrule
          & rs41492548 & 9     & 130607359 & ENG   & monocyte count \\
    \midrule
    \rowcolor[rgb]{ .851,  .851,  .851}       & rs4679760 & 3     & 155855418 & KCNAB1 & birth weight, parental genotype effect measurement \\
    \midrule
          & rs744680 & 10    & 131741695 & EBF3  & visual perception measurement \\
    \midrule
    \rowcolor[rgb]{ .851,  .851,  .851}       & rs76496105 & 2     & 110447667 & BMS1P19 - SRSF3P6 & platelet count platelet crit \\
    \bottomrule
    \end{tabular}}%
    \caption{Traits and genes mapped in GWAS catalog from \texttt{ThreSPCA} informed variants.}
  \label{tab:traittab}%
\end{table}%

\begin{figure*}[!htb]
\begin{subfigure}{0.5\textwidth}
  \centering
  \includegraphics[width=1\linewidth]{figures/GO_pathway.png}
  \caption{Significant pathways from GO}
  \label{fig:go}
\end{subfigure}%
\begin{subfigure}{0.5\textwidth}
  \centering
  \includegraphics[width=1\linewidth]{figures/KEGG_pathway.png}
  \caption{Significant pathways from KEGG}
  \label{fig:kegg}
\end{subfigure}
\caption{GO pathway analyses of the \texttt{ThreSPCA} informed variants, colored by p-values.}
\label{fig:pathways}
\end{figure*}

\pagebreak
\subsection{Comparing \texttt{ThreSPCA} with the state-of-the-art.}

\subsubsection{Simulated data.}
We observed that increasing the threshold of true positives (markers which contribute to genetic structure) $t$ led to an increase of the number of true positives observed in \texttt{ThreSPCA}. It found l

\begin{figure*}[!htbp]
\begin{subfigure}{0.5\textwidth}
  \centering
  \includegraphics[width=1\linewidth]{figures/TP_PSD_Fst_500by5k_boxplot.png}
  \caption{True positives}
%   \label{fig:sfig5}
\end{subfigure}%
\begin{subfigure}{0.5\textwidth}
  \centering
  \includegraphics[width=1\linewidth]{figures/FP_PSD_Fst_500by5k_boxplot.png}
  \caption{False positives}
%   \label{fig:simeval}
\end{subfigure}
\caption{Box and whisker plots comparing between \texttt{ThreSPCA}, \texttt{L-PCA} and \texttt{AL-PCA} for true and false positives obtained from the simulated dataset of $m=5,000$ SNPs and $n=500$ individuals and varying $t$. }
\end{figure*}

\subsubsection{Real data.}

On 1KG data we found perfect correlation with \texttt{ThreSPCA} and \texttt{AL-PCA} for PC1 and PC2 with $r^2 = 0.97$ and 0.94 respectively. 
\begin{figure*}[!htbp]
\begin{center}
\begin{subfigure}{0.4\textwidth}
  \centering
  \includegraphics[width=1\linewidth]{figures/PC1_vs_SPC1_k500.png}
%   \caption{\texttt{ThreSPCA} with $k=500$}
%   \label{fig:sfig7}
\end{subfigure}%
\begin{subfigure}{0.4\textwidth}
  \centering
  \includegraphics[width=1\linewidth]{figures/PC2_vs_SPC2_k500.png}
%  \caption{\texttt{AL-PCA} of \cite{lee2012sparse} with $k=500$}
%   \label{fig:sfig8}
\end{subfigure}
\end{center}
%
\caption{Relationship between the \emph{PC scores} for PC1 and PC2 between \texttt{ThreSPCA} and \texttt{AL-PCA}}
\label{fig:comparison2}
\end{figure*}

\begin{figure*}[!htbp]
\begin{center}
\begin{subfigure}{0.5\textwidth}
  \centering
  \includegraphics[width=1\linewidth]{figures/SPC2_vs_SPC1_k500.png}
  \caption{\texttt{ThreSPCA} with $k=500$}
  \label{fig:sfig11}
\end{subfigure}%
\begin{subfigure}{0.5\textwidth}
  \centering
  \includegraphics[width=1\linewidth]{figures/SPC2_vs_SPC1_lee_k500.png}
  \caption{\texttt{AL-PCA} of \cite{lee2012sparse} with $k=500$}
  \label{fig:sfig6}
\end{subfigure}
\end{center}
%
\caption{PCA plots of 1KG data with \emph{PC scores}}
\label{fig:comparison}
\end{figure*}

\begin{figure*}[!htbp]
\begin{center}
\begin{subfigure}{0.45\textwidth}
  \centering
  \includegraphics[width=1\linewidth]{figures/PC2_vs_PC1.png}
  \caption{Standard PC Scores}
  \label{fig:sfig1}
\end{subfigure}%
\begin{subfigure}{0.45\textwidth}
  \centering
  \includegraphics[width=1\linewidth]{figures/SPC2_vs_SPC1_k500.png}
  \caption{Sparse PC Scores with $k=500$}
  \label{fig:sfig2}
\end{subfigure}
\\
\begin{subfigure}{0.45\textwidth}
  \centering
  \includegraphics[width=1\linewidth]{figures/SPC2_vs_SPC1_k1000.png}
  \caption{Sparse PC Scores with $k=1000$}
  \label{fig:sfig3}
\end{subfigure}
\begin{subfigure}{0.45\textwidth}
  \centering
  \includegraphics[width=1\linewidth]{figures/SPC2_vs_SPC1_k5000.png}
  \caption{Sparse PC Scores with $k=5000$}
  \label{fig:sfig4}
\end{subfigure}
\end{center}
\caption{ Scatterplots of the first two PC scores. Scatterplots of the first two PC scores from standard PCA (top left) and our \texttt{ThreSPCA} algorithm with $k=500$ (top right), $k=1000$ (bottom left)  and $k=5000$ (bottom right). Combination of different marker-shapes and marker-colors denote various population subgroups.}
\label{fig:SPCA_PCscores}
\end{figure*}

\end{appendices}


% Table generated by Excel2LaTeX from sheet 'Sheet1'
