## Supplementary material for "A Fast, Provably Accurate Approximation Algorithm for Sparse Principal Component Analysis Reveals Human Genetic Variation Across the World": zip: methods.tex

\subsection{Awesome methods}
\begin{figure}[!t]
  \begin{center}
    \includegraphics[scale=0.4]{figs/tardis.png}
  \end{center}
\caption{A simple sketch of the TARDIS.}
\label{fig:tardis}
\end{figure}
