## Supplementary material for "A Fast, Provably Accurate Approximation Algorithm for Sparse Principal Component Analysis Reveals Human Genetic Variation Across the World": zip: mybib.bib

%  a sample bibliography file
%  

@article{small,
author = {Freely, I.P.},
title = {A small paper},
journal = {The journal of small papers},
year = 1997,
volume = {-1},
note = {to appear},
}

@article{big,
author = {Jass, Hugh},
title = {A big paper},
journal = {The journal of big papers},
year = 7991,
volume = {MCMXCVII},
}

@inproceedings{dOrsiKNS20,
  author    = {Tommaso d'Orsi and
               Pravesh K. Kothari and
               Gleb Novikov and
               David Steurer},
  title     = {Sparse {PCA:} Algorithms, Adversarial Perturbations and Certificates},
  booktitle = {61st {IEEE} Annual Symposium on Foundations of Computer Science, {FOCS}},
  pages     = {553--564},
  year      = {2020}
}

@inproceedings{MuscoM15,
  author    = {Cameron Musco and
               Christopher Musco},
  title     = {Randomized Block Krylov Methods for Stronger and Faster Approximate
               Singular Value Decomposition},
  booktitle = {Advances in Neural Information Processing Systems 28: Annual Conference
               on Neural Information Processing Systems},
  pages     = {1396--1404},
  year      = {2015}
}

@article{Ali1995,
author = {Alizadeh, Farid},
title = {{Interior Point Methods in Semidefinite Programming with Applications to Combinatorial Optimization}},
journal = {SIAM Journal on Optimization},
volume = {5},
number = {1},
pages = {13-51},
year = {1995},
}

@inproceedings{DK2001,
author = {Drineas, Petros and Kannan, Ravi},
title = {{Fast Monte-Carlo Algorithms for Approximate Matrix Multiplication}},
year = {2001},
booktitle = {Proceedings of the 42nd IEEE Symposium on Foundations of Computer Science},
pages = {452-461},
}

@article{DKM2006,
author = {Drineas, Petros and Kannan, Ravi and Mahoney, Michael W.},
title = {{Fast {M}onte {C}arlo Algorithms for Matrices {II}: Computing a Low-Rank Approximation to a Matrix}},
journal = {SIAM Journal on Computing},
volume = {36},
number = {1},
pages = {158-183},
year = {2006},
}

@article{Woodruff14,
	Author = {Woodruff, David P.},
	Journal = {Foundations and Trends in Theoretical Computer Science},
	Number = {1-2},
	Title = {{Sketching as a Tool for Numerical Linear Algebra}},
	Volume = {10},
	Year = {2014}}

@Book{DM2018,
	title     = {{Lectures on Randomized Numerical Linear Algebra}},
	publisher = {American Mathematical Society},
	year      = {2018},
	author    = {Drineas, Petros and Mahoney, Michael W.},
	volume    = {25},
	series    = {The Mathematics of Data, IAS/Park City Mathematics Series},
	pages     = {1--45},
}

@article{Drineas2016,
	Author = {Drineas, Petros and Mahoney, Michael W.},
	Journal = {Communications of the ACM},
	Number = {6},
	Pages = {80--90},
	Title = {{{RandNLA}: Randomized Numerical Linear Algebra}},
	Volume = {59},
	Year = {2016}
}

@misc{Wainwright15,
  author={Wainwright, Martin J.},
  title={{Lecture Notes}},
  key={Wain15},
  note = {\url{https://www.stat.berkeley.edu/~mjwain/stat210b/Chap2_TailBounds_Jan22_2015.pdf}},
  year = {2015}
}

@article{AvronT11,
	author = {Haim Avron and 
						Sivan Toledo},
	journal = {Journal of the ACM},
	number = {2},
	pages = {8:1--8:34},
	title = {{Randomized Algorithms for Estimating the Trace of an Implicit Symmetric Positive Semi-definite Matrix}},
	volume = {58},
	year = {2011}}

@article{FountoulakisKKD17,
	author = {Kimon Fountoulakis and 
						Abhisek Kundu and 
						Eugenia{-}Maria Kontopoulou and 
						Petros Drineas},
	journal = {ACM Transactions on Knowledge Discovery from Data},
	number = {3},
	pages = {38:1--38:26},
	title = {{A Randomized Rounding Algorithm for Sparse {PCA}}},
	volume = {11},
	year = {2017}
}

@inproceedings{CohenEMMP15,
  author    = {Michael B. Cohen and
               Sam Elder and
               Cameron Musco and
               Christopher Musco and
               Madalina Persu},
  title     = {Dimensionality Reduction for k-Means Clustering and Low Rank Approximation},
  booktitle = {Proceedings of the Forty-Seventh Annual {ACM} on Symposium on Theory
               of Computing, {STOC}},
  pages     = {163--172},
  year      = {2015}
}

@article{jolliffe1995rotation,
  title={{Rotation of principal components: Choice of Normalization Constraints}},
  author={Jolliffe, Ian T.},
  journal={Journal of Applied Statistics},
  volume={22},
  number={1},
  pages={29--35},
  year={1995},
  publisher={Taylor \& Francis}
}

@article{ando2009approximating,
  title={{Approximating the Longest Path Length of a Stochastic DAG by a Normal Distribution in Linear Time}},
  author={Ando, Ei and Nakata, Toshio and Yamashita, Masafumi},
  journal={Journal of Discrete Algorithms},
  volume={7},
  number={4},
  pages={420--438},
  year={2009},
  publisher={Elsevier}
}

@article{jolliffe2003modified,
  title={{A Modified Principal Component Technique Based on the {LASSO}}},
  author={Jolliffe, Ian T. and Trendafilov, Nickolay T. and Uddin, Mudassir},
  journal={Journal of Computational and Graphical Statistics},
  volume={12},
  number={3},
  pages={531--547},
  year={2003},
  publisher={Taylor \& Francis}
}

@article{zou2006sparse,
  title={{Sparse Principal Component Analysis}},
  author={Zou, Hui and Hastie, Trevor and Tibshirani, Robert},
  journal={Journal of Computational and Graphical Statistics},
  volume={15},
  number={2},
  pages={265--286},
  year={2006},
  publisher={Taylor \& Francis}
}

@article{zou2003regression,
author = {Zou, Hui and Hastie, Trevor},
title = {{Regularization and Variable Selection via the Elastic Net}},
journal = {Journal of the Royal Statistical Society: Series B},
volume = {67},
number = {2},
pages = {301-320},
year = {2005}
}


@inproceedings{Kuleshov13,
  author    = {Volodymyr Kuleshov},
  title     = {{Fast Algorithms for Sparse Principal Component Analysis Based on {R}ayleigh
               Quotient Iteration}},
  booktitle = {Proceedings of the 30th International Conference on Machine Learning},
  pages     = {1418--1425},
  year      = {2013}
}

@article{journee2010generalized,
  title={{Generalized Power Method for Sparse Principal Component Analysis}},
  author={Journ{\'e}e, Michel and Nesterov, Yurii and Richt{\'a}rik, Peter and Sepulchre, Rodolphe},
  journal={Journal of Machine Learning Research},
  volume={11},
  number={2},
  year={2010}
}

@article{yuan2013truncated,
  title={{Truncated Power Method for Sparse Eigenvalue Problems}},
  author={Yuan, Xiao-Tong and Zhang, Tong},
  journal={Journal of Machine Learning Research},
  volume={14},
  number={Apr},
  pages={899--925},
  year={2013}
}

@article{shen2008sparse,
  title={{Sparse Principal Component Analysis via Regularized Low Rank Matrix Approximation}},
  author={Shen, Haipeng and Huang, Jianhua Z.},
  journal={Journal of Multivariate Analysis},
  volume={99},
  number={6},
  pages={1015--1034},
  year={2008},
  publisher={Elsevier}
}

@inproceedings{moghaddam2006spectral,
  title={{Spectral Bounds for Sparse {PCA}: Exact and Greedy Algorithms}},
  author={Moghaddam, Baback and Weiss, Yair and Avidan, Shai},
  booktitle={Advances in Neural Information Processing Systems},
  pages={915--922},
  year={2006}
}

@article{d2008optimal,
  title={{Optimal Solutions for Sparse Principal Component Analysis}},
  author={d’Aspremont, Alexandre and Bach, Francis and Ghaoui, Laurent El},
  journal={Journal of Machine Learning Research},
  volume={9},
  number={Jul},
  pages={1269--1294},
  year={2008}
}

@inproceedings{sriperumbudur2007sparse,
  title={{Sparse Eigen Methods by {D.C.} Programming}},
  author={Sriperumbudur, Bharath K. and Torres, David A. and Lanckriet, Gert R.G.},
  booktitle={Proceedings of the 24th International Conference on Machine Learning},
  pages={831--838},
  year={2007}
}

@article{magdon2017np,
  title={{{NP}-Hardness and Inapproximability of Sparse {PCA}}},
  author={Magdon-Ismail, Malik},
  journal={Information Processing Letters},
  volume={126},
  pages={35--38},
  year={2017},
  publisher={Elsevier}
}

@inproceedings{ChanPR16,
  author    = {Siu On Chan and
               Dimitris Papailliopoulos and
               Aviad Rubinstein},
  title     = {{On the Approximability of Sparse {PCA}}},
  booktitle = {Proceedings of the 29th Conference on Learning Theory},
  pages     = {623--646},
  year      = {2016}
}

@article{d2014approximation,
  title={{Approximation Bounds for Sparse Principal Component Analysis}},
  author={d’Aspremont, Alexandre and Bach, Francis and El Ghaoui, Laurent},
  journal={Mathematical Programming},
  volume={148},
  number={1-2},
  pages={89--110},
  year={2014},
  publisher={Springer}
}

@inproceedings{asteris2011sparse,
  title={{Sparse Principal Component of a Rank-deficient Matrix}},
  author={Asteris, Megasthenis and Papailiopoulos, Dimitris and Karystinos, George N},
  booktitle={2011 IEEE International Symposium on Information Theory Proceedings},
  pages={673--677},
  year={2011}
}

@inproceedings{asteris2015sparse,
  title={{Sparse {PCA} via Bipartite Matchings}},
  author={Asteris, Megasthenis and Papailiopoulos, Dimitris and Kyrillidis, Anastasios and Dimakis, Alexandros G},
  booktitle={Advances in Neural Information Processing Systems},
  pages={766--774},
  year={2015}
}

@article{kundu2017recovering,
  title={Recovering PCA and sparse PCA via hybrid-(l1, l2) sparse sampling of data elements},
  author={Kundu, Abhisek and Drineas, Petros and Magdon-Ismail, Malik},
  journal={Journal of Machine Learning Research},
  volume={18},
  number={1},
  pages={2558--2591},
  year={2017},
}

@article{sdp_AGJL,
	author = {d'Aspremont, Alexandre and El Ghaoui, Laurent and Jordan, Michael I. and Lanckriet, Gert R. G.},
	title = {A Direct Formulation for Sparse PCA Using Semidefinite Programming},
	year = {2007},
	issue_date = {July 2007},
	publisher = {Society for Industrial and Applied Mathematics},
	address = {USA},
	volume = {49},
	number = {3},
	issn = {0036-1445},
	url = {https://doi.org/10.1137/050645506},
	doi = {10.1137/050645506},
	journal = {SIAM Review},
	month = jul,
	pages = {434–448},
	numpages = {15}
}

@article{jeffers1967two,
	title={Two case studies in the application of principal component analysis},
	author={Jeffers, John NR},
	journal={Journal of the Royal Statistical Society: Series C (Applied Statistics)},
	volume={16},
	number={3},
	pages={225--236},
	year={1967},
	publisher={Wiley Online Library}
}

@inproceedings{fujisawa2012high,
  title={High-performance {G}eneral {S}olver for {E}xtremely {L}arge-scale {S}emidefinite {P}rogramming {P}roblems},
  author={Fujisawa, Katsuki and Sato, Hitoshi and Matsuoka, Satoshi and Endo, Toshio and Yamashita, Makoto and Nakata, Maho},
  booktitle={SC'12: Proceedings of the International Conference on High Performance Computing, Networking, Storage and Analysis},
  pages={1--11},
  year={2012},
  organization={IEEE}
}

@article{ma2013alternating,
  title={Alternating {D}irection {M}ethod of {M}ultipliers for {S}parse {P}rincipal {C}omponent {A}nalysis},
  author={Ma, Shiqian},
  journal={Journal of the Operations Research Society of China},
  volume={1},
  number={2},
  pages={253--274},
  year={2013},
  publisher={Springer}
}

@Article{Paschou2010,
   Author={Paschou, P.  and Lewis, J.  and Javed, A.  and Drineas, P. },
   Title={{A}ncestry informative markers for fine-scale individual assignment to worldwide populations},
   Journal={J. Med. Genet.},
   Year={2010},
   Volume={47},
   Number={12},
   Pages={835--847},
   Month={Dec}
}

@Article{hapmap07,
   Author={The International HapMap Consortium},
   Title={{A} second generation human haplotype map of over 3.1 Million {S}{N}{P}s},
   journal={Nature},
   Year={2007},
   Volume={449},
   Pages={851--861}
}

@Article{Li08,
   Author={Li, J. Z.  and Absher, D. M.  and Tang, H.  and Southwick, A. M.  and Casto, A. M.  and Ramachandran, S.  and Cann, H. M.  and Barsh, G. S.  and Feldman,
   M.  and Cavalli-Sforza, L. L.  and Myers, R. M.},
   Title="{{W}orldwide human relationships inferred from genome-wide patterns of variation}",
   journal={Science},
   Year={2008},
   Volume={319},
   Pages={1100--1104}
}


@inproceedings{AHK06,
 author    = {S. Arora and E. Hazan and S. Kale},
 title     = {{A Fast Random Sampling Algorithm for Sparsifying Matrices}},
 year      = {2006},
 pages     = {272-279, vol 4110},
 booktitle     = {Approximation, Randomization, and Combinatorial Optimization. Algorithms and Techniques},
 publisher = {Springer}
}

@inproceedings{PDK2013,
  title={{Sparse {PCA} through Low-rank Approximations}},
  author={Papailiopoulos, Dimitris and Dimakis, Alexandros and Korokythakis, Stavros},
  booktitle={Proceedings of the 30th International Conference on Machine Learning},
  pages={747--755},
  year={2013}
}

@inproceedings{AKL13,
 author    = {D. Achlioptas and Z. Karnin and E. Liberty},
 title     = {{Matrix entry-wise sampling: Simple is best}},
 year      = {2013},
 pages		= {},
 booktitle     = {\url{http://citeseerx.ist.psu.edu/viewdoc/download?doi=10.1.1.297.576&rep=rep1&type=pdf}}
}

@inproceedings{AKL13near,
title	={Near-Optimal Entrywise Sampling for Data Matrices},
author	={D. Achlioptas and Z. Karnin and E. Liberty},
booktitle	={Advances in Neural Information Processing Systems},
pages	={1565--1573},
year={2013}
}

@inproceedings{AM01,
 author    = {D. Achlioptas and F. McSherry},
 title		= {{Fast computation of low rank matrix approximations}},
 year      = {2001},
 pages 		= {611--618},
 booktitle     = {Proceedings of Symposium on the Theory of Computing}
}

@Article{J95,
 author    = {I. T. Jolliffe},
 title		= {{Rotation of principal components: choice of normalization constraints}},
 year      = {1995},
 pages 		= {29--35},
 journal     = {Journal of Applied Statistics}
}

@Article{CJ95,
  title={{Loading and Correlations in the Interpretation of Principal Components}},
  author={Cadima, Jorge and Jolliffe, Ian T.},
  journal={Journal of Applied Statistics},
  volume={22},
  number={2},
  pages={203--214},
  year={1995},
  publisher={Taylor \& Francis}
}

@Article{JTU2003,
 author    = {I. T. Jolliffe and N. T. Trendafilov and M. Uddin},
 title		= {{A modified principal component technique based on the lasso}},
 year      = {2003},
 pages 		= {531--547},
 journal     = {Journal of Computational and Graphical Statistics}
}

@Article{ZHT12,
 author    = {H. Zou and T. Hastie and R Tibshirani},
 title		= {{Sparse Principal Component Analysis}},
 year      = {2006},
 pages 		= {265--286},
 journal     = {Journal of Computational and Graphical Statistics}
}

@Article{AM07,
 author    = {D. Achlioptas and F. McSherry},
 title     = {{Fast computation of low-rank matrix approximations}},
 year      = {2007},
 pages     = {54(2):9},
 journal     = {Journal of the ACM}
}

@article{BCSW13,
 author    = {S Bhojanapalli and Y Chen and S Sanghavi and R Ward},
 title     = {{Coherent Matrix Completion}},
 year      = {2013},
 pages		= {},
 journal     = {\url{http://arxiv.org/pdf/1306.2979.pdf}}
}

@article{BCSW14,
 author    = {Y Chen and S Bhojanapalli and S Sanghavi and R Ward},
 title     = {{Coherent Matrix Completion}},
 year      = {2014},
 pages		= {674-682},
 journal     = {Proceedings of International Conference on Machine Learning}
}

@inproceedings{CC13,
 author    = {Y Chen and Y Chi},
 title     = {{Robust Spectral Compressed Sensing via Structured Matrix Completion}},
 year      = {2013},
 pages		= {},
 booktitle     = {Proceedings of International Conference on Machine Learning}
}

@inproceedings{CJSC11,
 author    = {Y. Chen and A. Jalali and S. Sanghavi and C. Caramanis},
 title     = {{Low-rank matrix recovery from errors and erasures}},
 year      = {2011},
 pages     = {2313-2317},
 booktitle     = {Proceedings of International Symposium on Information Theory}
}

@Article{CP10,
 author    = {E. J. Candes and Y. Plan},
 title     = {{Matrix completion with noise}},
 year      = {2010},
 pages     = {925-936, 98(6)},
 journal     = {Proceedings of the IEEE}
}

@inproceedings{CR09,
 author    = {E. J. Candes and B. Recht},
 title     = {{Exact matrix completion via convex optimization}},
 year      = {2009},
 pages     = {717-772, 9(6)},
 booktitle     = {Foundations of Computational Mathematics},
}

@inproceedings{CR12,
 author    = {E. J. Candes and B. Recht},
 title     = {{Exact matrix completion via convex optimization}},
 year      = {2012},
 pages     = {111-119, Volume 55 Issue 6},
 booktitle     = {Communications of the ACM}
}

@inproceedings{CSPW11,
 author    = {V. Chandrasekaran and S. Sanghavi and P. A. Parrilo and A. S. Willsky},
 title     = {{Rank-sparsity incoherence for matrix decomposition}},
 year      = {2011},
 pages     = {572-596, 21(2)},
 booktitle     = {SIAM Journal on Optimization}
}

@inproceedings{CT10,
 author    = {E. J. Candes and T. Tao},
 title     = {{The power of convex relaxation: Near-optimal matrix completion}},
 year      = {2010},
 pages     = {2053-2080, 56(5)},
 booktitle     = {IEEE Transactions on Information Theory}
}

@inproceedings{CLMW11,
 author    = {E. J. Candes and X. Li and Y. Ma and J. Wright},
 title     = {{Robust principal component analysis?}},
 year      = {2011},
 pages     = {58(3):11},
 booktitle     = {Journal of the ACM}
}

@inproceedings{CZ13,
 author    = {T. Tony Cai and W. Zhou},
 title     = {{Matrix Completion via Max-Norm Constrained Optimization}},
 year      = {2013},
 booktitle     = {\url{http://arxiv.org/pdf/1303.0341.pdf}}
}

@inproceedings{DKM06,
 author    = {P. Drineas and R. Kannan and M. W. Mahoney},
 title     = {{Fast Monte Carlo Algorithms for Matrices I: Approximating Matrix Multiplication}},
 year      = {2006},
 pages		= {132--157, 36(1)},
 booktitle     = {SIAM Journal on Computing}
}

@inproceedings{DZ11,
 author    = {P. Drineas and A. Zouzias},
 title     = {{A note on element-wise matrix sparsification via a matrix-valued Bernstein inequality}},
 year      = {2011},
 pages		= {385-389, 111(8)},		
 booktitle     = {Information Processing Letters}
}

@inproceedings{GM04,
 author    = {E. Gabrilovich and S. Markovitch},
 title     = {{Text categorization with many redundant features: using aggressive feature selection to make SVMs competitive with C4.5}},
 year      = {2004},
 pages		= {},
 booktitle     =  {Proceedings of International Conference on Machine Learning}
}

@inproceedings{Gross11,
 author    = {D. Gross},
 title     = {{Recovering low-rank matrices from few coefficients in any basis}},
 year      = {2011},
 pages		= {1548-1566, 57(3)},		
 booktitle     = {IEEE Transactions on Information Theory}
}

@inproceedings{JNS12,
 author    = {P. Jain and P. Netrapalli and S. Sanghavi},
 title     = {{Low-rank matrix completion using alternating minimization}},
 year      = {2012},
 pages		= {},
 booktitle     = {\url{http://arxiv.org/pdf/1212.0467}}
}

@inproceedings{KD14,
 author    = {A. Kundu and P. Drineas},
 title     = {{A Note on Randomized Element-wise Matrix Sparsification}},
 year      = {2014},
 pages		= {},
 booktitle     = {\url{http://arxiv.org/pdf/1404.0320v1.pdf}}
}

@inproceedings{Kundu15,
 author    = {A. Kundu},
 title     = {{Relaxed Leverage Sampling for Low-rank Matrix Completion}},
 year      = {2015},
 pages		= {},
 booktitle     = {\url{http://arxiv.org/pdf/1503.06379v2.pdf}}
}

@inproceedings{KMO10,
 author    = {R. H. Keshavan and A. Montanari and S. Oh},
 title     = {{Matrix completion from a few entries}},
 year      = {2010},
 pages		= {2980-2998, 56(6)},
 booktitle     = {IEEE Transactions on Information Theory}
}

@incollection{KS13,
title = {Low-Rank Matrix and Tensor Completion via Adaptive Sampling},
author = {A. Krishnamurthy and A. Singh},
booktitle = {Advances in Neural Information Processing Systems 26},
pages = {836--844},
year = {2013}
}

@inproceedings{NW12,
 author    = {S. Negahban and M. J. Wainwright},
 title     = {{Restricted strong convexity and weighted matrix completion: Optimal bounds with noise}},
 year      = {2012},
 pages		= {1665-1697, 98888},
 booktitle     = {The Journal of Machine Learning Research}
}

@inproceedings{Recht09,
 author    = {B. Recht},
 title     = {{A simpler approach to matrix completion}},
 year      = {2011},
 pages		= {3413--3430, 12},
 booktitle     = {The Journal of Machine Learning Research}
}

@inproceedings{SS10,
 author    = {R. Salakhutdinov and N. Srebro},
 title     = {{Collaborative filtering in a non-uniform world: Learning with the weighted trace norm}},
 year      = {2010},
 pages		= {},
 booktitle     = {arXiv preprint arXiv:1002.2780}
}

@inproceedings{Hull94,
 author    = {J. J. Hull},
 title     = {{A database for handwritten text recognition research}},
 year      = {1994},
 pages		= {550--554, 16(5)},
 booktitle     = {IEEE Transactions on Pattern Analysis and Machine Intelligence}
}

@inproceedings{Tropp12,
 author    = {J. Tropp},
 title     = {{User-friendly tail bounds for sums of random matrices}},
 year      = {2012},
 pages		= {12(4):389-434},
 booktitle     = {Foundations of Computational Mathematics}
}

@inproceedings{Hou10,
 author    = {J. Hou, and J. Aerts and B. den Hamer, and et al},
 title     = {{Gene Expression-Based Classification of Non-Small Cell Lung Carcinomas and Survival Prediction}},
 year      = {2010},
 pages		= {5(4):e10312},
 booktitle     = {PLoS One}
}

@inproceedings{Dracheva07,
 author    = {T. Dracheva and R. Philip and W. Xiao and AG Gee and et al},
 title     = {{Distinguishing Lung Tumours From Normal Lung Based on a Small Set of Genes}},
 year      = {2007},
 pages		= {157-164, 55(2)},
 booktitle     = {Lung Cancer}
}

@inproceedings{GM04,
 author    = {E. Gabrilovich and S. Markovitch},
 title     = {{Text categorization with many redundant features: using aggressive feature selection to make SVMs competitive with C4.5}},
 year      = {2004},
 pages		= {},
 booktitle     =  {Proceedings of International Conference on Machine Learning}
}

@inproceedings{MD09,
 author    = {Michael W. Mahoney and P. Drineas},
 title     = {{{CUR} Matrix Decompositions for Improved Data Analysis}},
 year      = {2009},
 pages 		= {697--702, 106 (3)},
 booktitle     = {Proceedings of the National Academy of Sciences}
}

@inproceedings{DMMW12a,
 author    = {P. Drineas and M. Magdon-Ismail and M. W. Mahoney and D. Woodruff},
 title     = {{Fast Approximation of Matrix Coherence and Statistical Leverage}},
 year      = {2012},
 pages 		= {},
 booktitle     = {Proceedings of International Conference on Machine Learning}
}

@inproceedings{MWA2006,
 author    = {Baback Moghaddam and Yair Weiss and Shai Avidan},
 title     = {{Generalized Spectral Bounds for Sparse {LDA}}},
 year      = {2006},
 pages 		= {641--648},
 booktitle     = {Proceedings of the 23rd International Conference on Machine learning}
}

@Article{AGJ2007,
  author    = {Alexandre d'Aspremont and
               Laurent El Ghaoui and
               Michael I. Jordan and
               Gert R. G. Lanckriet},
  title     = {{A Direct Formulation for Sparse {PCA} using Semidefinite Programming}},
  journal   = {{SIAM} Review},
  volume    = {49},
  number    = {3},
  pages     = {434--448},
  year      = {2007}
}

@Article{ABG2008,
 author    = {A. d'Aspremont and F. Bach and L. El Ghaoui},
 title     = {{Optimal Solutions for Sparse Principal Component Analysis}},
 year      = {2008},
 pages 		= {1269--1294},
 journal     = {The Journal of Machine Learning Research}
}

@Article{ABG2014,
 author    = {A. d'Aspremont and F. Bach and L. El Ghaoui},
 title     = {{Approximation Bounds for Sparse Principal Component Analysis}},
 year      = {2014},
 pages 		= {89--110},
 journal     = {Math. Program., Ser. B}
}

@inproceedings{ABG2007b,
 author    = {A. d'Aspremont and F. Bach and L. El Ghaoui},
 title     = {{Full Regularization Path for Sparse Principal Component Analysis}},
 year      = {2007},
 pages 		= {177--184},
 booktitle     = {Proceedings of the 24th international conference on Machine learning, ICML Õ07}
}

@Article{YZ2013,
 author    = {X.-T. Yuan and T. Zhang},
 title     = {{Truncated Power Method for Sparse Eigenvalue Problems}},
 year      = {2013},
 pages 		= {899--925},
 journal     = {Journal of Machine Learning Research}
}

@Article{JNRS2010,
 author    = {Michel Journ\'{e}e and Y. Nesterov and P. Richt\'{a}rik  and R. Sepulchre},
 title     = {{Generalized Power Method for Sparse Principal Component Analysis}},
 year      = {2010},
 pages 		= {517--553},
 journal     = {Journal of Machine Learning Research}
}

@Article{DMMW12b,
 author    = {P. Drineas and M. Magdon-Ismail and M. W. Mahoney and D. Woodruff},
 title     = {{Fast Approximation of Matrix Coherence and Statistical Leverage}},
 year      = {2012},
 pages 		= {3475--3506, 13},
 journal     = {Journal of Machine Learning Research}
}

@inproceedings{DMM08,
 author    = {P. Drineas and M.W. Mahoney and S. Muthukrishnan},
 title     = {{Relative-error CUR matrix decompositions}},
 year      = {2008},
 pages 		= {844--881, 30(2)},
 booktitle     = {SIAM Journal on Matrix Analysis and Applications}
}

@inproceedings{M10,
 author    = {M.W. Mahoney},
 title     = {{Randomized Algorithms for Matrices and Data}},
 year      = {2010},
 pages		= {123--224, Vol 3, Issue 2},
 booktitle     = {Foundations and Trends in Machine Learning}
}

@inproceedings{KCLE12,
author	= {K. Sj\"{o}strand and L.H. Clemmensen and R. Larsen and B. Ersb\o{}ll},
title	= {{Spasm: A matlab toolbox for sparse statistical modeling}},
year	= {2012},
 pages		= {},
booktitle	= {Journal of Statistical Software (Accepted for publication)}
}

@incollection{TMG-incoll-06,
	author = {D. Zeimpekis and E. Gallopoulos},
	title = {{TMG: A MATLAB toolbox for generating term document matrices from text collections}},
	year = {2006},
	pages = {187--210},
	publisher = {J. Kogan and C. Nicholas and M. Teboulle},
    booktitle = {Grouping Multidimensional Data: Recent Advances in Clustering}	
}

@article{Tao:2007:GTD:1313053.1313256,
 author = {D. Tao and X. Li and X. Wu and S. Maybank},
 title = {General Tensor Discriminant Analysis and Gabor Features for Gait Recognition},
 journal = {IEEE Trans. Pattern Anal. Mach. Intell.},
 volume = {29},
 number = {10},
 year = {2007},
 pages = {1700--1715},
} 

@article{plaplacian,
 author = {W. Liu and Z. Zha and Y. Wang and K. Lu and D. Tao},
 title = {$p$-Laplacian Regularized Sparse Coding for Human Activity Recognition},
 journal = {IEEE Trans. on Industrial Electronics},
 volume = {63},
 number = {8},
 year = {2016},
 pages = {5120--5129},
} 

@article{Imageclass,
 author = {Y. Luo and Y. Wen and D. Tao and J. Gui and C. Xu},
 title = {Large Margin Multi-Modal Multi-Task Feature Extraction for Image Classification},
 journal = {IEEE Trans. on Image Processing},
 volume = {25},
 number = {1},
 year = {2016},
 pages = {414-427},
}

@article{int07,
  title={{A Second Generation Human Haplotype Map of over 3.1 Million {SNP}s}},
  author={International HapMap Consortium},
  journal={Nature},
  volume={449},
  number={7164},
  pages={851},
  year={2007},
  publisher={Nature Publishing Group}
}

@article{li2008,
  title={{Worldwide Human Relationships Inferred from Genome-Wide Patterns of Variation}},
  author={Li, Jun Z. and Absher, Devin M. and Tang, Hua and Southwick, Audrey M. and Casto, Amanda M. and Ramachandran, Sohini and Cann, Howard M. and Barsh, Gregory S. and Feldman, Marcus and Cavalli-Sforza, Luigi L. and others},
  journal={Science},
  volume={319},
  number={5866},
  pages={1100--1104},
  year={2008},
  publisher={American Association for the Advancement of Science}
}

@article{landi2008gene,
  title={{Gene Expression Signature of Cigarette Smoking and Its Role in Lung Adenocarcinoma Development and Survival}},
  author={Landi, Maria Teresa and Dracheva, Tatiana and Rotunno, Melissa and Figueroa, Jonine D. and Liu, Huaitian and Dasgupta, Abhijit and Mann, Felecia E. and Fukuoka, Junya and Hames, Megan and Bergen, Andrew W. and others},
  journal={PloS one},
  volume={3},
  number={2},
  year={2008},
  publisher={Public Library of Science}
}

@article{zou2005regularization,
  title={Regularization and variable selection via the elastic net},
  author={Zou, Hui and Hastie, Trevor},
  journal={Journal of the royal statistical society: series B (statistical methodology)},
  volume={67},
  number={2},
  pages={301--320},
  year={2005},
  publisher={Wiley Online Library}
}

@incollection{zeimpekis2006tmg,
  title={{{TMG}: A {MATLAB} Toolbox for Generating Term-Document Matrices from Text Collections}},
  author={Zeimpekis, Dimitrios and Gallopoulos, Efstratios},
  booktitle={Grouping Multidimensional Data},
  pages={187--210},
  year={2006},
  publisher={Springer}
}

@article{Amini09,
    author = {Amini, Arash A. and Wainwright, Martin J.},
    title = {{High-dimensional Analysis of Semidefinite Relaxations for Sparse Principal Components}},
    journal = {Annals of Statistics},
    volume = {37},
    year = {2009},
    pages = {2877--2921}
}

@InProceedings{Yuan_2019_CVPR,
author = {Yuan, Ganzhao and Shen, Li and Zheng, Wei-Shi},
title = {A {D}ecomposition {A}lgorithm for the {S}parse {G}eneralized {E}igenvalue {P}roblem},
booktitle = {Proceedings of the IEEE/CVF Conference on Computer Vision and Pattern Recognition (CVPR)},
year = {2019}
}

@article{beck2016sparse,
  title={The {S}parse {P}rincipal {C}omponent {A}nalysis {P}roblem: {O}ptimality {C}onditions and {A}lgorithms},
  author={Beck, Amir and Vaisbourd, Yakov},
  journal={Journal of Optimization Theory and Applications},
  volume={170},
  number={1},
  pages={119--143},
  year={2016}
}

@inproceedings{papailiopoulos2013sparse,
  title={Sparse PCA through low-rank approximations},
  author={Papailiopoulos, Dimitris and Dimakis, Alexandros and Korokythakis, Stavros},
  booktitle={International Conference on Machine Learning},
  pages={747--755},
  year={2013}
}


%  The authors mentioned here are almost, but not quite,
%  entirely unrelated to Matt Groening.
