## Supplementary material for "A Fast, Provably Accurate Approximation Algorithm for Sparse Principal Component Analysis Reveals Human Genetic Variation Across the World": zip: preamble.tex

%\usepackage[protrusion=true,expansion=true]{microtype}
\usepackage{amsmath,amssymb,amsfonts}
\usepackage{latexsym}
\usepackage{subcaption}
\usepackage{graphicx}
\usepackage{booktabs}
% \usepackage[font={small,it}]{caption}
%\usepackage{fullpage}
\usepackage[backref=page]{hyperref}
\usepackage{url}
\usepackage{color}
\usepackage{wrapfig}
\usepackage{tikz}
\usetikzlibrary{decorations.pathreplacing}
\usepackage{setspace}
\usepackage{algorithm}
\usepackage[noend]{algpseudocode}
\usepackage[framemethod=tikz]{mdframed}
\usepackage{xspace}
\usepackage{pgfplots}
\usepackage{framed}
\usepackage{thmtools,thm-restate}
\usepackage{nicefrac}
\pgfplotsset{compat=1.5}
\allowdisplaybreaks

\newenvironment{proof}{\noindent{\bf Proof : \ }}{\hfill$\Box$\par\medskip}

\newenvironment{proofhack}{\noindent {\bf Proof}.\ }{}
\newtheorem{theorem}{Theorem}[section]
\newtheorem{corollary}[theorem]{Corollary}
\newtheorem{lemma}[theorem]{Lemma}
\newtheorem{proposition}[theorem]{Proposition}
\newtheorem{definition}[theorem]{Definition}
\newtheorem{remark}[theorem]{Remark}
\newtheorem{claim}[theorem]{Claim}
\newtheorem{invariant}[theorem]{Invariant}
\newtheorem{observation}[theorem]{Observation}
\newtheorem{example}[theorem]{Example}
\newtheorem{fact}[theorem]{Fact}
\newtheorem{conjecture}[theorem]{Conjecture}
\newtheorem{problem}[theorem]{Problem}

\newenvironment{proofof}[1]{\begin{trivlist} \item {\bf Proof
#1:~~}}
  {\qed\end{trivlist}}
\renewenvironment{proofof}[1]{\par\medskip\noindent{\bf Proof of #1: \ }}{\hfill$\Box$\par\medskip}
\newcommand{\namedref}[2]{\hyperref[#2]{#1~\ref*{#2}}}
%\newcommand{\namedref}[2]{#1~\ref{#2}}
\newcommand{\thmlab}[1]{\label{thm:#1}}
\newcommand{\thmref}[1]{\namedref{Theorem}{thm:#1}}
\newcommand{\lemlab}[1]{\label{lem:#1}}
\newcommand{\lemref}[1]{\namedref{Lemma}{lem:#1}}
\newcommand{\claimlab}[1]{\label{claim:#1}}
\newcommand{\claimref}[1]{\namedref{Claim}{claim:#1}}
\newcommand{\corlab}[1]{\label{cor:#1}}
\newcommand{\corref}[1]{\namedref{Corollary}{cor:#1}}
\newcommand{\seclab}[1]{\label{sec:#1}}
\newcommand{\secref}[1]{\namedref{Section}{sec:#1}}
\newcommand{\applab}[1]{\label{app:#1}}
\newcommand{\appref}[1]{\namedref{Appendix}{app:#1}}
\newcommand{\factlab}[1]{\label{fact:#1}}
\newcommand{\factref}[1]{\namedref{Fact}{fact:#1}}
\newcommand{\remlab}[1]{\label{rem:#1}}
\newcommand{\remref}[1]{\namedref{Remark}{rem:#1}}
\newcommand{\figlab}[1]{\label{fig:#1}}
\newcommand{\figref}[1]{\namedref{Figure}{fig:#1}}
\newcommand{\alglab}[1]{\label{alg:#1}}
\renewcommand{\algref}[1]{\namedref{Algorithm}{alg:#1}}
\newcommand{\tablelab}[1]{\label{tab:#1}}
\newcommand{\tableref}[1]{\namedref{Table}{tab:#1}}
\newcommand{\deflab}[1]{\label{def:#1}}
\newcommand{\defref}[1]{\namedref{Definition}{def:#1}}
\newcommand{\eqnlab}[1]{\label{eq:#1}}
\newcommand{\eqnref}[1]{\namedref{Equation}{eq:#1}}
\newcommand{\constref}[1]{\namedref{Construction}{const:#1}}
\newcommand{\constlab}[1]{\label{const:#1}}
\newcommand{\obslab}[1]{\label{obs:#1}}
\newcommand{\obsref}[1]{\namedref{Observation}{obs:#1}}
\newcommand{\exlab}[1]{\label{ex:#1}}
\newcommand{\exref}[1]{\namedref{Example}{ex:#1}}
\newcommand{\stepref}[1]{\namedref{Step}{step:#1}}
\newcommand{\steplab}[1]{\label{step:#1}}
\newcommand{\propref}[1]{\namedref{Property}{prop:#1}}
\newcommand{\proplab}[1]{\label{prop:#1}}
\newcommand{\probref}[1]{\namedref{Problem}{prob:#1}}
\newcommand{\problab}[1]{\label{prob:#1}}
\newcommand{\conjref}[1]{\namedref{Conjecture}{conj:#1}}
\newcommand{\conjlab}[1]{\label{conj:#1}}

\newenvironment{remindertheorem}[1]{\medskip \noindent {\bf Reminder of  #1.  }\em}{}


\def \HAM    {\mdef{\mathsf{HAM}}}
\renewcommand{\algorithmicrequire}{\textbf{Input:}}
\renewcommand{\algorithmicensure}{\textbf{Output:}}
\renewcommand{\algorithmiccomment}[1]{\hfill$\triangleright${\color{purple} #1}}
%---------------------------------------------------------------------------------------------------------------------------------------------------

%---------------------------------------------------------------------------------------------------------------------------------------------------

\newcommand\norm[1]{\left\lVert#1\right\rVert}
\newcommand{\PPr}[1]{\ensuremath{\mathbf{Pr}\left[#1\right]}}
\newcommand{\PPPr}[2]{\ensuremath{\underset{#1}{\mathbf{Pr}}\left[#2\right]}}
\newcommand{\Ex}[1]{\ensuremath{\mathbb{E}\left[#1\right]}}
\newcommand{\EEx}[2]{\ensuremath{\underset{#1}{\mathbb{E}}\left[#2\right]}}
\renewcommand{\O}[1]{\ensuremath{\mathcal{O}\left(#1\right)}}
\newcommand{\tO}[1]{\ensuremath{\tilde{\mathcal{O}}\left(#1\right)}}
\newcommand{\eps}{\varepsilon}
\newcommand{\nnz}{\text{nnz}}

\def \Sigmab    {\mdef{\mathbf{\Sigma}}}
\def \sparsify    {\mdef{\textsc{Sparsify}}}
\def \a    {\mdef{\mathbf{a}}}
\def \A    {\mdef{\mathbf{A}}}
\def \B    {\mdef{\mathbf{B}}}
\def \C    {\mdef{\mathbf{C}}}
\def \D    {\mdef{\mathbf{D}}}
\def \G    {\mdef{\mathbf{G}}}
\def \H    {\mdef{\mathbf{H}}}
\def \I    {\mdef{\mathbb{I}}}
\def \M    {\mdef{\mathbf{M}}}
\def \P    {\mdef{\mathbf{P}}}
\def \R    {\mdef{\mathbf{R}}}
\def \S    {\mdef{\mathbf{S}}}
\def \T    {\mdef{\mathbf{T}}}
\def \Q    {\mdef{\mathbf{Q}}}
\def \V    {\mdef{\mathbf{V}}}
\def \U    {\mdef{\mathbf{U}}}
\def \X    {\mdef{\mathbf{X}}}
\def \Y    {\mdef{\mathbf{Y}}}
\def \Z    {\mdef{\mathbf{Z}}}
\def \W    {\mdef{\mathbf{W}}}
\def \b    {\mdef{\mathbf{b}}}
\def \e    {\mdef{\mathbf{e}}}
\def \f    {\mdef{\mathbf{f}}}
\def \m    {\mdef{\mathbf{m}}}
\def \p    {\mdef{\mathbf{p}}}
\def \q    {\mdef{\mathbf{q}}}
\def \r    {\mdef{\mathbf{r}}}
\def \u    {\mdef{\mathbf{u}}}
\def \w    {\mdef{\mathbf{w}}}
\def \v    {\mdef{\mathbf{v}}}
\def \x    {\mdef{\mathbf{x}}}
\def \y    {\mdef{\mathbf{y}}}
\def \z    {\mdef{\mathbf{z}}}
\def \g    {\mdef{\mathbf{g}}}
\def \h    {\mdef{\mathbf{h}}}


\newcommand{\diag}{\mathop{\mathrm{diag}}}
\newcommand{\RR}[2]{\mathbb{R}^{#1 \times #2}}
\newcommand{\iid}{\emph{iid}}
\newcommand{\zero}{\mathbf{0}}
\newcommand{\nbr}[1]{\left\|#1\right\|}
\newcommand{\defeq}{\triangleq}
\newcommand{\Zopt}{\ensuremath{\mathcal{Z}_{\mathsf{opt}}}}
\newcommand{\xopt}{\ensuremath{\mathbf{x}_{\mathsf{opt}}}}
\newcommand{\xout}{\ensuremath{\x_{\out}}}
\newcommand{\yout}{\ensuremath{\y_{\out}}}

\renewcommand{\labelenumi}{(\arabic{enumi})}
\renewcommand\labelitemii{\mbox{$\circ$}}
\renewcommand\labelitemiii{\mbox{\tiny$\spadesuit$}}
\renewcommand{\figurename}{Fig.}

%%% Unified Formating
\newcommand{\mdef}[1]{{\ensuremath{#1}}\xspace}  % Math Def which can also be used in normal text.
\newcommand{\mydist}[1]{\mdef{\mathcal{#1}}}     % Distribution should use mathcal.
\newcommand{\myset}[1]{\mdef{\mathbb{#1}}}       % (Important) Sets should use mathbb.
\newcommand{\myalg}[1]{\mdef{\mathcal{#1}}}       % Algorithms should use mathtt.
\newcommand{\myfunc}[1]{\mdef{\mathsf{#1}}}      % Functions denoted with text (e.g. size()) should use mathsf.
\newcommand{\myvec}[1]{\mathbf{#1}}               % Vector
\def \ds {\displaystyle}                         % Shorthand for ``displaystyle''.
\DeclareMathOperator*{\argmin}{argmin}
\DeclareMathOperator*{\argmax}{argmax}
\DeclareMathOperator*{\polylog}{polylog}
\DeclareMathOperator*{\poly}{poly}
\DeclareMathOperator*{\trace}{Tr}
\DeclareMathOperator*{\rank}{rank}
\DeclareMathOperator*{\Var}{var}

%%% Text notation
\newcommand{\superscript}[1]{\ensuremath{^{\mbox{\tiny{\textit{#1}}}}}\xspace}
\def \th {\superscript{th}}     % 'The i-th entry it a list...' --> i\th
\def \st {\superscript{st}}     % 'The 1-st entry it a list...' --> 1\st
\def \nd {\superscript{nd}}     % 'The 2-nd entry it a list...' --> 2\nd
\def \rd {\superscript{rd}}     % 'The 3-rd entry it a list...' --> 3\rd
\def \etal{{\it et~al.}}

%%% Common Sets & Symbols & Functions
\def \size     {\mdef{\myfunc{size}}}                % Function outputing vector of cardinalities of a given vector of sets.
\def \negl     {\mdef{\myfunc{negl}}}                % Negliglbe function
%\def \polylog  {\mdef{\myfunc{polylog}}}             % Polylogarithm function
\def \polyloglog  {\mdef{\myfunc{polyloglog}}}       % Any function that is polynomial in the logarithm of the logarithm of the argument
\newcommand{\abs}[1]{\mdef{\left|#1\right|}}         % Absolute value
\newcommand{\flr}[1]{\mdef{\left\lfloor#1\right\rfloor}}              % Absolute value
\newcommand{\ceil}[1]{\mdef{\left\lceil#1\right\rceil}}               % Absolute value
\newcommand{\rdim}[1]{\mdef{\dim\left(#1\right)}}                     % Dimension of a vector
\newcommand{\set}[1]{\mdef{\left\{#1\right\}}}                        % Absolute value
\newcommand{\E}[2][]{\mdef{\underset{#1}{\mathbb{E}}\left[#2\right]}} % Expected value

\newcommand{\ignore}[1]{}

\newif\ifnotes\notestrue %set this to true if notes are visible and to false (next line) if they should be hidden
% \newif\ifnotes\notesfalse
\ifnotes
\newcommand{\samson}[1]{\textcolor{purple}{{\bf (Samson:} {#1}{\bf ) }} \marginpar{\tiny\bf
             \begin{minipage}[t]{0.5in}
               \raggedright S:
            \end{minipage}}}            							
\else
\newcommand{\samson}[1]{}
\fi

\definecolor{darkred}{rgb}{0.55, 0.0, 0.0}

\hypersetup{
     colorlinks   = true,
     citecolor    = mahogany,
     linkcolor	  = darkblue
}

\makeatletter
\renewcommand*{\@fnsymbol}[1]{\textcolor{darkred}{\ensuremath{\ifcase#1\or *\or \dagger\or \ddagger\or
 \mathsection\or \triangledown\or \mathparagraph\or \|\or **\or \dagger\dagger
   \or \ddagger\ddagger \else\@ctrerr\fi}}}
\makeatother

\providecommand{\email}[1]{\href{mailto:#1}{\nolinkurl{#1}\xspace}}

\definecolor{mahogany}{rgb}{0.75, 0.25, 0.0}
\definecolor{darkblue}{rgb}{0.0, 0.0, 0.55}
%\definecolor{darkpastelgreen}{rgb}{0.01, 0.75, 0.24}
%\definecolor{darkgoldenrod}{rgb}{0.72, 0.53, 0.04}
%\definecolor{darkred}{rgb}{0.55, 0.0, 0.0}
%\hypersetup{
     %colorlinks   = true,
     %citecolor    = mahogany,
		 %linkcolor		= olive
%}


%%%%%%%% Stock standard definitions %%%%%%%%%%%%%%%

\newcommand{\ab}{\mathbf{a}}
\newcommand{\bb}{\mathbf{b}}
\newcommand{\cbb}{\mathbf{c}}
\newcommand{\db}{\mathbf{d}}
\newcommand{\eb}{\mathbf{e}}
\newcommand{\fb}{\mathbf{f}}
\newcommand{\gb}{\mathbf{g}}
\newcommand{\hb}{\mathbf{h}}
\newcommand{\ib}{\mathbf{i}}
\newcommand{\jb}{\mathbf{j}}
\newcommand{\kb}{\mathbf{k}}
\newcommand{\lb}{\mathbf{l}}
\newcommand{\mb}{\mathbf{m}}
\newcommand{\nbb}{\mathbf{n}}
\newcommand{\ob}{\mathbf{o}}
\newcommand{\pb}{\mathbf{p}}
\newcommand{\qb}{\mathbf{q}}
\newcommand{\rb}{\mathbf{r}}
\newcommand{\sbb}{\mathbf{s}}
\newcommand{\tb}{\mathbf{t}}
\newcommand{\ub}{\mathbf{u}}
\newcommand{\vb}{\mathbf{v}}
\newcommand{\wb}{\mathbf{w}}
\newcommand{\xb}{\mathbf{x}}
\newcommand{\yb}{\mathbf{y}}
\newcommand{\zb}{\mathbf{z}}

\newcommand{\abtil}{\tilde{\ab}}
\newcommand{\bbtil}{\tilde{\bb}}
\newcommand{\cbtil}{\tilde{\cbb}}
\newcommand{\dbtil}{\tilde{\db}}
\newcommand{\ebtil}{\tilde{\eb}}
\newcommand{\fbtil}{\tilde{\fb}}
\newcommand{\gbtil}{\tilde{\gb}}
\newcommand{\hbtil}{\tilde{\hb}}
\newcommand{\ibtil}{\tilde{\ib}}
\newcommand{\jbtil}{\tilde{\jb}}
\newcommand{\kbtil}{\tilde{\kb}}
\newcommand{\lbtil}{\tilde{\lb}}
\newcommand{\mbtil}{\tilde{\mb}}
\newcommand{\nbtil}{\tilde{\nbb}}
\newcommand{\obtil}{\tilde{\ob}}
\newcommand{\pbtil}{\tilde{\pb}}
\newcommand{\qbtil}{\tilde{\qb}}
\newcommand{\rbtil}{\tilde{\rb}}
\newcommand{\sbtil}{\tilde{\sbb}}
\newcommand{\tbtil}{\tilde{\tb}}
\newcommand{\ubtil}{\tilde{\ub}}
\newcommand{\vbtil}{\tilde{\vb}}
\newcommand{\wbtil}{\tilde{\wb}}
\newcommand{\xbtil}{\tilde{\xb}}
\newcommand{\ybtil}{\tilde{\yb}}
\newcommand{\zbtil}{\tilde{\zb}}

\newcommand{\atil}{\tilde{a}}
\newcommand{\btil}{\tilde{b}}
\newcommand{\ctil}{\tilde{c}}
\newcommand{\dtil}{\tilde{d}}
\newcommand{\etil}{\tilde{e}}
\newcommand{\ftil}{\tilde{f}}
\newcommand{\gtil}{\tilde{g}}
\newcommand{\htil}{\tilde{h}}
\newcommand{\itil}{\tilde{i}}
\newcommand{\jtil}{\tilde{j}}
\newcommand{\ktil}{\tilde{k}}
\newcommand{\ltil}{\tilde{l}}
\newcommand{\mtil}{\tilde{m}}
\newcommand{\ntil}{\tilde{n}}
\newcommand{\otil}{\tilde{o}}
\newcommand{\ptil}{\tilde{p}}
\newcommand{\qtil}{\tilde{q}}
\newcommand{\rtil}{\tilde{r}}
\newcommand{\stil}{\tilde{s}}
\newcommand{\ttil}{\tilde{t}}
\newcommand{\util}{\tilde{u}}
\newcommand{\vtil}{\tilde{v}}
\newcommand{\wtil}{\tilde{w}}
\newcommand{\xtil}{\tilde{x}}
\newcommand{\ytil}{\tilde{y}}
\newcommand{\ztil}{\tilde{z}}

\newcommand{\ahat}{\hat{a}}
\newcommand{\bhat}{\hat{b}}
\newcommand{\chat}{\hat{c}}
\newcommand{\dhat}{\hat{d}}
\newcommand{\ehat}{\hat{e}}
\newcommand{\fhat}{\hat{f}}
\newcommand{\ghat}{\hat{g}}
\newcommand{\hhat}{\hat{h}}
\newcommand{\ihat}{\hat{i}}
\newcommand{\jhat}{\hat{j}}
\newcommand{\khat}{\hat{k}}
\newcommand{\lhat}{\hat{l}}
\newcommand{\mhat}{\hat{m}}
\newcommand{\nhat}{\hat{n}}
\newcommand{\ohat}{\hat{o}}
\newcommand{\phat}{\hat{p}}
\newcommand{\qhat}{\hat{q}}
\newcommand{\rhat}{\hat{r}}
\newcommand{\shat}{\hat{s}}
\newcommand{\that}{\hat{t}}
\newcommand{\uhat}{\hat{u}}
\newcommand{\vhat}{\hat{v}}
\newcommand{\what}{\hat{w}}
\newcommand{\xhat}{\hat{x}}
\newcommand{\yhat}{\hat{y}}
\newcommand{\zhat}{\hat{z}}

\newcommand{\abhat}{\hat{\ab}}
\newcommand{\bbhat}{\hat{\bb}}
\newcommand{\cbhat}{\hat{\cbb}}
\newcommand{\dbhat}{\hat{\db}}
\newcommand{\ebhat}{\hat{\eb}}
\newcommand{\fbhat}{\hat{\fb}}
\newcommand{\gbhat}{\hat{\gb}}
\newcommand{\hbhat}{\hat{\hb}}
\newcommand{\ibhat}{\hat{\ib}}
\newcommand{\jbhat}{\hat{\jb}}
\newcommand{\kbhat}{\hat{\kb}}
\newcommand{\lbhat}{\hat{\lb}}
\newcommand{\mbhat}{\hat{\mb}}
\newcommand{\nbhat}{\hat{\nb}}
\newcommand{\obhat}{\hat{\ob}}
\newcommand{\pbhat}{\hat{\pb}}
\newcommand{\qbhat}{\hat{\qb}}
\newcommand{\rbhat}{\hat{\rb}}
\newcommand{\sbhat}{\hat{\sb}}
\newcommand{\tbhat}{\hat{\tb}}
\newcommand{\ubhat}{\hat{\ub}}
\newcommand{\vbhat}{\hat{\vb}}
\newcommand{\wbhat}{\hat{\wb}}
\newcommand{\xbhat}{\hat{\xb}}
\newcommand{\ybhat}{\hat{\yb}}
\newcommand{\zbhat}{\hat{\zb}}

\newcommand{\Atil}{\tilde{A}}
\newcommand{\Btil}{\tilde{B}}
\newcommand{\Ctil}{\tilde{C}}
\newcommand{\Dtil}{\tilde{D}}
\newcommand{\Etil}{\tilde{E}}
\newcommand{\Ftil}{\tilde{F}}
\newcommand{\Gtil}{\tilde{G}}
\newcommand{\Htil}{\tilde{H}}
\newcommand{\Itil}{\tilde{I}}
\newcommand{\Jtil}{\tilde{J}}
\newcommand{\Ktil}{\tilde{K}}
\newcommand{\Ltil}{\tilde{L}}
\newcommand{\Mtil}{\tilde{M}}
\newcommand{\Ntil}{\tilde{N}}
\newcommand{\Otil}{\tilde{O}}
\newcommand{\Ptil}{\tilde{P}}
\newcommand{\Qtil}{\tilde{Q}}
\newcommand{\Rtil}{\tilde{R}}
\newcommand{\Stil}{\tilde{S}}
\newcommand{\Ttil}{\tilde{T}}
\newcommand{\Util}{\tilde{U}}
\newcommand{\Vtil}{\tilde{V}}
\newcommand{\Wtil}{\tilde{W}}
\newcommand{\Xtil}{\tilde{X}}
\newcommand{\Ytil}{\tilde{Y}}
\newcommand{\Ztil}{\tilde{Z}}

\newcommand{\abar}{\bar{a}}
\newcommand{\bbar}{\bar{b}}
\newcommand{\cbar}{\bar{c}}
\newcommand{\dbar}{\bar{d}}
\newcommand{\ebar}{\bar{e}}
\newcommand{\fbar}{\bar{f}}
\newcommand{\gbar}{\bar{g}}
\newcommand{\hbr}{\bar{h}}
\newcommand{\ibar}{\bar{i}}
\newcommand{\jbar}{\bar{j}}
\newcommand{\kbar}{\bar{k}}
\newcommand{\lbar}{\bar{l}}
\newcommand{\mbar}{\bar{m}}
\newcommand{\nbar}{\bar{n}}
\newcommand{\obar}{\bar{o}}
\newcommand{\pbar}{\bar{p}}
\newcommand{\qbar}{\bar{q}}
\newcommand{\rbar}{\bar{r}}
\newcommand{\sbar}{\bar{s}}
\newcommand{\tbar}{\bar{t}}
\newcommand{\ubar}{\bar{u}}
\newcommand{\vbar}{\bar{v}}
\newcommand{\wbar}{\bar{w}}
\newcommand{\xbar}{\bar{x}}
\newcommand{\ybar}{\bar{y}}
\newcommand{\zbar}{\bar{z}}

\newcommand{\abbar}{\bar{\ab}}
\newcommand{\bbbar}{\bar{\bb}}
\newcommand{\cbbar}{\bar{\cb}}
\newcommand{\dbbar}{\bar{\db}}
\newcommand{\ebbar}{\bar{\eb}}
\newcommand{\fbbar}{\bar{\fb}}
\newcommand{\gbbar}{\bar{\gb}}
\newcommand{\hbbar}{\bar{\hb}}
\newcommand{\ibbar}{\bar{\ib}}
\newcommand{\jbbar}{\bar{\jb}}
\newcommand{\kbbar}{\bar{\kb}}
\newcommand{\lbbar}{\bar{\lb}}
\newcommand{\mbbar}{\bar{\mb}}
\newcommand{\nbbar}{\bar{\nbb}}
\newcommand{\obbar}{\bar{\ob}}
\newcommand{\pbbar}{\bar{\pb}}
\newcommand{\qbbar}{\bar{\qb}}
\newcommand{\rbbar}{\bar{\rb}}
\newcommand{\sbbar}{\bar{\sbb}}
\newcommand{\tbbar}{\bar{\tb}}
\newcommand{\ubbar}{\bar{\ub}}
\newcommand{\vbbar}{\bar{\vb}}
\newcommand{\wbbar}{\bar{\wb}}
\newcommand{\xbbar}{\bar{\xb}}
\newcommand{\ybbar}{\bar{\yb}}
\newcommand{\zbbar}{\bar{\zb}}

\newcommand{\Ab}{\mathbf{A}}
\newcommand{\Bb}{\mathbf{B}}
\newcommand{\Cb}{\mathbf{C}}
\newcommand{\Db}{\mathbf{D}}
\newcommand{\Eb}{\mathbf{E}}
\newcommand{\Fb}{\mathbf{F}}
\newcommand{\Gb}{\mathbf{G}}
\newcommand{\Hb}{\mathbf{H}}
\newcommand{\Ib}{\mathbf{I}}
\newcommand{\Jb}{\mathbf{J}}
\newcommand{\Kb}{\mathbf{K}}
\newcommand{\Lb}{\mathbf{L}}
\newcommand{\Mb}{\mathbf{M}}
\newcommand{\Nb}{\mathbf{N}}
\newcommand{\Ob}{\mathbf{O}}
\newcommand{\Pb}{\mathbf{P}}
\newcommand{\Qb}{\mathbf{Q}}
\newcommand{\Rb}{\mathbf{R}}
\newcommand{\Sb}{\mathbf{S}}
\newcommand{\Tb}{\mathbf{T}}
\newcommand{\Ub}{\mathbf{U}}
\newcommand{\Vb}{\mathbf{V}}
\newcommand{\Wb}{\mathbf{W}}
\newcommand{\Xb}{\mathbf{X}}
\newcommand{\Yb}{\mathbf{Y}}
\newcommand{\Zb}{\mathbf{Z}}

% \newcommand{\Abtil}{\tilde{\Ab}}
\newcommand{\Abtil}{\widetilde{\Ab}}
\newcommand{\Bbtil}{\tilde{\Bb}}
\newcommand{\Cbtil}{\tilde{\Cb}}
\newcommand{\Dbtil}{\tilde{\Db}}
\newcommand{\Ebtil}{\tilde{\Eb}}
\newcommand{\Fbtil}{\tilde{\Fb}}
\newcommand{\Gbtil}{\tilde{\Gb}}
\newcommand{\Hbtil}{\tilde{\Hb}}
\newcommand{\Ibtil}{\tilde{\Ib}}
\newcommand{\Jbtil}{\tilde{\Jb}}
\newcommand{\Kbtil}{\tilde{\Kb}}
\newcommand{\Lbtil}{\tilde{\Lb}}
\newcommand{\Mbtil}{\tilde{\Mb}}
\newcommand{\Nbtil}{\tilde{\Nb}}
\newcommand{\Obtil}{\tilde{\Ob}}
\newcommand{\Pbtil}{\tilde{\Pb}}
\newcommand{\Qbtil}{\tilde{\Qb}}
\newcommand{\Rbtil}{\tilde{\Rb}}
\newcommand{\Sbtil}{\tilde{\Sb}}
\newcommand{\Tbtil}{\tilde{\Tb}}
\newcommand{\Ubtil}{\tilde{\Ub}}
\newcommand{\Vbtil}{\tilde{\Vb}}
\newcommand{\Wbtil}{\tilde{\Wb}}
\newcommand{\Xbtil}{\tilde{\Xb}}
\newcommand{\Ybtil}{\tilde{\Yb}}
\newcommand{\Zbtil}{\tilde{\Zb}}

\newcommand{\Abar}{\bar{A}}
\newcommand{\Bbar}{\bar{B}}
\newcommand{\Cbar}{\bar{C}}
\newcommand{\Dbar}{\bar{D}}
\newcommand{\Ebar}{\bar{E}}
\newcommand{\Fbar}{\bar{F}}
\newcommand{\Gbar}{\bar{G}}
\newcommand{\Hbar}{\bar{H}}
\newcommand{\Ibar}{\bar{I}}
\newcommand{\Jbar}{\bar{J}}
\newcommand{\Kbar}{\bar{K}}
\newcommand{\Lbar}{\bar{L}}
\newcommand{\Mbar}{\bar{M}}
\newcommand{\Nbar}{\bar{N}}
\newcommand{\Obar}{\bar{O}}
\newcommand{\Pbar}{\bar{P}}
\newcommand{\Qbar}{\bar{Q}}
\newcommand{\Rbar}{\bar{R}}
\newcommand{\Sbar}{\bar{S}}
\newcommand{\Tbar}{\bar{T}}
\newcommand{\Ubar}{\bar{U}}
\newcommand{\Vbar}{\bar{V}}
\newcommand{\Wbar}{\bar{W}}
\newcommand{\Xbar}{\bar{X}}
\newcommand{\Ybar}{\bar{Y}}
\newcommand{\Zbar}{\bar{Z}}

\newcommand{\Abbar}{\bar{\Ab}}
\newcommand{\Bbbar}{\bar{\Bb}}
\newcommand{\Cbbar}{\bar{\Cb}}
\newcommand{\Dbbar}{\bar{\Db}}
\newcommand{\Ebbar}{\bar{\Eb}}
\newcommand{\Fbbar}{\bar{\Fb}}
\newcommand{\Gbbar}{\bar{\Gb}}
\newcommand{\Hbbar}{\bar{\Hb}}
\newcommand{\Ibbar}{\bar{\Ib}}
\newcommand{\Jbbar}{\bar{\Jb}}
\newcommand{\Kbbar}{\bar{\Kb}}
\newcommand{\Lbbar}{\bar{\Lb}}
\newcommand{\Mbbar}{\bar{\Mb}}
\newcommand{\Nbbar}{\bar{\Nb}}
\newcommand{\Obbar}{\bar{\Ob}}
\newcommand{\Pbbar}{\bar{\Pb}}
\newcommand{\Qbbar}{\bar{\Qb}}
\newcommand{\Rbbar}{\bar{\Rb}}
\newcommand{\Sbbar}{\bar{\Sb}}
\newcommand{\Tbbar}{\bar{\Tb}}
\newcommand{\Ubbar}{\bar{\Ub}}
\newcommand{\Vbbar}{\bar{\Vb}}
\newcommand{\Wbbar}{\bar{\Wb}}
\newcommand{\Xbbar}{\bar{\Xb}}
\newcommand{\Ybbar}{\bar{\Yb}}
\newcommand{\Zbbar}{\bar{\Zb}}

\newcommand{\Ahat}{\hat{A}}
\newcommand{\Bhat}{\hat{B}}
\newcommand{\Chat}{\hat{C}}
\newcommand{\Dhat}{\hat{D}}
\newcommand{\Ehat}{\hat{E}}
\newcommand{\Fhat}{\hat{F}}
\newcommand{\Ghat}{\hat{G}}
\newcommand{\Hhat}{\hat{H}}
\newcommand{\Ihat}{\hat{I}}
\newcommand{\Jhat}{\hat{J}}
\newcommand{\Khat}{\hat{K}}
\newcommand{\Lhat}{\hat{L}}
\newcommand{\Mhat}{\hat{M}}
\newcommand{\Nhat}{\hat{N}}
\newcommand{\Ohat}{\hat{O}}
\newcommand{\Phat}{\hat{P}}
\newcommand{\Qhat}{\hat{Q}}
\newcommand{\Rhat}{\hat{R}}
\newcommand{\Shat}{\hat{S}}
\newcommand{\That}{\hat{T}}
\newcommand{\Uhat}{\hat{U}}
\newcommand{\Vhat}{\hat{V}}
\newcommand{\What}{\hat{W}}
\newcommand{\Xhat}{\hat{X}}
\newcommand{\Yhat}{\hat{Y}}
\newcommand{\Zhat}{\hat{Z}}

\newcommand{\Abhat}{\hat{\Ab}}
\newcommand{\Bbhat}{\hat{\Bb}}
\newcommand{\Cbhat}{\hat{\Cb}}
\newcommand{\Dbhat}{\hat{\Db}}
\newcommand{\Ebhat}{\hat{\Eb}}
\newcommand{\Fbhat}{\hat{\Fb}}
\newcommand{\Gbhat}{\hat{\Gb}}
\newcommand{\Hbhat}{\hat{\Hb}}
\newcommand{\Ibhat}{\hat{\Ib}}
\newcommand{\Jbhat}{\hat{\Jb}}
\newcommand{\Kbhat}{\hat{\Kb}}
\newcommand{\Lbhat}{\hat{\Lb}}
\newcommand{\Mbhat}{\hat{\Mb}}
\newcommand{\Nbhat}{\hat{\Nb}}
\newcommand{\Obhat}{\hat{\Ob}}
\newcommand{\Pbhat}{\hat{\Pb}}
\newcommand{\Qbhat}{\hat{\Qb}}
\newcommand{\Rbhat}{\hat{\Rb}}
\newcommand{\Sbhat}{\hat{\Sb}}
\newcommand{\Tbhat}{\hat{\Tb}}
\newcommand{\Ubhat}{\hat{\Ub}}
\newcommand{\Vbhat}{\hat{\Vb}}
\newcommand{\Wbhat}{\hat{\Wb}}
\newcommand{\Xbhat}{\hat{\Xb}}
\newcommand{\Ybhat}{\hat{\Yb}}
\newcommand{\Zbhat}{\hat{\Zb}}

\newcommand{\Acal}{\mathcal{A}}
\newcommand{\Bcal}{\mathcal{B}}
\newcommand{\Ccal}{\mathcal{C}}
\newcommand{\Dcal}{\mathcal{D}}
\newcommand{\Ecal}{\mathcal{E}}
\newcommand{\Fcal}{\mathcal{F}}
\newcommand{\Gcal}{\mathcal{G}}
\newcommand{\Hcal}{\mathcal{H}}
\newcommand{\Ical}{\mathcal{I}}
\newcommand{\Jcal}{\mathcal{J}}
\newcommand{\Kcal}{\mathcal{K}}
\newcommand{\Lcal}{\mathcal{L}}
\newcommand{\Mcal}{\mathcal{M}}
\newcommand{\Ncal}{\mathcal{N}}
\newcommand{\Ocal}{\mathcal{O}}
\newcommand{\Pcal}{\mathcal{P}}
\newcommand{\Qcal}{\mathcal{Q}}
\newcommand{\Rcal}{\mathcal{R}}
\newcommand{\Scal}{\mathcal{S}}
\newcommand{\Tcal}{\mathcal{T}}
\newcommand{\Ucal}{\mathcal{U}}
\newcommand{\Vcal}{\mathcal{V}}
\newcommand{\Wcal}{\mathcal{W}}
\newcommand{\Xcal}{\mathcal{X}}
\newcommand{\Ycal}{\mathcal{Y}}
\newcommand{\Zcal}{\mathcal{Z}}

\newcommand{\Ascr}{\mathscr{A}}
\newcommand{\Bscr}{\mathscr{B}}
\newcommand{\Cscr}{\mathscr{C}}
\newcommand{\Dscr}{\mathscr{D}}
\newcommand{\Escr}{\mathscr{E}}
\newcommand{\Fscr}{\mathscr{F}}
\newcommand{\Gscr}{\mathscr{G}}
\newcommand{\Hscr}{\mathscr{H}}
\newcommand{\Iscr}{\mathscr{I}}
\newcommand{\Jscr}{\mathscr{J}}
\newcommand{\Kscr}{\mathscr{K}}
\newcommand{\Lscr}{\mathscr{L}}
\newcommand{\Mscr}{\mathscr{M}}
\newcommand{\Nscr}{\mathscr{N}}
\newcommand{\Oscr}{\mathscr{O}}
\newcommand{\Pscr}{\mathscr{P}}
\newcommand{\Qscr}{\mathscr{Q}}
\newcommand{\Rscr}{\mathscr{R}}
\newcommand{\Sscr}{\mathscr{S}}
\newcommand{\Tscr}{\mathscr{T}}
\newcommand{\Uscr}{\mathscr{U}}
\newcommand{\Vscr}{\mathscr{V}}
\newcommand{\Wscr}{\mathscr{W}}
\newcommand{\Xscr}{\mathscr{X}}
\newcommand{\Yscr}{\mathscr{Y}}
\newcommand{\Zscr}{\mathscr{Z}}

\newcommand{\Ascrb}{\boldsymbol{\Ascr}}
\newcommand{\Bscrb}{\boldsymbol{\Bscr}}
\newcommand{\Cscrb}{\boldsymbol{\Cscr}}
\newcommand{\Dscrb}{\boldsymbol{\Dscr}}
\newcommand{\Escrb}{\boldsymbol{\Escr}}
\newcommand{\Fscrb}{\boldsymbol{\Fscr}}
\newcommand{\Gscrb}{\boldsymbol{\Gscr}}
\newcommand{\Hscrb}{\boldsymbol{\Hscr}}
\newcommand{\Iscrb}{\boldsymbol{\Iscr}}
\newcommand{\Jscrb}{\boldsymbol{\Jscr}}
\newcommand{\Kscrb}{\boldsymbol{\Kscr}}
\newcommand{\Lscrb}{\boldsymbol{\Lscr}}
\newcommand{\Mscrb}{\boldsymbol{\Mscr}}
\newcommand{\Nscrb}{\boldsymbol{\Nscr}}
\newcommand{\Oscrb}{\boldsymbol{\Oscr}}
\newcommand{\Pscrb}{\boldsymbol{\Pscr}}
\newcommand{\Qscrb}{\boldsymbol{\Qscr}}
\newcommand{\Rscrb}{\boldsymbol{\Rscr}}
\newcommand{\Sscrb}{\boldsymbol{\Sscr}}
\newcommand{\Tscrb}{\boldsymbol{\Tscr}}
\newcommand{\Uscrb}{\boldsymbol{\Uscr}}
\newcommand{\Vscrb}{\boldsymbol{\Vscr}}
\newcommand{\Wscrb}{\boldsymbol{\Wscr}}
\newcommand{\Xscrb}{\boldsymbol{\Xscr}}
\newcommand{\Xscrbh}{\hat{\Xscrb}}
\newcommand{\Yscrb}{\boldsymbol{\Yscr}}
\newcommand{\Zscrb}{\boldsymbol{\Zscr}}

\newcommand{\Atilde}{\widetilde{A}}
\newcommand{\Btilde}{\widetilde{B}}
\newcommand{\Ctilde}{\widetilde{C}}
\newcommand{\Dtilde}{\widetilde{D}}
\newcommand{\Etilde}{\widetilde{E}}
\newcommand{\Ftilde}{\widetilde{F}}
\newcommand{\Gtilde}{\widetilde{G}}
\newcommand{\Htilde}{\widetilde{H}}
\newcommand{\Itilde}{\widetilde{I}}
\newcommand{\Jtilde}{\widetilde{J}}
\newcommand{\Ktilde}{\widetilde{K}}
\newcommand{\Ltilde}{\widetilde{L}}
\newcommand{\Mtilde}{\widetilde{M}}
\newcommand{\Ntilde}{\widetilde{N}}
\newcommand{\Otilde}{\widetilde{O}}
\newcommand{\Ptilde}{\widetilde{P}}
\newcommand{\Qtilde}{\widetilde{Q}}
\newcommand{\Rtilde}{\widetilde{R}}
\newcommand{\Stilde}{\widetilde{S}}
\newcommand{\Ttilde}{\widetilde{T}}
\newcommand{\Utilde}{\widetilde{U}}
\newcommand{\Vtilde}{\widetilde{V}}
\newcommand{\Wtilde}{\widetilde{W}}
\newcommand{\Xtilde}{\widetilde{X}}
\newcommand{\Ytilde}{\widetilde{Y}}
\newcommand{\Ztilde}{\widetilde{Z}}

\renewcommand{\vec}[1]{\mathbf{\boldsymbol{#1}}}

\newcommand{\avec}{\vec{a}}
\newcommand{\bvec}{\vec{b}}
\newcommand{\cvec}{\vec{c}}
\newcommand{\dvec}{\vec{d}}
\newcommand{\evec}{\vec{e}}
\newcommand{\fvec}{\vec{f}}
\newcommand{\gvec}{\vec{g}}
\newcommand{\hvec}{\vec{h}}
\newcommand{\ivec}{\vec{i}}
\newcommand{\jvec}{\vec{j}}
\newcommand{\kvec}{\vec{k}}
\newcommand{\lvec}{\vec{l}}
\newcommand{\mvec}{\vec{m}}
\newcommand{\nvec}{\vec{n}}
\newcommand{\ovec}{\vec{o}}
\newcommand{\pvec}{\vec{p}}
\newcommand{\qvec}{\vec{q}}
\newcommand{\rvec}{\vec{r}}
\newcommand{\svec}{\vec{s}}
\newcommand{\tvec}{\vec{t}}
\newcommand{\uvec}{\vec{u}}
\newcommand{\vvec}{\vec{v}}
\newcommand{\wvec}{\vec{w}}
\newcommand{\xvec}{\vec{x}}
\newcommand{\yvec}{\vec{y}}
\newcommand{\zvec}{\vec{z}}

\newcommand{\EE}{\mathbb{E}} % Expectation
\newcommand{\txb}{\tilde{\mathbf{x}}} 
