## Supplementary material for "A Fast, Provably Accurate Approximation Algorithm for Sparse Principal Component Analysis Reveals Human Genetic Variation Across the World": zip: references.bib

@inproceedings{dOrsiKNS20,
  author    = {Tommaso d'Orsi and
               Pravesh K. Kothari and
               Gleb Novikov and
               David Steurer},
  title     = {Sparse {PCA:} Algorithms, Adversarial Perturbations and Certificates},
  booktitle = {61st {IEEE} Annual Symposium on Foundations of Computer Science, {FOCS}},
  pages     = {553--564},
  year      = {2020}
}

@inproceedings{MuscoM15,
  author    = {Cameron Musco and
               Christopher Musco},
  title     = {Randomized Block Krylov Methods for Stronger and Faster Approximate
               Singular Value Decomposition},
  booktitle = {Advances in Neural Information Processing Systems 28: Annual Conference
               on Neural Information Processing Systems},
  pages     = {1396--1404},
  year      = {2015}
}

@article{Ali1995,
author = {Alizadeh, Farid},
title = {{Interior Point Methods in Semidefinite Programming with Applications to Combinatorial Optimization}},
journal = {SIAM Journal on Optimization},
volume = {5},
number = {1},
pages = {13-51},
year = {1995},
}

@inproceedings{DK2001,
author = {Drineas, Petros and Kannan, Ravi},
title = {{Fast Monte-Carlo Algorithms for Approximate Matrix Multiplication}},
year = {2001},
booktitle = {Proceedings of the 42nd IEEE Symposium on Foundations of Computer Science},
pages = {452-461},
}

@article{DKM2006,
author = {Drineas, Petros and Kannan, Ravi and Mahoney, Michael W.},
title = {{Fast {M}onte {C}arlo Algorithms for Matrices {II}: Computing a Low-Rank Approximation to a Matrix}},
journal = {SIAM Journal on Computing},
volume = {36},
number = {1},
pages = {158-183},
year = {2006},
}

@article{Woodruff14,
	Author = {Woodruff, David P.},
	Journal = {Foundations and Trends in Theoretical Computer Science},
	Number = {1-2},
	Title = {{Sketching as a Tool for Numerical Linear Algebra}},
	Volume = {10},
	Year = {2014}}

@Book{DM2018,
	title     = {{Lectures on Randomized Numerical Linear Algebra}},
	publisher = {American Mathematical Society},
	year      = {2018},
	author    = {Drineas, Petros and Mahoney, Michael W.},
	volume    = {25},
	series    = {The Mathematics of Data, IAS/Park City Mathematics Series},
	pages     = {1--45},
}

@article{Drineas2016,
	Author = {Drineas, Petros and Mahoney, Michael W.},
	Journal = {Communications of the ACM},
	Number = {6},
	Pages = {80--90},
	Title = {{{RandNLA}: Randomized Numerical Linear Algebra}},
	Volume = {59},
	Year = {2016}
}

@misc{Wainwright15,
  author={Wainwright, Martin J.},
  title={{Lecture Notes}},
  key={Wain15},
  note = {\url{https://www.stat.berkeley.edu/~mjwain/stat210b/Chap2_TailBounds_Jan22_2015.pdf}},
  year = {2015}
}

@article{AvronT11,
	author = {Haim Avron and 
						Sivan Toledo},
	journal = {Journal of the ACM},
	number = {2},
	pages = {8:1--8:34},
	title = {{Randomized Algorithms for Estimating the Trace of an Implicit Symmetric Positive Semi-definite Matrix}},
	volume = {58},
	year = {2011}}

@article{FountoulakisKKD17,
	author = {Kimon Fountoulakis and 
						Abhisek Kundu and 
						Eugenia{-}Maria Kontopoulou and 
						Petros Drineas},
	journal = {ACM Transactions on Knowledge Discovery from Data},
	number = {3},
	pages = {38:1--38:26},
	title = {{A Randomized Rounding Algorithm for Sparse {PCA}}},
	volume = {11},
	year = {2017}
}

@inproceedings{CohenEMMP15,
  author    = {Michael B. Cohen and
               Sam Elder and
               Cameron Musco and
               Christopher Musco and
               Madalina Persu},
  title     = {Dimensionality Reduction for k-Means Clustering and Low Rank Approximation},
  booktitle = {Proceedings of the Forty-Seventh Annual {ACM} on Symposium on Theory
               of Computing, {STOC}},
  pages     = {163--172},
  year      = {2015}
}

@article{jolliffe1995rotation,
  title={{Rotation of principal components: Choice of Normalization Constraints}},
  author={Jolliffe, Ian T.},
  journal={Journal of Applied Statistics},
  volume={22},
  number={1},
  pages={29--35},
  year={1995},
  publisher={Taylor \& Francis}
}

@article{ando2009approximating,
  title={{Approximating the Longest Path Length of a Stochastic DAG by a Normal Distribution in Linear Time}},
  author={Ando, Ei and Nakata, Toshio and Yamashita, Masafumi},
  journal={Journal of Discrete Algorithms},
  volume={7},
  number={4},
  pages={420--438},
  year={2009},
  publisher={Elsevier}
}

@article{jolliffe2003modified,
  title={{A Modified Principal Component Technique Based on the {LASSO}}},
  author={Jolliffe, Ian T. and Trendafilov, Nickolay T. and Uddin, Mudassir},
  journal={Journal of Computational and Graphical Statistics},
  volume={12},
  number={3},
  pages={531--547},
  year={2003},
  publisher={Taylor \& Francis}
}

@article{zou2006sparse,
  title={{Sparse Principal Component Analysis}},
  author={Zou, Hui and Hastie, Trevor and Tibshirani, Robert},
  journal={Journal of Computational and Graphical Statistics},
  volume={15},
  number={2},
  pages={265--286},
  year={2006},
  publisher={Taylor \& Francis}
}

@article{zou2003regression,
author = {Zou, Hui and Hastie, Trevor},
title = {{Regularization and Variable Selection via the Elastic Net}},
journal = {Journal of the Royal Statistical Society: Series B},
volume = {67},
number = {2},
pages = {301-320},
year = {2005}
}


@inproceedings{Kuleshov13,
  author    = {Volodymyr Kuleshov},
  title     = {{Fast Algorithms for Sparse Principal Component Analysis Based on {R}ayleigh
               Quotient Iteration}},
  booktitle = {Proceedings of the 30th International Conference on Machine Learning},
  pages     = {1418--1425},
  year      = {2013}
}

@article{journee2010generalized,
  title={{Generalized Power Method for Sparse Principal Component Analysis}},
  author={Journ{\'e}e, Michel and Nesterov, Yurii and Richt{\'a}rik, Peter and Sepulchre, Rodolphe},
  journal={Journal of Machine Learning Research},
  volume={11},
  number={2},
  year={2010}
}

@article{yuan2013truncated,
  title={{Truncated Power Method for Sparse Eigenvalue Problems}},
  author={Yuan, Xiao-Tong and Zhang, Tong},
  journal={Journal of Machine Learning Research},
  volume={14},
  number={Apr},
  pages={899--925},
  year={2013}
}

@article{shen2008sparse,
  title={{Sparse Principal Component Analysis via Regularized Low Rank Matrix Approximation}},
  author={Shen, Haipeng and Huang, Jianhua Z.},
  journal={Journal of Multivariate Analysis},
  volume={99},
  number={6},
  pages={1015--1034},
  year={2008},
  publisher={Elsevier}
}

@inproceedings{moghaddam2006spectral,
  title={{Spectral Bounds for Sparse {PCA}: Exact and Greedy Algorithms}},
  author={Moghaddam, Baback and Weiss, Yair and Avidan, Shai},
  booktitle={Advances in Neural Information Processing Systems},
  pages={915--922},
  year={2006}
}

@article{d2008optimal,
  title={{Optimal Solutions for Sparse Principal Component Analysis}},
  author={d’Aspremont, Alexandre and Bach, Francis and Ghaoui, Laurent El},
  journal={Journal of Machine Learning Research},
  volume={9},
  number={Jul},
  pages={1269--1294},
  year={2008}
}

@inproceedings{sriperumbudur2007sparse,
  title={{Sparse Eigen Methods by {D.C.} Programming}},
  author={Sriperumbudur, Bharath K. and Torres, David A. and Lanckriet, Gert R.G.},
  booktitle={Proceedings of the 24th International Conference on Machine Learning},
  pages={831--838},
  year={2007}
}

@article{magdon2017np,
  title={{{NP}-Hardness and Inapproximability of Sparse {PCA}}},
  author={Magdon-Ismail, Malik},
  journal={Information Processing Letters},
  volume={126},
  pages={35--38},
  year={2017},
  publisher={Elsevier}
}

@inproceedings{ChanPR16,
  author    = {Siu On Chan and
               Dimitris Papailliopoulos and
               Aviad Rubinstein},
  title     = {{On the Approximability of Sparse {PCA}}},
  booktitle = {Proceedings of the 29th Conference on Learning Theory},
  pages     = {623--646},
  year      = {2016}
}

@article{d2014approximation,
  title={{Approximation Bounds for Sparse Principal Component Analysis}},
  author={d’Aspremont, Alexandre and Bach, Francis and El Ghaoui, Laurent},
  journal={Mathematical Programming},
  volume={148},
  number={1-2},
  pages={89--110},
  year={2014},
  publisher={Springer}
}

@inproceedings{asteris2011sparse,
  title={{Sparse Principal Component of a Rank-deficient Matrix}},
  author={Asteris, Megasthenis and Papailiopoulos, Dimitris and Karystinos, George N},
  booktitle={2011 IEEE International Symposium on Information Theory Proceedings},
  pages={673--677},
  year={2011}
}

@inproceedings{asteris2015sparse,
  title={{Sparse {PCA} via Bipartite Matchings}},
  author={Asteris, Megasthenis and Papailiopoulos, Dimitris and Kyrillidis, Anastasios and Dimakis, Alexandros G},
  booktitle={Advances in Neural Information Processing Systems},
  pages={766--774},
  year={2015}
}

@article{kundu2017recovering,
  title={Recovering PCA and sparse PCA via hybrid-(l1, l2) sparse sampling of data elements},
  author={Kundu, Abhisek and Drineas, Petros and Magdon-Ismail, Malik},
  journal={Journal of Machine Learning Research},
  volume={18},
  number={1},
  pages={2558--2591},
  year={2017},
}

@article{sdp_AGJL,
	author = {d'Aspremont, Alexandre and El Ghaoui, Laurent and Jordan, Michael I. and Lanckriet, Gert R. G.},
	title = {A Direct Formulation for Sparse PCA Using Semidefinite Programming},
	year = {2007},
	issue_date = {July 2007},
	publisher = {Society for Industrial and Applied Mathematics},
	address = {USA},
	volume = {49},
	number = {3},
	issn = {0036-1445},
	url = {https://doi.org/10.1137/050645506},
	doi = {10.1137/050645506},
	journal = {SIAM Review},
	month = jul,
	pages = {434–448},
	numpages = {15}
}

@article{jeffers1967two,
	title={Two case studies in the application of principal component analysis},
	author={Jeffers, John NR},
	journal={Journal of the Royal Statistical Society: Series C (Applied Statistics)},
	volume={16},
	number={3},
	pages={225--236},
	year={1967},
	publisher={Wiley Online Library}
}

@inproceedings{fujisawa2012high,
  title={High-performance {G}eneral {S}olver for {E}xtremely {L}arge-scale {S}emidefinite {P}rogramming {P}roblems},
  author={Fujisawa, Katsuki and Sato, Hitoshi and Matsuoka, Satoshi and Endo, Toshio and Yamashita, Makoto and Nakata, Maho},
  booktitle={SC'12: Proceedings of the International Conference on High Performance Computing, Networking, Storage and Analysis},
  pages={1--11},
  year={2012},
  organization={IEEE}
}

@article{ma2013alternating,
  title={Alternating {D}irection {M}ethod of {M}ultipliers for {S}parse {P}rincipal {C}omponent {A}nalysis},
  author={Ma, Shiqian},
  journal={Journal of the Operations Research Society of China},
  volume={1},
  number={2},
  pages={253--274},
  year={2013},
  publisher={Springer}
}

@article{lee2012sparse,
  title={Sparse {P}rincipal {C}omponent {A}nalysis for {I}dentifying {A}ncestry-informative {M}arkers in {G}enome-wide {A}ssociation {S}tudies},
  author={Lee, Seokho and Epstein, Michael P and Duncan, Richard and Lin, Xihong},
  journal={Genetic Epidemiology},
  volume={36},
  number={4},
  pages={293--302},
  year={2012}
}

@article{leng2009general,
  title={On general adaptive sparse principal component analysis},
  author={Leng, Chenlei and Wang, Hansheng},
  journal={Journal of Computational and Graphical Statistics},
  volume={18},
  number={1},
  pages={201--215},
  year={2009}}
}

@article{10002015global,
  title={A global reference for human genetic variation},
  author={1000 Genomes Project Consortium and others},
  journal={Nature},
  volume={526},
  number={7571},
  pages={68},
  year={2015},
  publisher={NIH Public Access}
}


@article{mclaren2016ensembl,
  title={The ensembl variant effect predictor},
  author={McLaren, William and Gil, Laurent and Hunt, Sarah E and Riat, Harpreet Singh and Ritchie, Graham RS and Thormann, Anja and Flicek, Paul and Cunningham, Fiona},
  journal={Genome biology},
  volume={17},
  number={1},
  pages={1--14},
  year={2016},
  publisher={BioMed Central}
}

@article{buniello2019nhgri,
  title={The NHGRI-EBI GWAS Catalog of published genome-wide association studies, targeted arrays and summary statistics 2019},
  author={Buniello, Annalisa and MacArthur, Jacqueline A L and Cerezo, Maria and Harris, Laura W and Hayhurst, James and Malangone, Cinzia and McMahon, Aoife and Morales, Joannella and Mountjoy, Edward and Sollis, Elliot and others},
  journal={Nucleic acids research},
  volume={47},
  number={D1},
  pages={D1005--D1012},
  year={2019},
  publisher={Oxford University Press}
}
@article{ng2003sift,
  title={SIFT: Predicting amino acid changes that affect protein function},
  author={Ng, Pauline C and Henikoff, Steven},
  journal={Nucleic acids research},
  volume={31},
  number={13},
  pages={3812--3814},
  year={2003},
  publisher={Oxford University Press}
}
@article{sunyaev2001prediction,
  title={Prediction of deleterious human alleles},
  author={Sunyaev, Shamil and Ramensky, Vasily and Koch, Ina and Lathe III, Warren and Kondrashov, Alexey S and Bork, Peer},
  journal={Human molecular genetics},
  volume={10},
  number={6},
  pages={591--597},
  year={2001},
  publisher={Oxford University Press}
}
@article{volgman2018atherosclerotic,
  title={Atherosclerotic cardiovascular disease in South Asians in the United States: epidemiology, risk factors, and treatments: a scientific statement from the American Heart Association},
  author={Volgman, Annabelle Santos and Palaniappan, Latha S and Aggarwal, Neelum T and Gupta, Milan and Khandelwal, Abha and Krishnan, Aruna V and Lichtman, Judith H and Mehta, Laxmi S and Patel, Hena N and Shah, Kevin S and others},
  journal={Circulation},
  volume={138},
  number={1},
  pages={e1--e34},
  year={2018},
  publisher={Am Heart Assoc}
}
@article{bose2020clustrat,
  title={CluStrat: a structure informed clustering strategy for population stratification},
  author={Bose, Aritra and Burch, Myson C and Chowdhury, Agniva and Paschou, Peristera and Drineas, Petros},
  journal={bioRxiv},
  year={2020},
  publisher={Cold Spring Harbor Laboratory}
}
@article{pritchard2000inference,
  title={Inference of population structure using multilocus genotype data},
  author={Pritchard, Jonathan K and Stephens, Matthew and Donnelly, Peter},
  journal={Genetics},
  volume={155},
  number={2},
  pages={945--959},
  year={2000},
  publisher={Oxford University Press}
}
@article{patterson2006population,
  title={Population structure and eigenanalysis},
  author={Patterson, Nick and Price, Alkes L and Reich, David},
  journal={PLoS genetics},
  volume={2},
  number={12},
  pages={e190},
  year={2006},
  publisher={Public Library of Science San Francisco, USA}
}
@article{price2006principal,
  title={Principal components analysis corrects for stratification in genome-wide association studies},
  author={Price, Alkes L and Patterson, Nick J and Plenge, Robert M and Weinblatt, Michael E and Shadick, Nancy A and Reich, David},
  journal={Nature genetics},
  volume={38},
  number={8},
  pages={904--909},
  year={2006},
  publisher={Nature Publishing Group}
}
@article{engelhardt2010analysis,
  title={Analysis of population structure: a unifying framework and novel methods based on sparse factor analysis},
  author={Engelhardt, Barbara E and Stephens, Matthew},
  journal={PLoS genetics},
  volume={6},
  number={9},
  pages={e1001117},
  year={2010},
  publisher={Public Library of Science San Francisco, USA}
}
@article{hsu2014sparse,
  title={Sparse principal component analysis in cancer research},
  author={Hsu, Ying-Lin and Huang, Po-Yu and Chen, Dung-Tsa},
  journal={Translational cancer research},
  volume={3},
  number={3},
  pages={182},
  year={2014},
  publisher={NIH Public Access}
}
@article{chang2015second,
  title={Second-generation PLINK: rising to the challenge of larger and richer datasets},
  author={Chang, Christopher C and Chow, Carson C and Tellier, Laurent CAM and Vattikuti, Shashaank and Purcell, Shaun M and Lee, James J},
  journal={Gigascience},
  volume={4},
  number={1},
  pages={s13742--015},
  year={2015},
  publisher={Oxford University Press}
}
@article{bose2019terapca,
  title={TeraPCA: a fast and scalable software package to study genetic variation in tera-scale genotypes},
  author={Bose, Aritra and Kalantzis, Vassilis and Kontopoulou, Eugenia-Maria and Elkady, Mai and Paschou, Peristera and Drineas, Petros},
  journal={Bioinformatics},
  volume={35},
  number={19},
  pages={3679--3683},
  year={2019},
  publisher={Oxford University Press}
}
@article{sohail2019polygenic,
  title={Polygenic adaptation on height is overestimated due to uncorrected stratification in genome-wide association studies},
  author={Sohail, Mashaal and Maier, Robert M and Ganna, Andrea and Bloemendal, Alex and Martin, Alicia R and Turchin, Michael C and Chiang, Charleston WK and Hirschhorn, Joel and Daly, Mark J and Patterson, Nick and others},
  journal={Elife},
  volume={8},
  pages={e39702},
  year={2019},
  publisher={eLife Sciences Publications Limited}
}
@article{yu2012clusterprofiler,
  title={clusterProfiler: an R package for comparing biological themes among gene clusters},
  author={Yu, Guangchuang and Wang, Li-Gen and Han, Yanyan and He, Qing-Yu},
  journal={Omics: a journal of integrative biology},
  volume={16},
  number={5},
  pages={284--287},
  year={2012},
  publisher={Mary Ann Liebert, Inc. 140 Huguenot Street, 3rd Floor New Rochelle, NY 10801 USA}
}
