## Supplementary figures and images for "A Fast, Provably Accurate Approximation Algorithm for Sparse Principal Component Analysis Reveals Human Genetic Variation Across the World"

### FP_PSD_Fst_1kby10k_boxplot.png

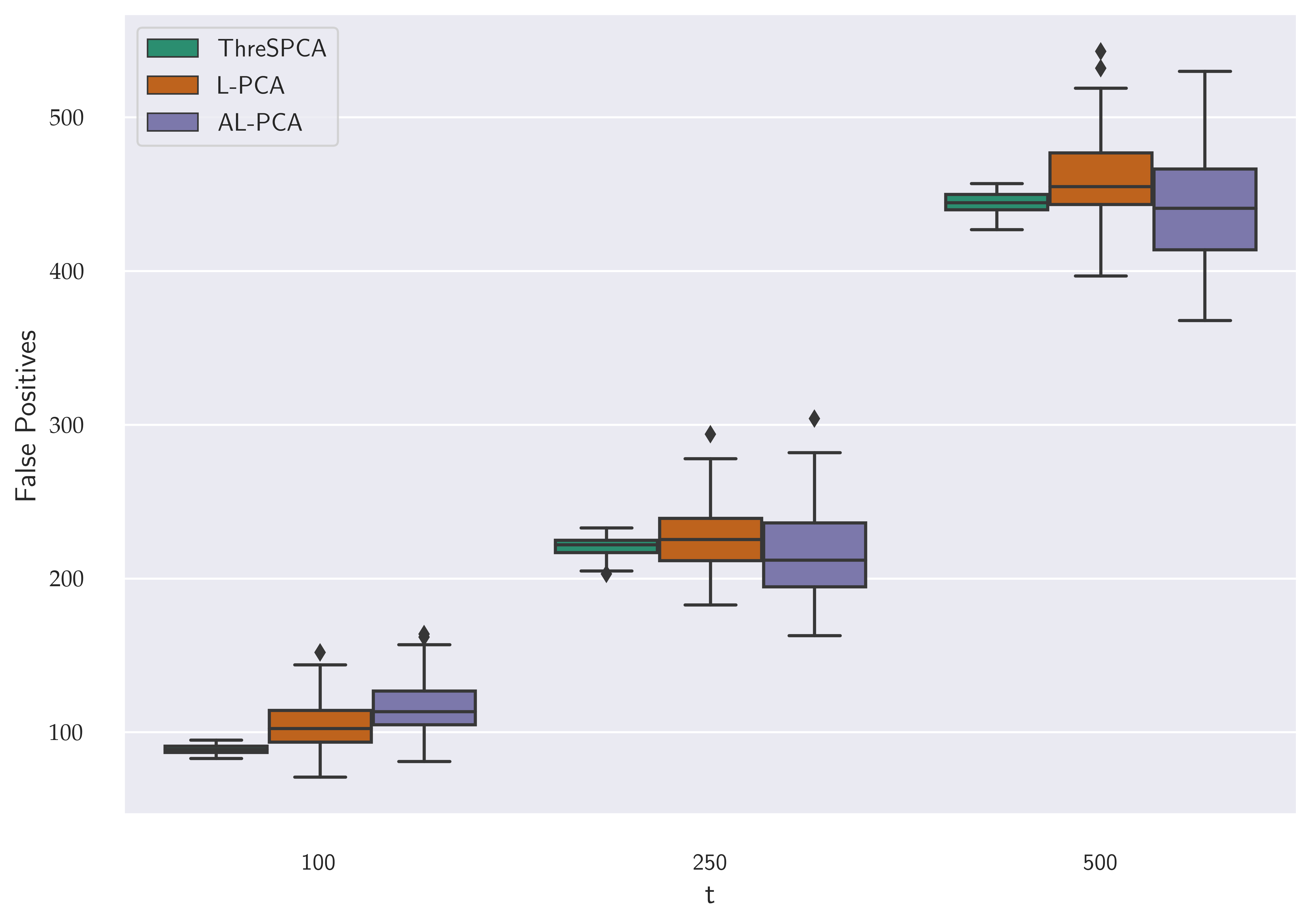

### FP_PSD_Fst_500by5k_boxplot.png

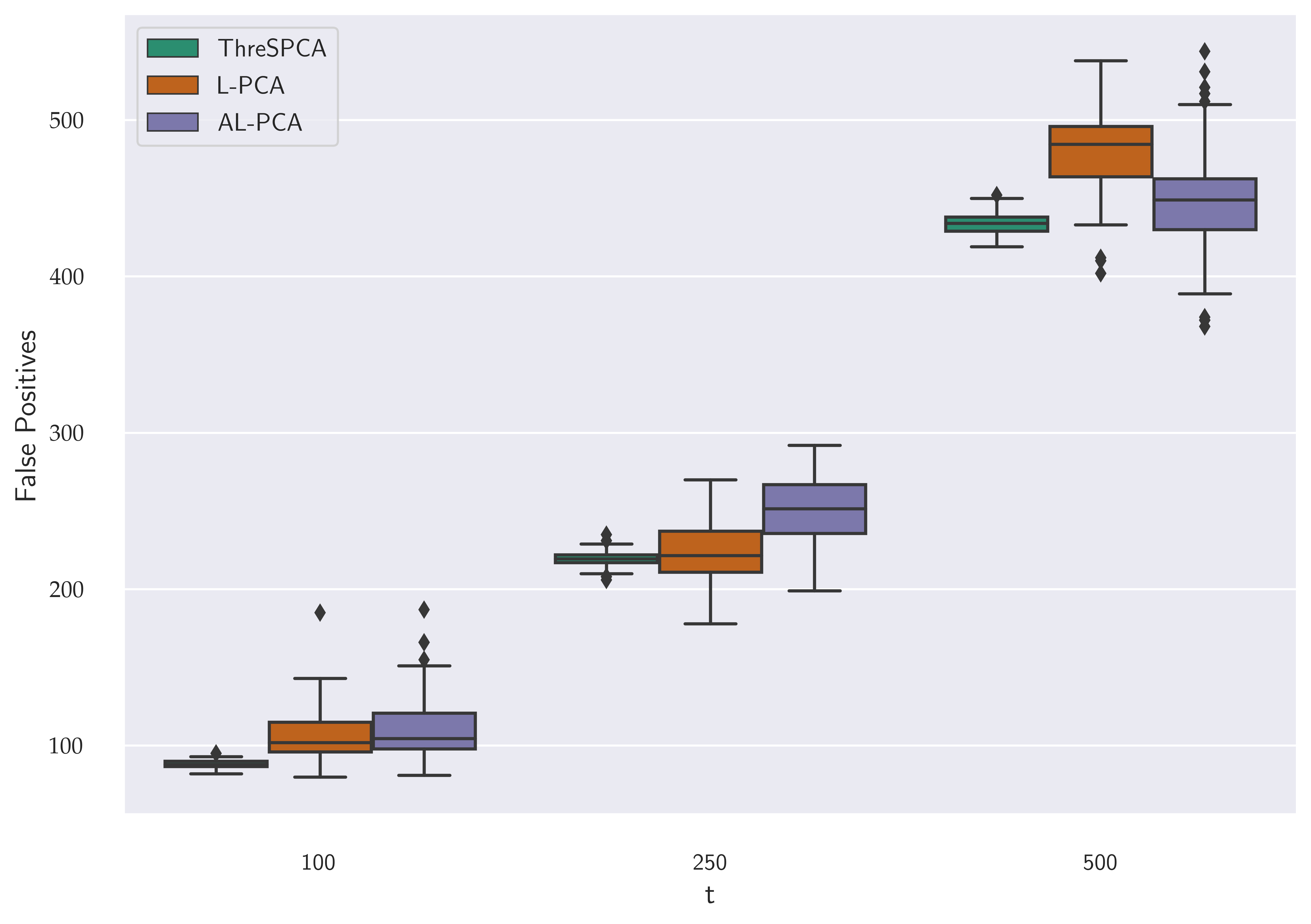

### GO_pathway.png

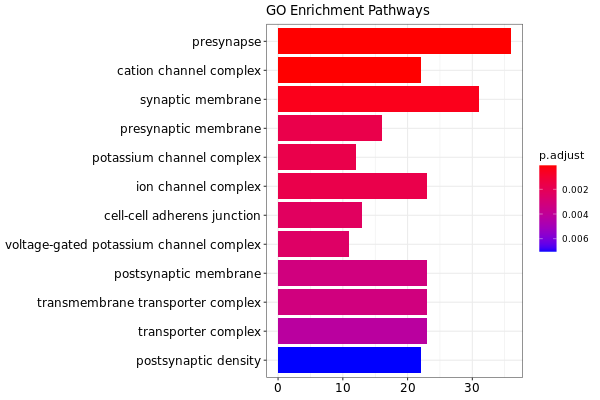

### k_by_r2.png

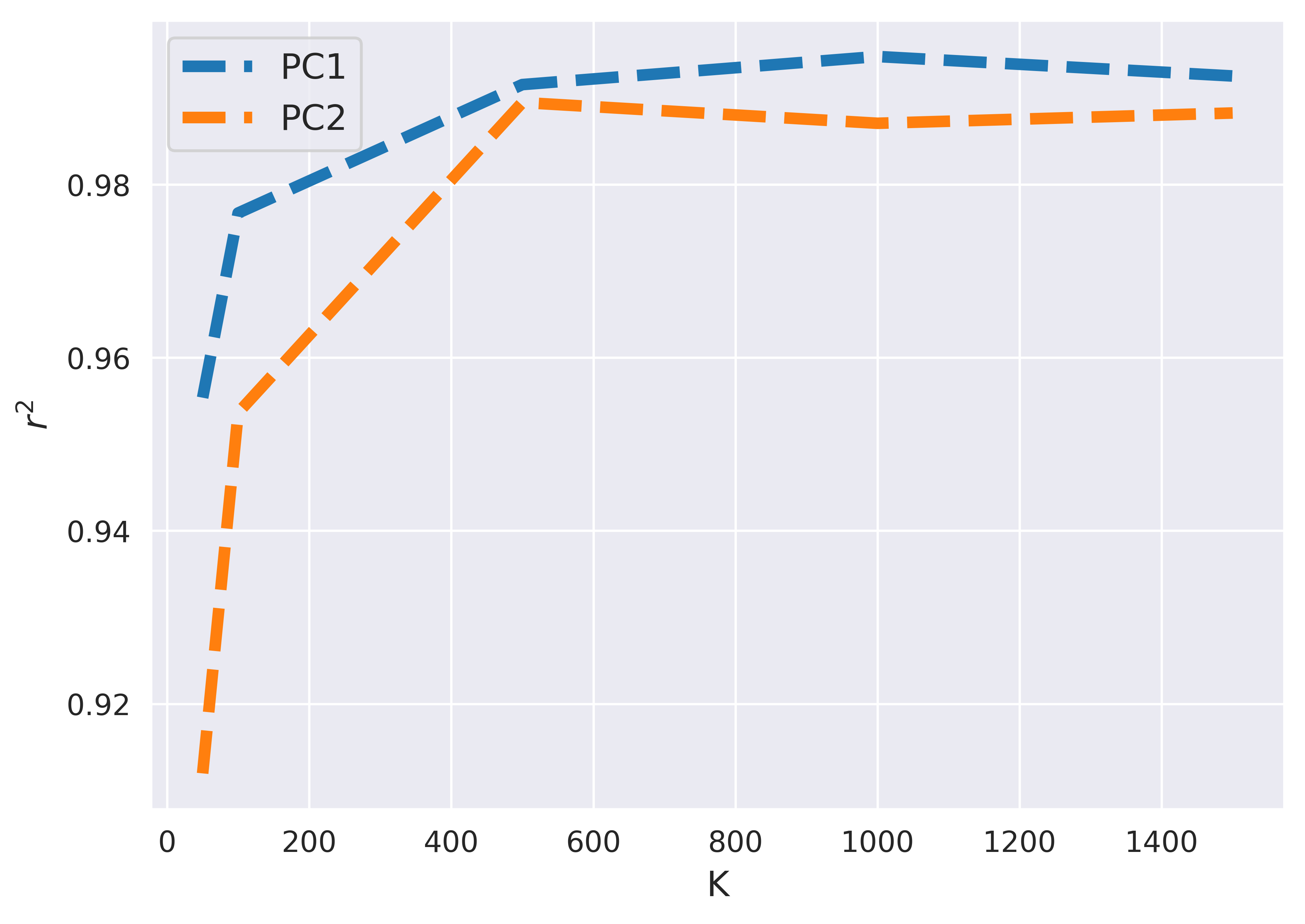

### KEGG_pathway.png

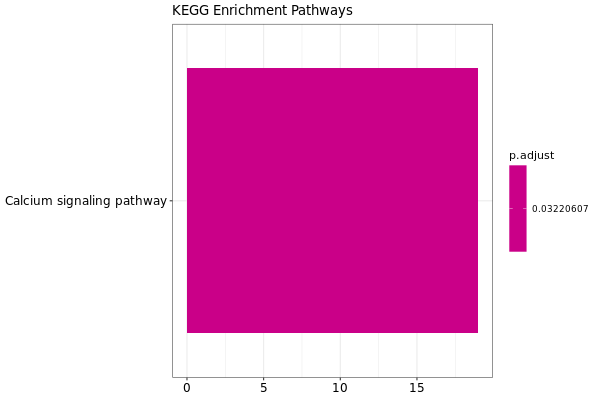

### PC1_vs_SPC1_k500.png

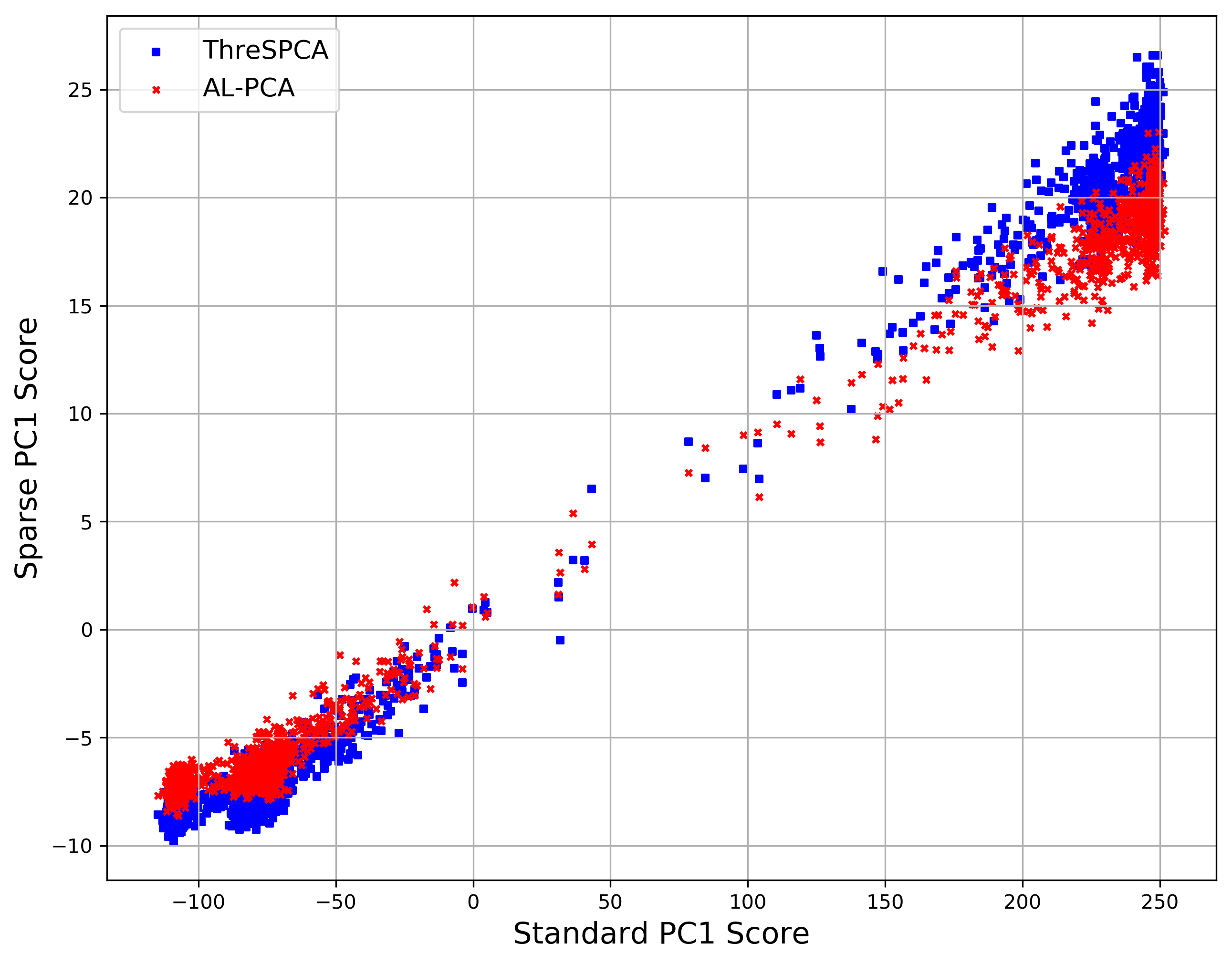

### PC2_vs_PC1.png

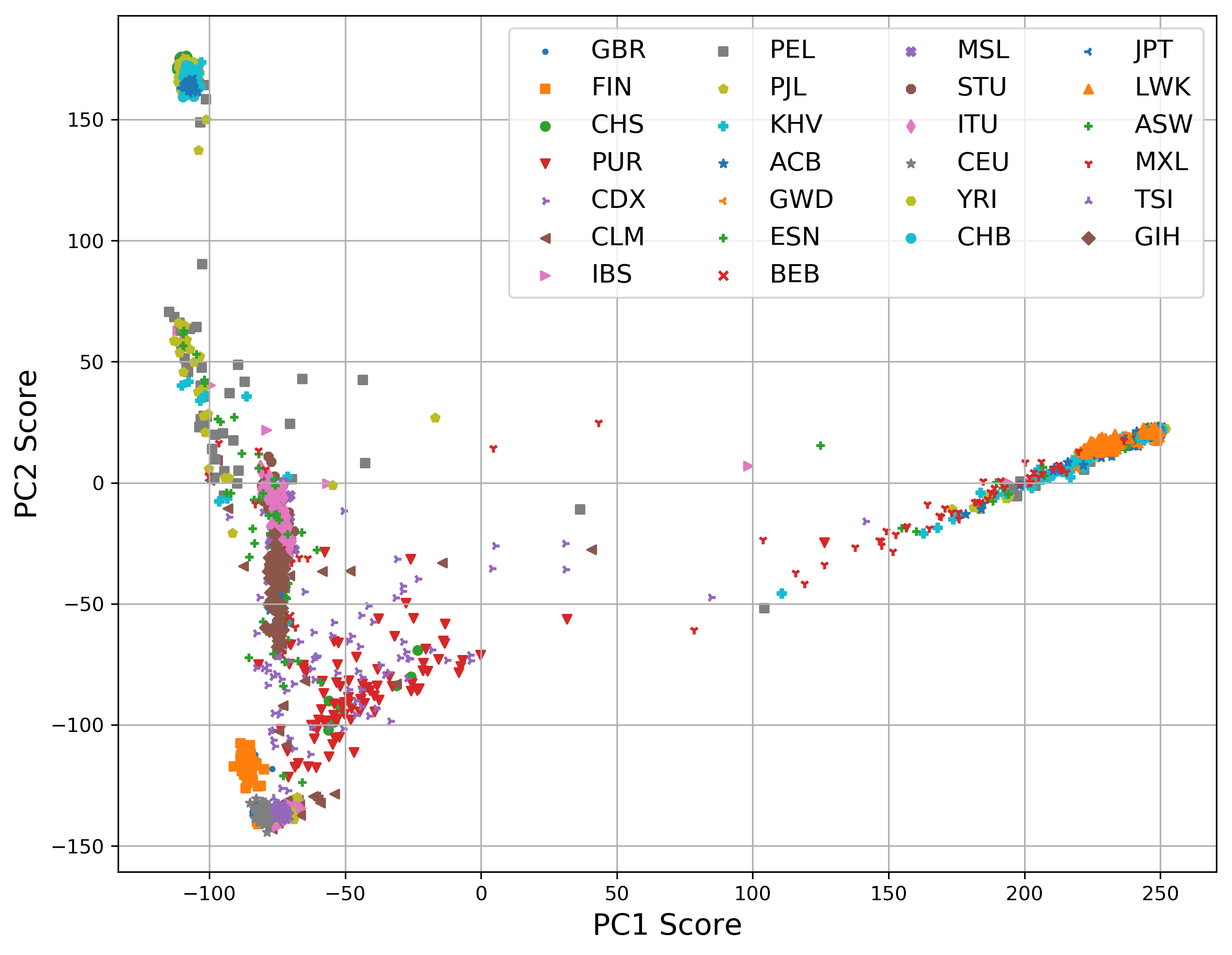

### PC2_vs_SPC2_k500.png

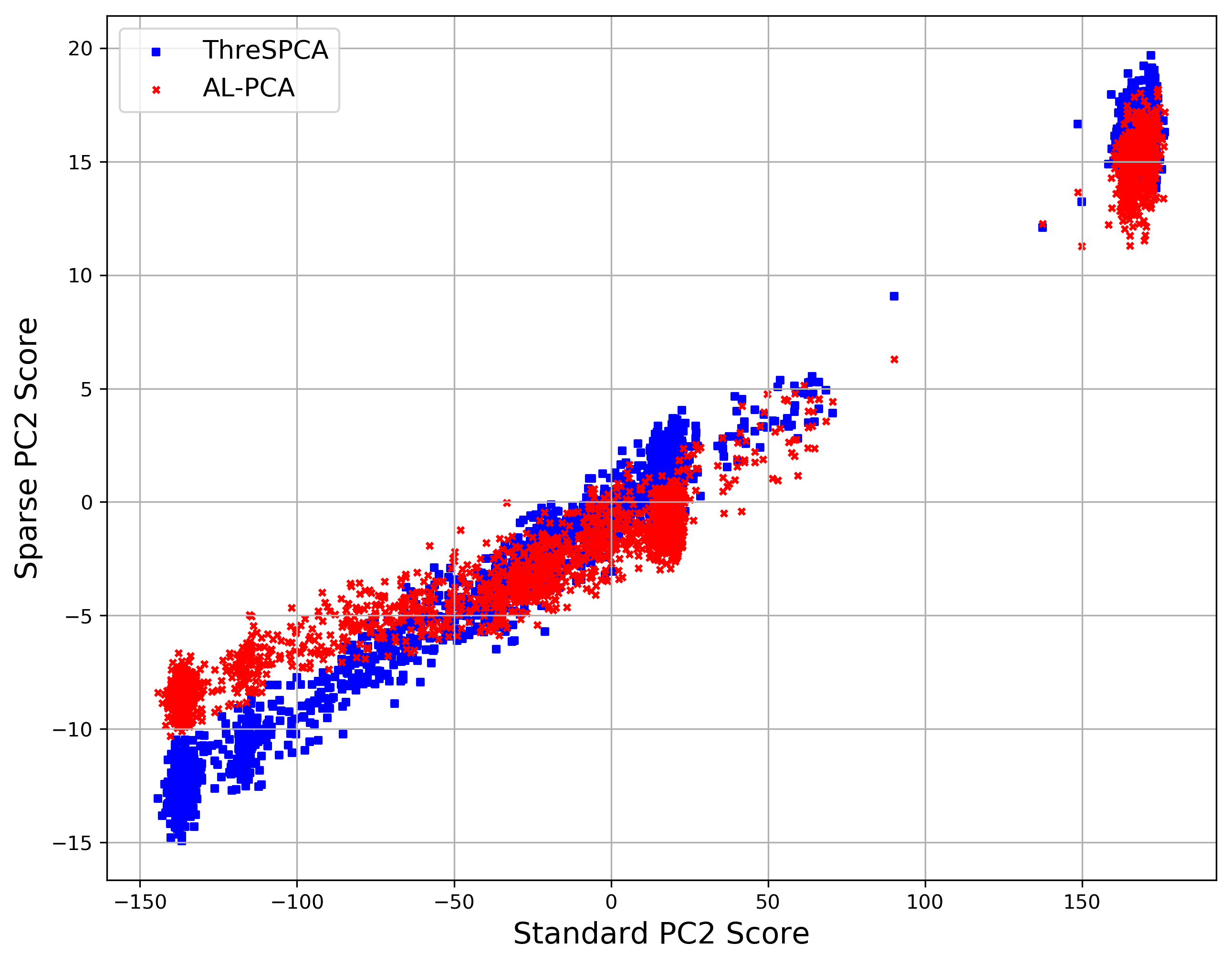

### PCcompare_1KG.png

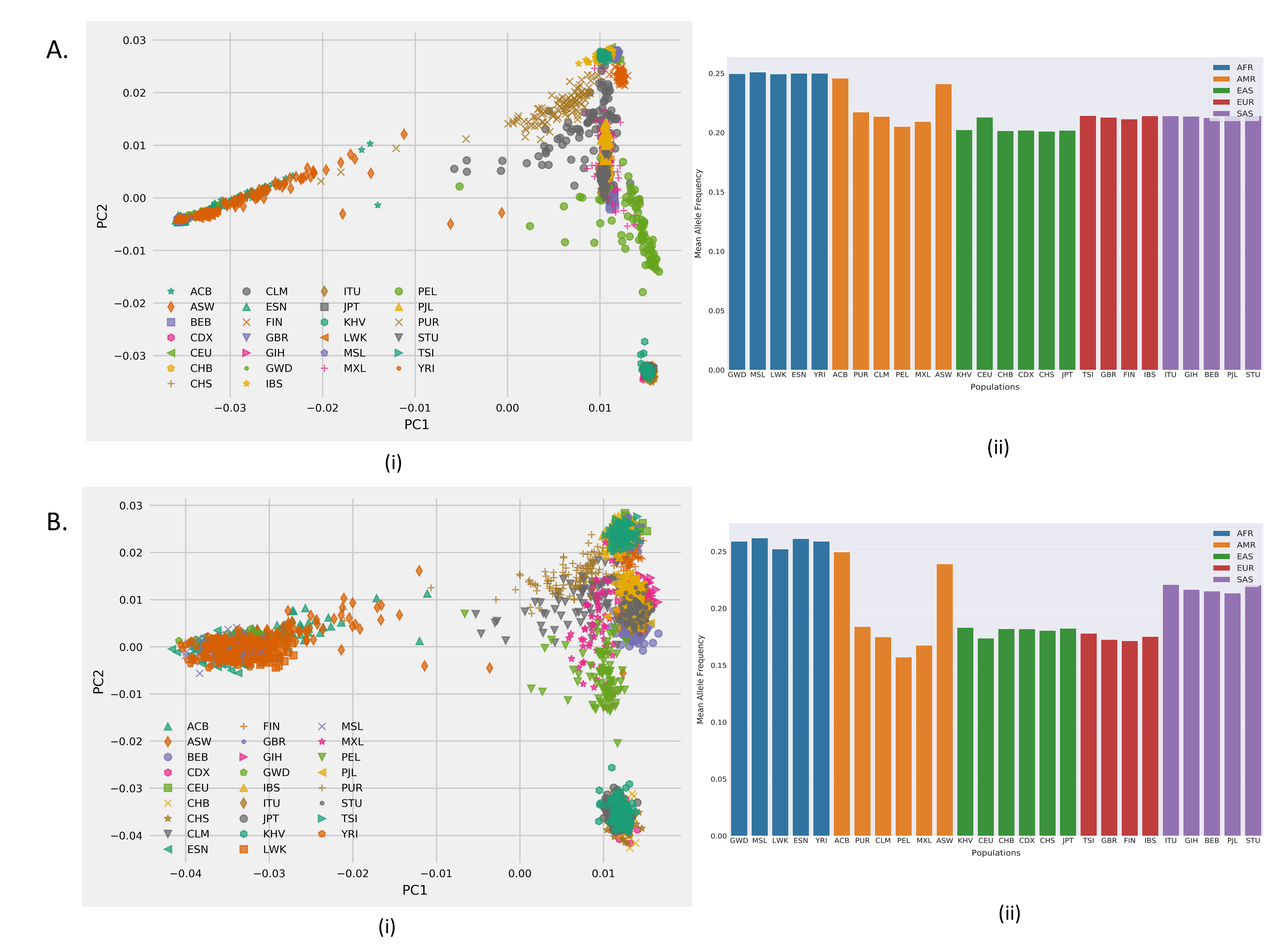

### PSD0.01_m10K_n1K_t100_PCs.png

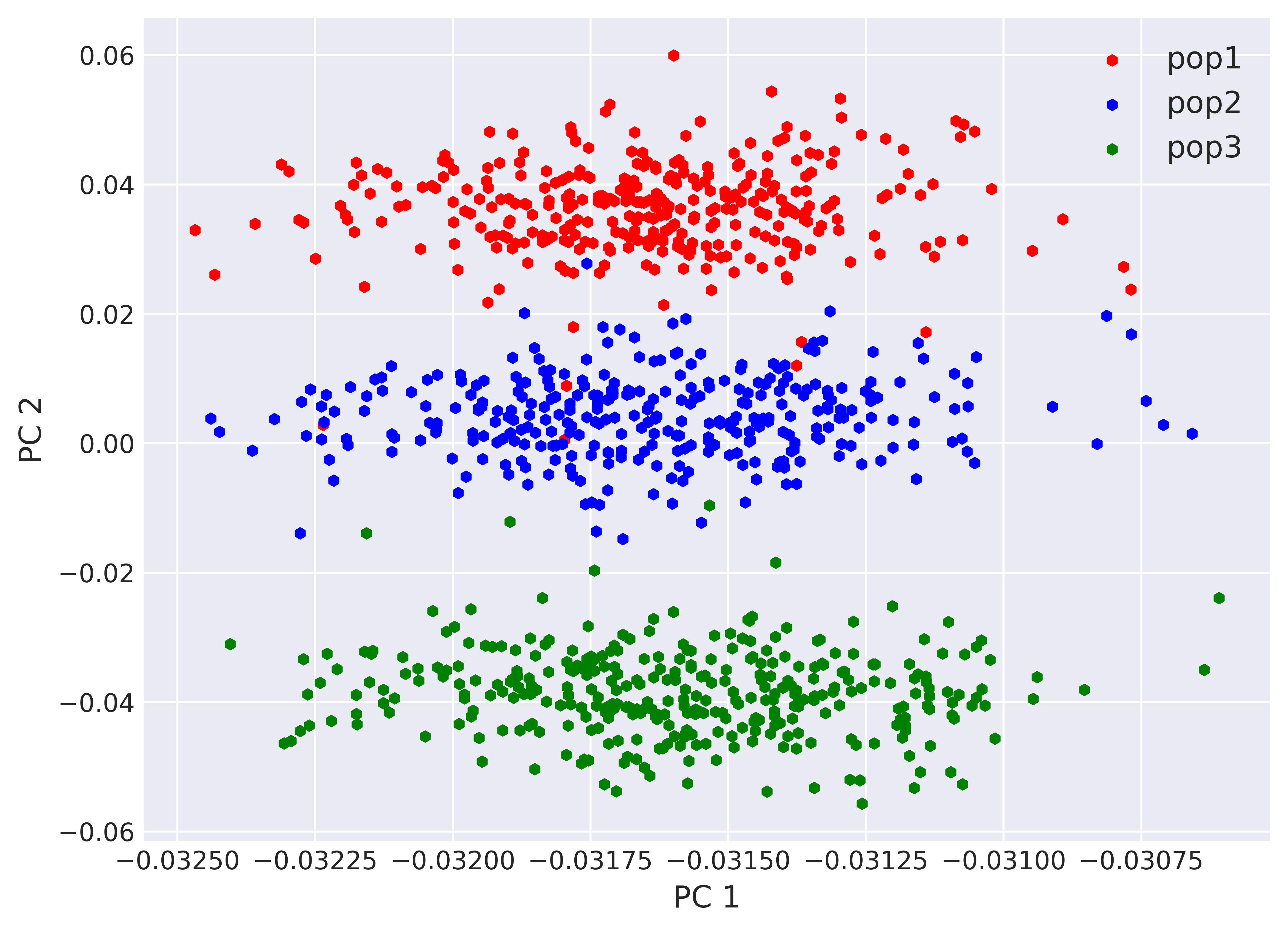

### PSD0.01_PCs.png

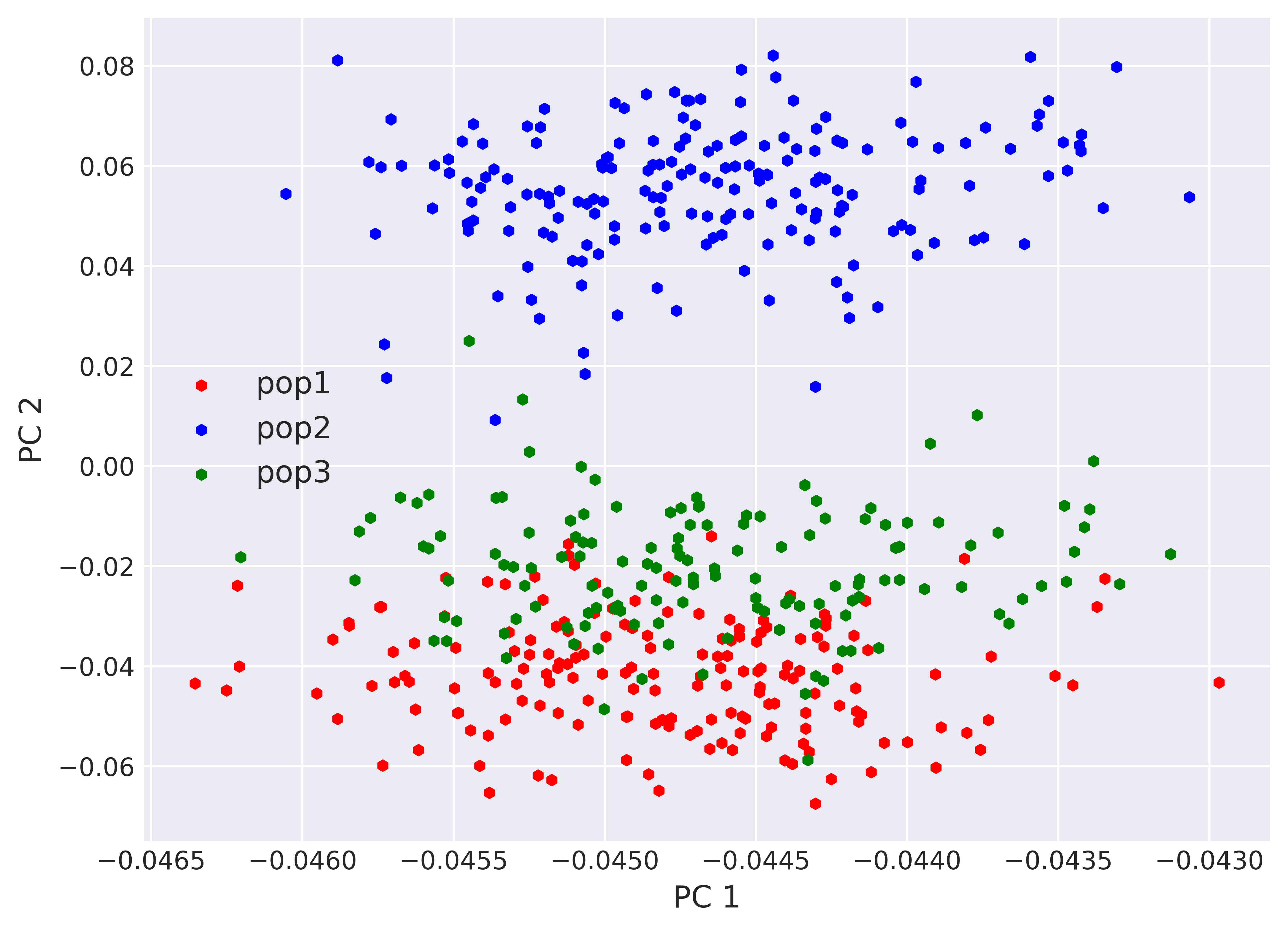

### SPC2_vs_SPC1_k500.png

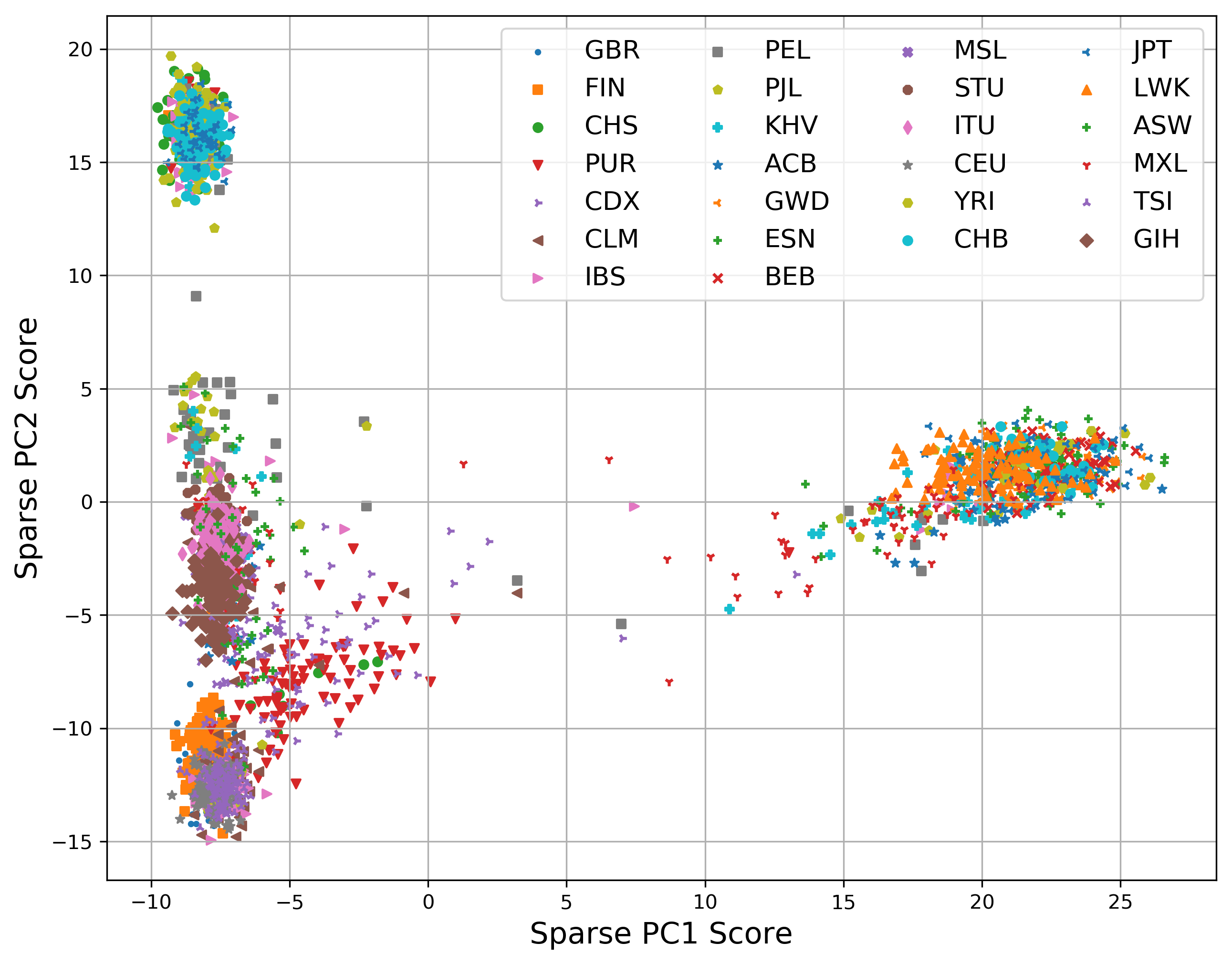

### SPC2_vs_SPC1_k1000.png

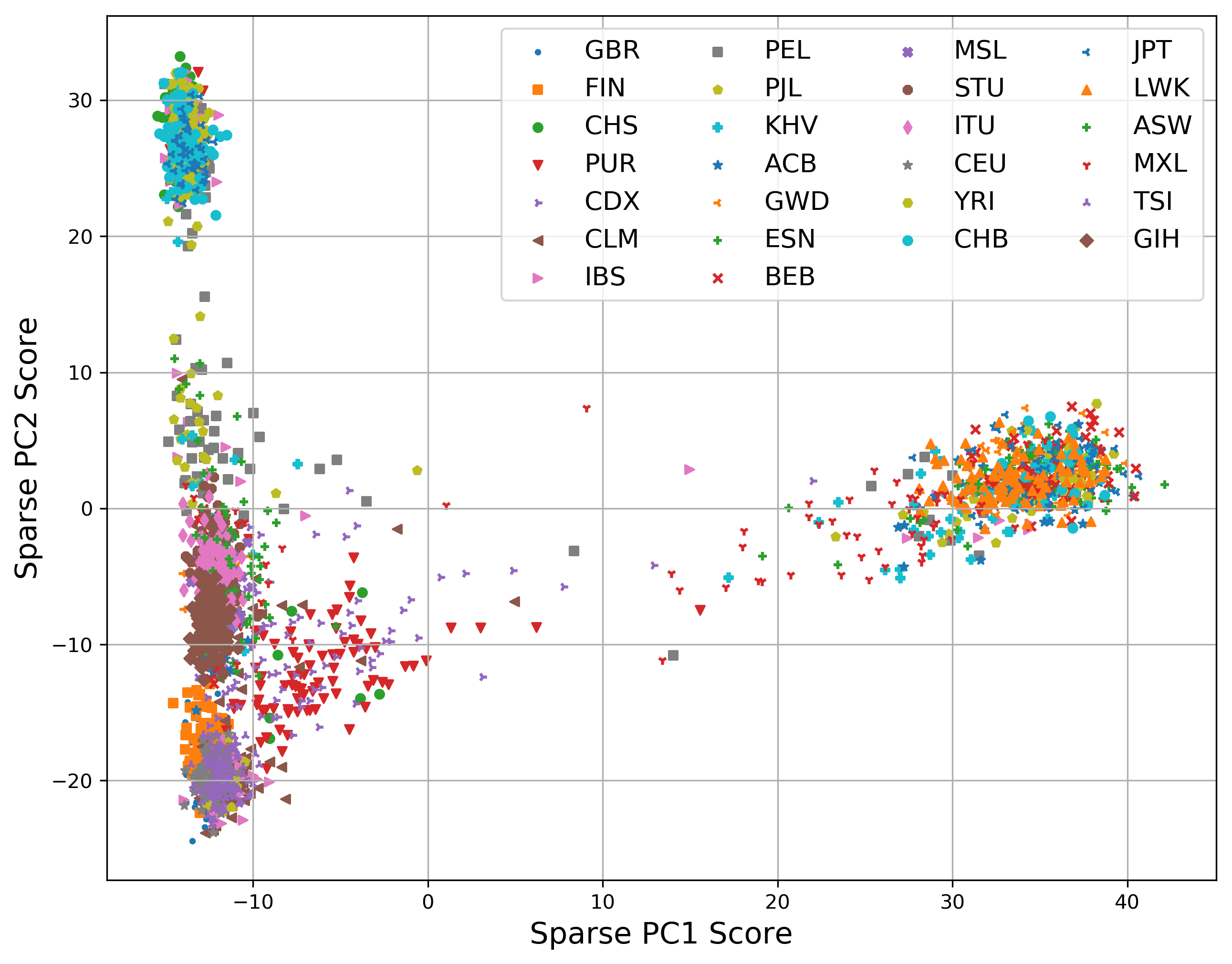

### SPC2_vs_SPC1_k5000.png

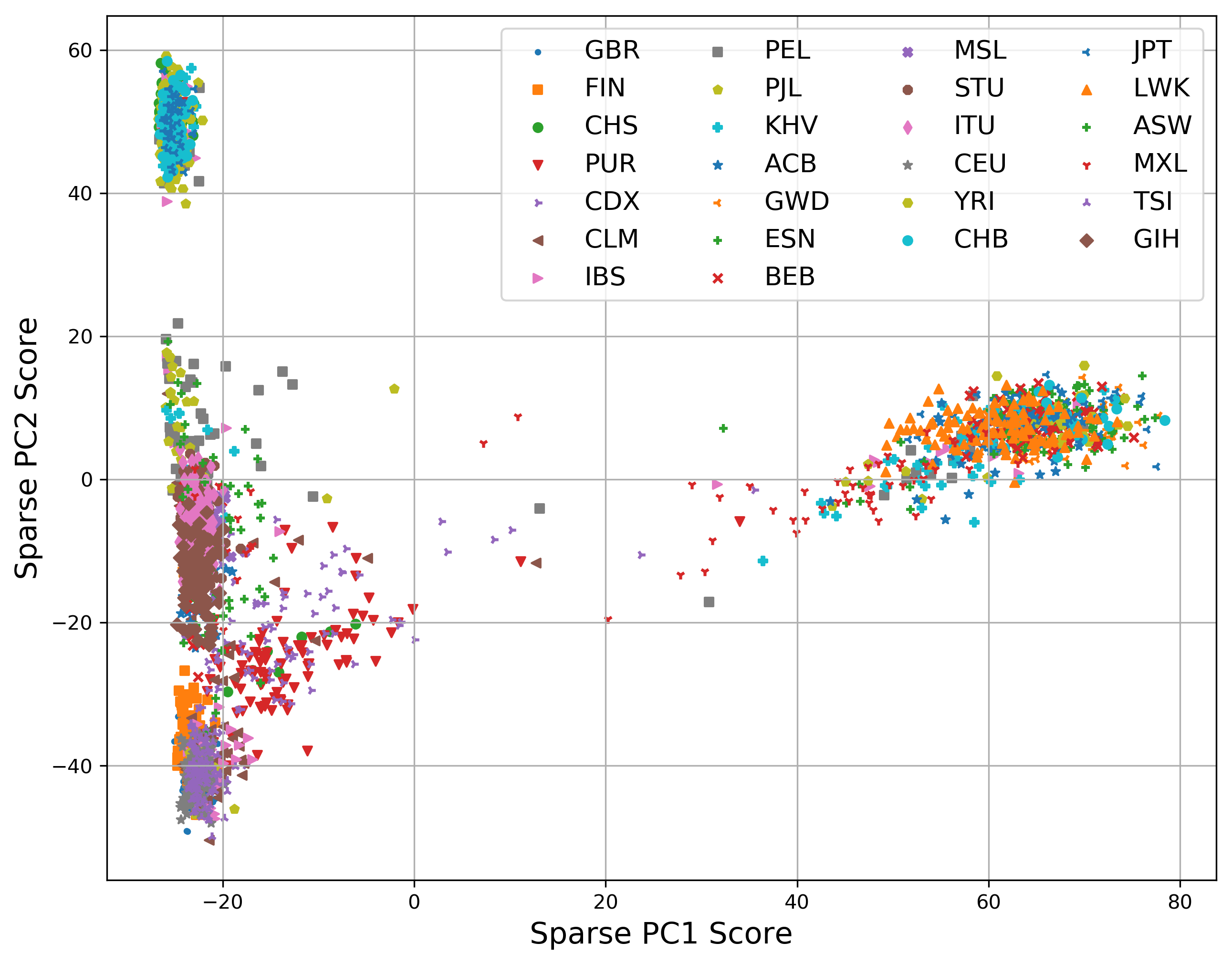

### SPC2_vs_SPC1_lee_k500.png

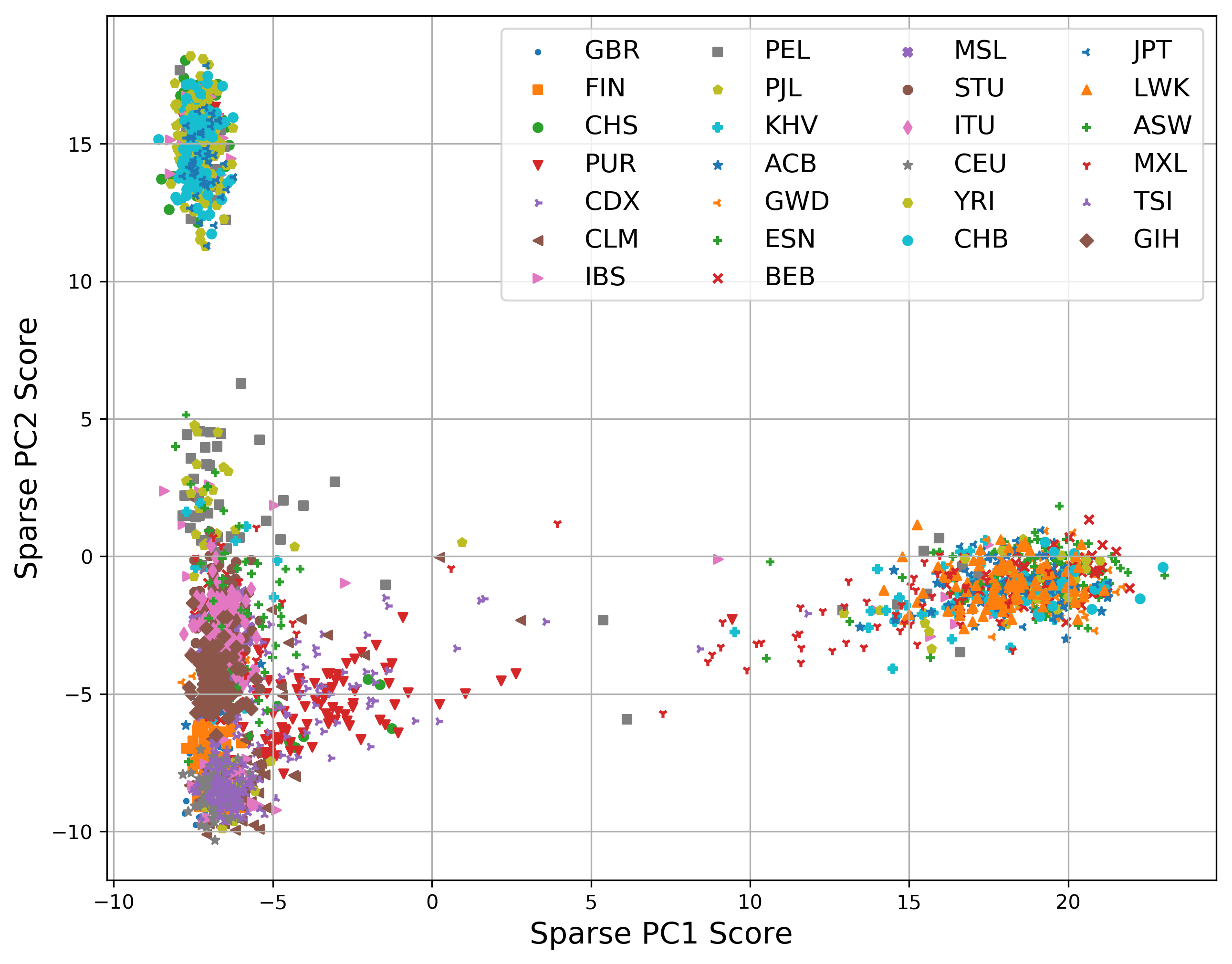

### tardis.png

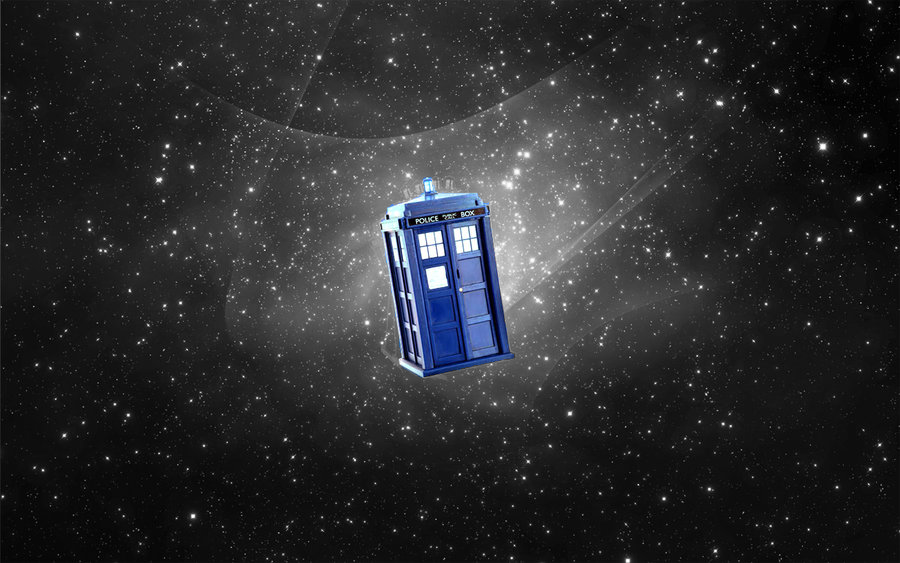

### ThreSPCA_PC1_k500.png

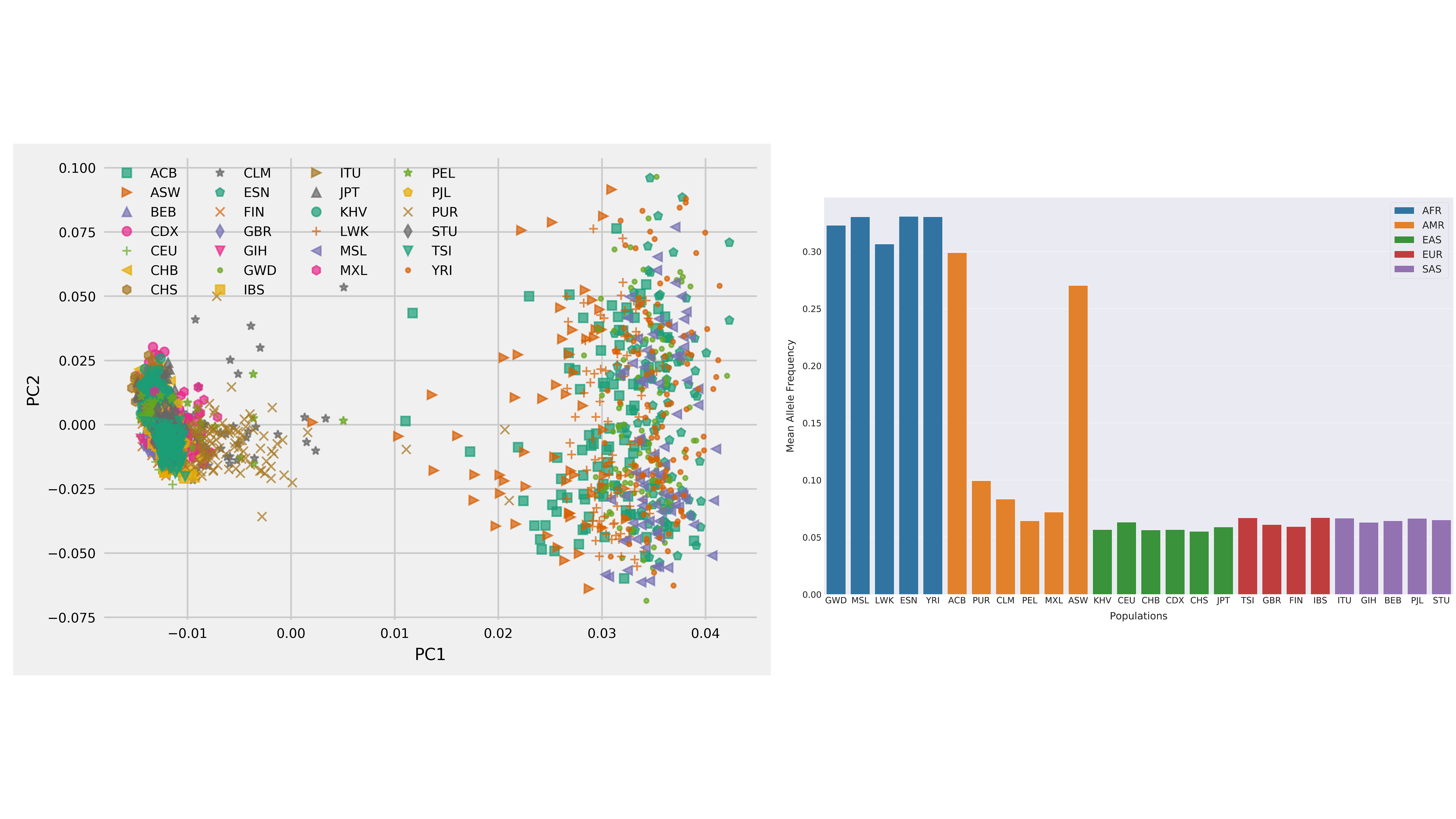

### ThreSPCA_PC2_k500.png

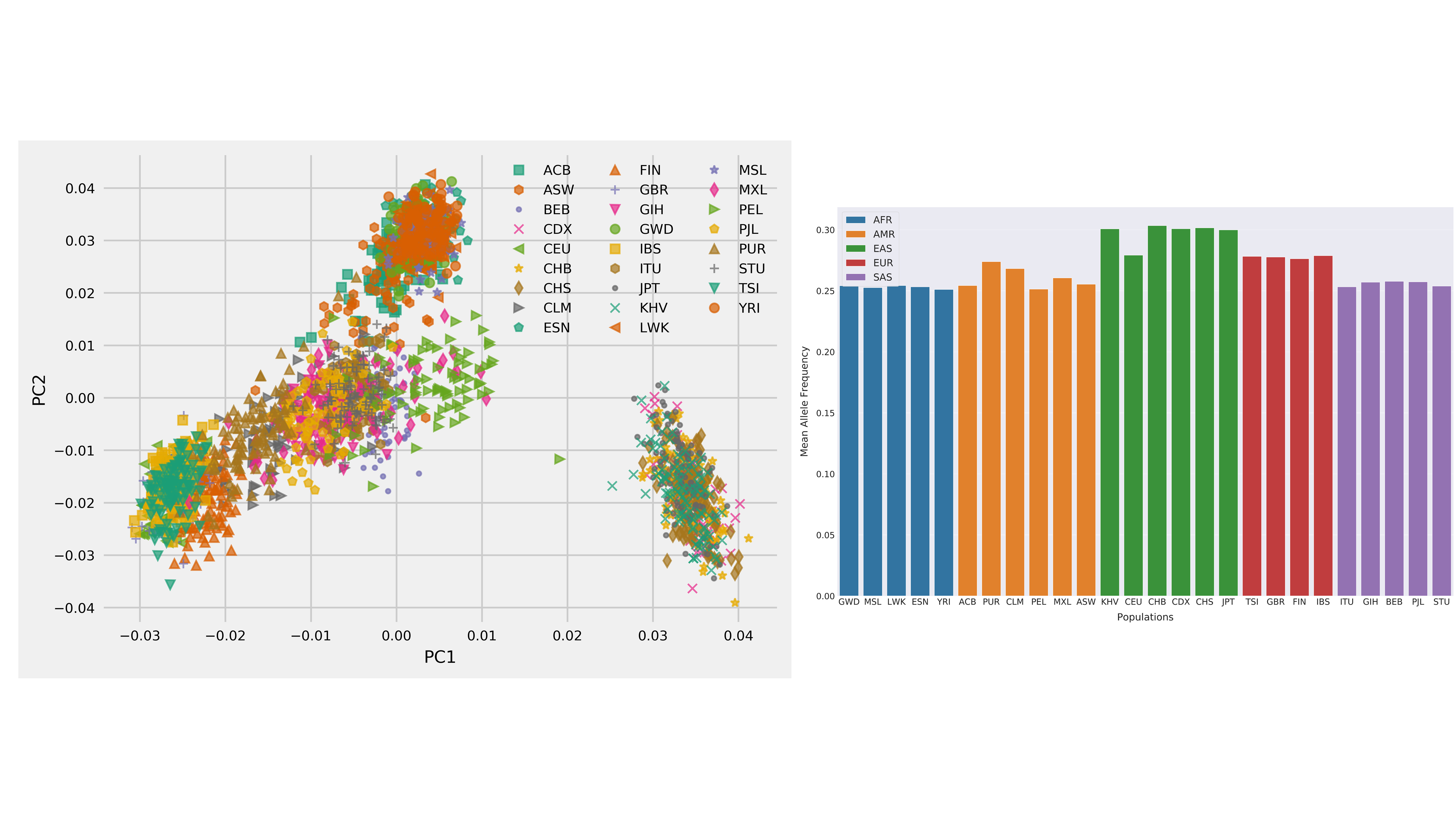

### ThreSPCA_PC3_k500.png

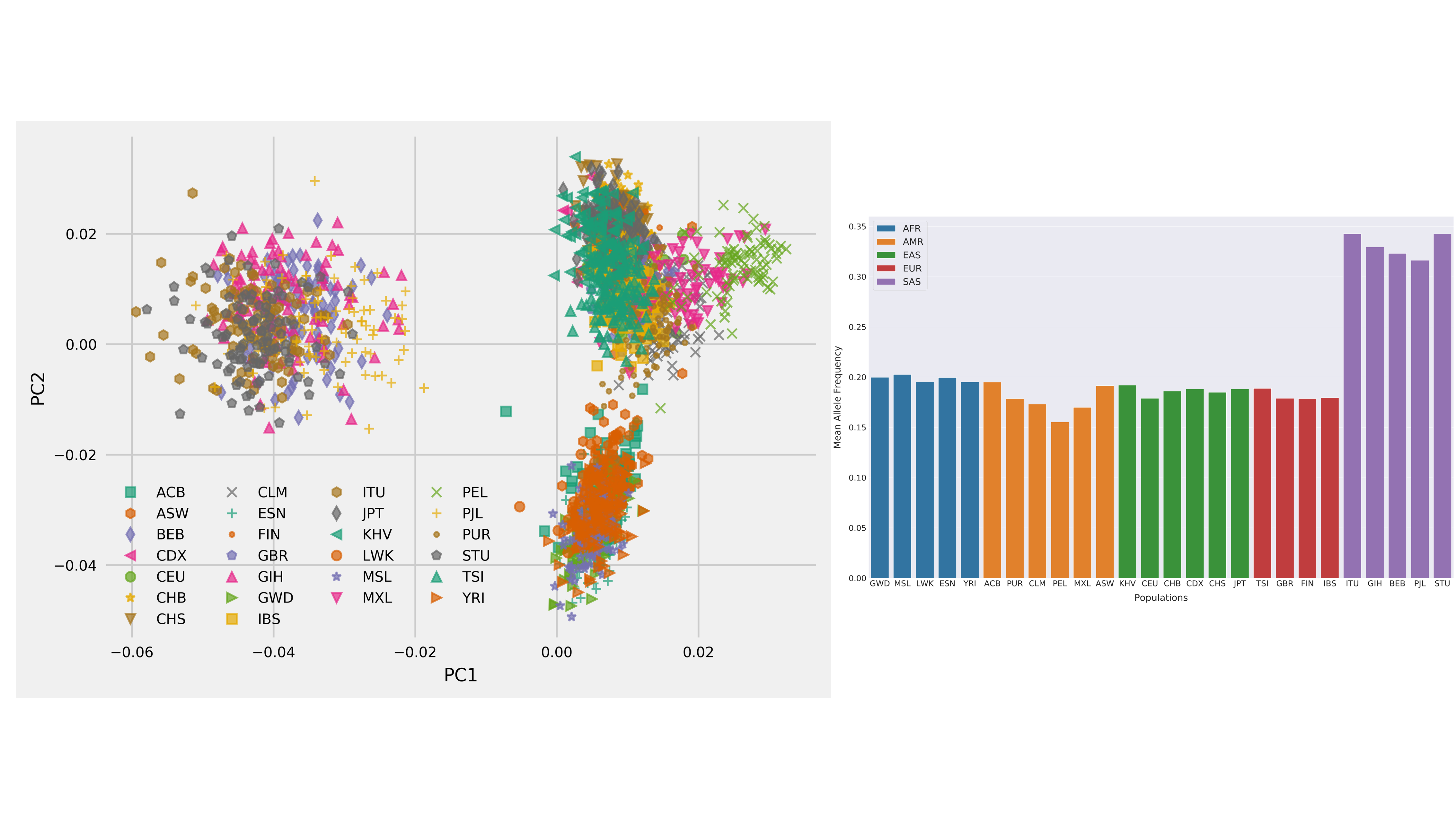

### TP_PSD_Fst_1kby10k_boxplot.png

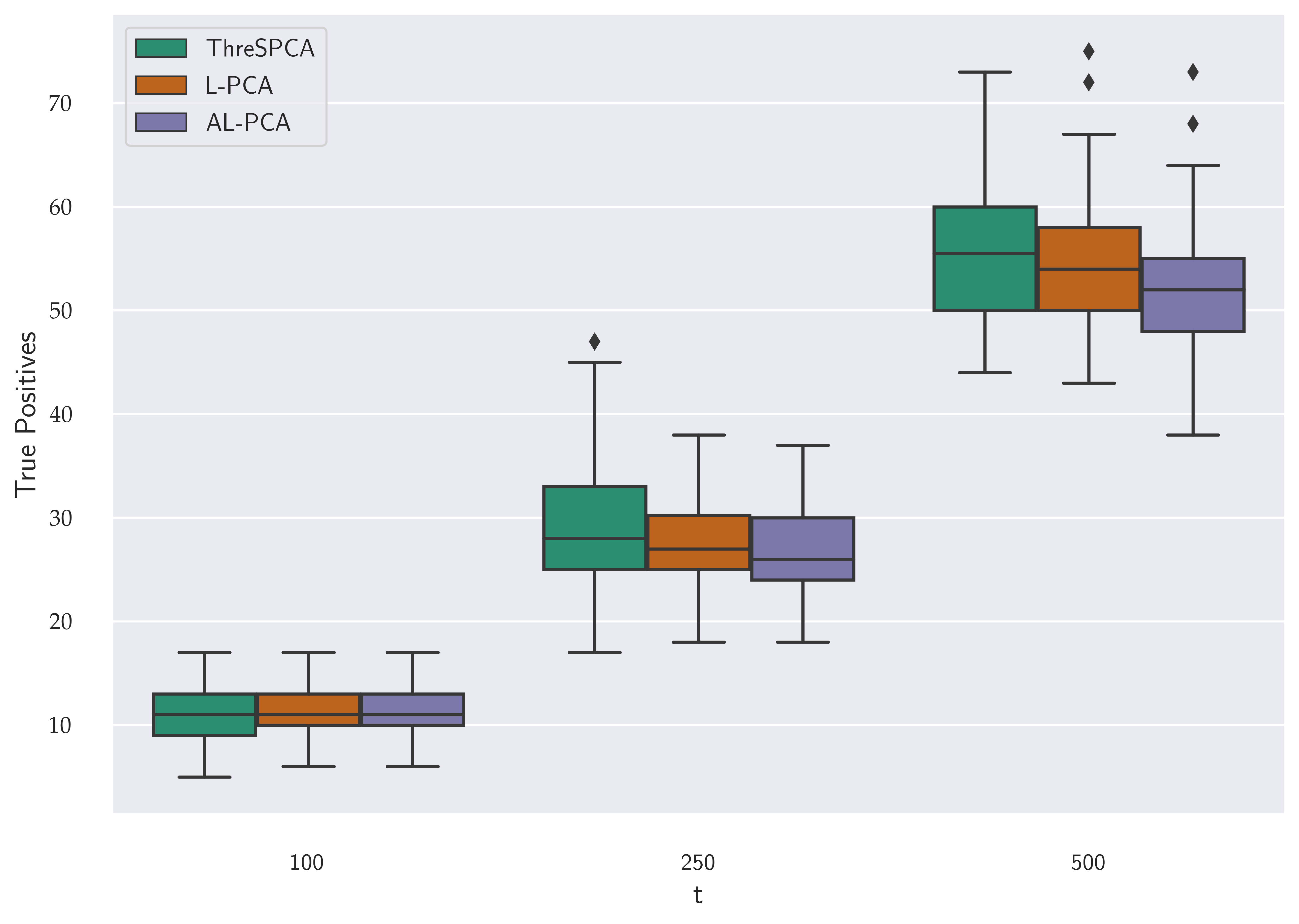

### TP_PSD_Fst_500by5k_boxplot.png

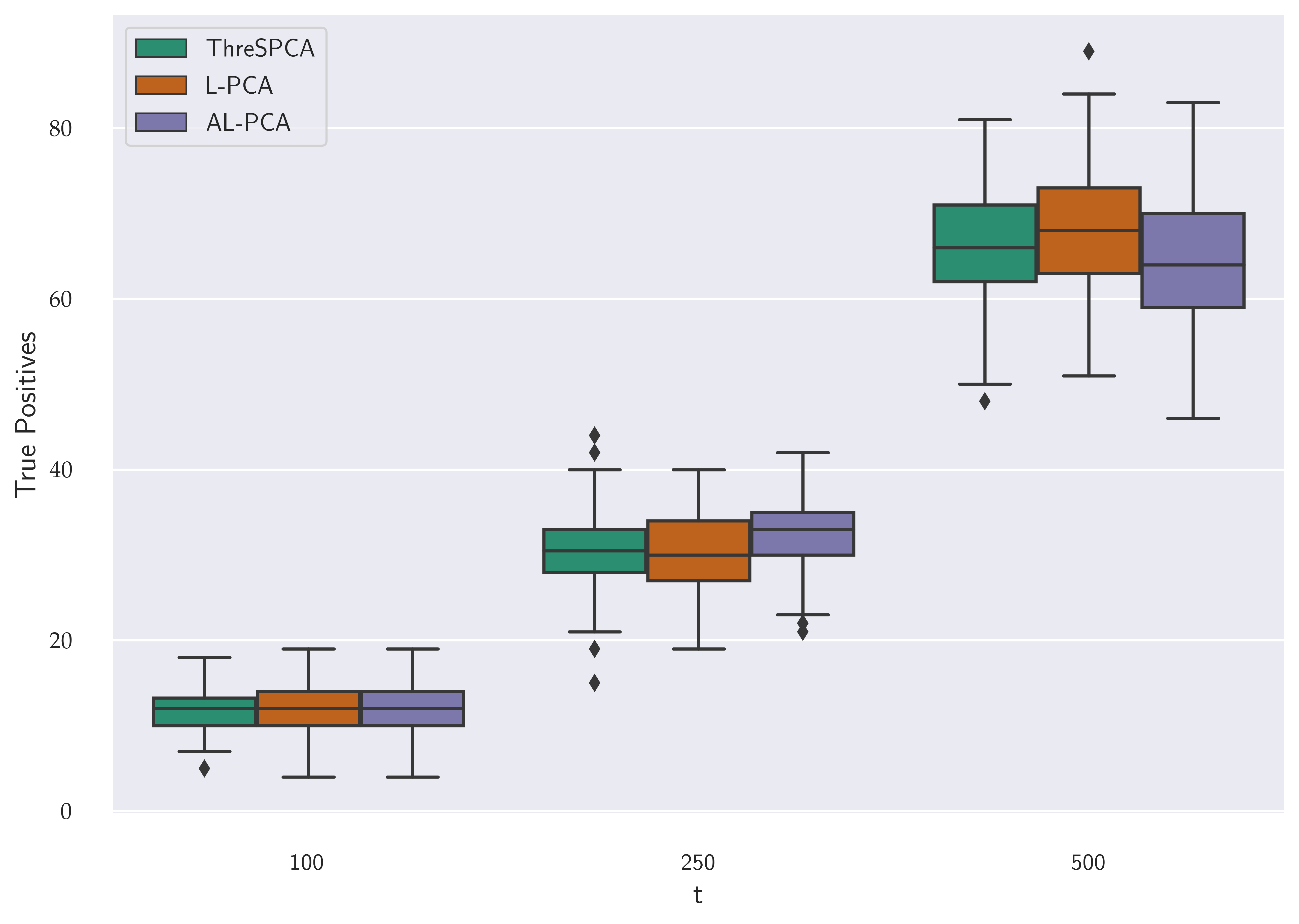

### vep_out.png

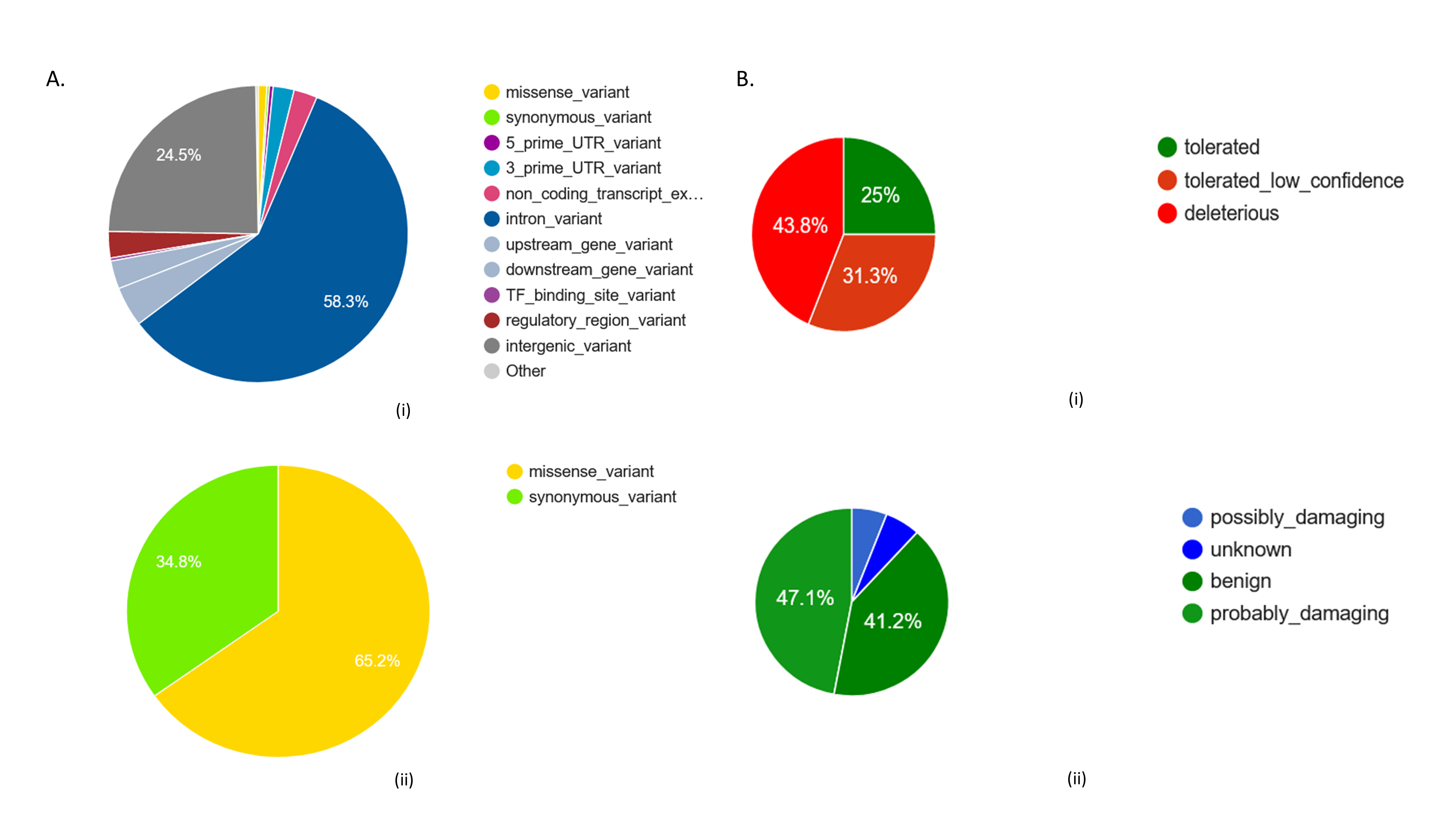
